# Determination of complete chromosomal haplotypes by bulk DNA sequencing

**DOI:** 10.1101/629337

**Authors:** Richard W. Tourdot, Cheng-Zhong Zhang

## Abstract

Haplotype phase represents the collective genetic variation between homologous chromosomes and is an essential feature of non-haploid genomes. Determining the haplotype phase requires knowledge of both the genotypes at variant sites and their linkage across each chromosome. Haplotype linkage can be either inferred statistically from a genotyped population, or determined by long-range sequencing of an individual genome. However, extending haplotype inference to the whole-chromosome scale remains challenging and usually requires special experimental techniques. Here we describe a general computational strategy to determine complete chromosomal haplotypes using a combination of bulk long-range sequencing and Hi-C sequencing. We demonstrate that this strategy can resolve the haplotypes of parental chromosomes in diploid human genomes at high precision (99%) and completeness (98%), and is further able to assemble the syntenic organization of aneuploid genomes (“digital karyotype”).

---

Haplotype phase (“haploid genotype”) is the combination of genotypes at sites of genetic variation along a chromosome (The International HapMap Consortium, 2005). Knowledge of chromosomal haplotypes is required for interrogating both the differences between homologous chromosomes, such as variations in the DNA sequence and in epigenetic features, and their impacts on gene expression (Shendure and Aiden, 2012; Adey et al., 2013a; Hansen and van Oudenaarden, 2013; Levesque and Raj, 2013; Deng et al., 2014; Snyder et al., 2015). Haplotype information is also needed for performing diploid genome assembly (Pendleton et al., 2015; Seo et al., 2016) and can significantly improve the precision and resolution of mutation detection in polyclonal populations (Nik-Zainal et al., 2012; Loh et al., 2018) or single cells (Zhang and Pellman, 2015).

There are two strategies of haplotype inference (Browning and Browning, 2011; Snyder et al., 2015). The first strategy (“statistical phasing”) (Scheet and Stephens, 2006; Browning and Browning, 2007; Loh et al., 2016a) infers haplotype phase based on the recombination probabilities between variant genotypes estimated from linkage disequilibrium in a population (Reich et al., 2001; Daly et al., 2001). Although statistical phasing can infer haplotype linkage between adjacent variant sites at reasonably high accuracy (99%), the accumulation of random switching errors (0.1%) precludes long-range (10Mb) haplotype inference unless the genotypes of closely related individuals are used (Kong et al., 2008; Loh et al., 2016b). Statistical phasing is also restricted to common polymorphisms and not applicable to *de novo* mutations.

The second strategy directly extracts haplotype linkage from the DNA sequences of single chromosomes or sub-haploid chromosomal fragments (“molecular linkage”) (Snyder et al., 2015). Direct sequencing of single chromosomes can produce whole-chromosome haplotypes (Ma et al., 2010; Fan et al., 2010; Yang et al., 2010; Zhang et al., 2015; Porubsky et al., 2017) but is only applicable to dividing cells and requires laborious experimental procedures of chromosome isolation or tagging. Long-read sequencing or long-range sequencing can either reveal haplotype linkage directly from long contiguous reads (~10kb), or indirectly from short DNA fragments derived from long DNA molecules (~100kb) that are tagged with unique molecular barcodes (Kitzman et al., 2010; Suk et al., 2011; Kaper et al., 2013; Adey et al., 2013b, 2014; Peters et al., 2014; Zheng et al., 2016; Chu et al., 2017; Marks et al., 2019). The typical size of DNA molecules in long-read or long-range sequencing (10-100kb) is sufficient for linking variants in regions of normal variant density (~1 per kb), but inadequate in regions of low variant density (1 per 10 kb), and unable to bridge large gaps (100kb) with no identifiable variants, including all centromeres.

Molecular linkage across individual chromosomes (*cis* linkage) is also contained in Hi-C contacts generated by proximity-based chromatin ligation (Rao et al., 2014). As chromosomes are spatially isolated in separate territories in the cell nucleus, Hi-C fragments are predominantly formed within the same chromosome and Hi-C sequencing can preserve intra-chromosomal linkage extending to the whole chromosome (Selvaraj et al., 2013) without single-chromosome isolation. The main shortcoming of haplotype linkage from Hi-C data is its sparsity: Only a small fraction of genetic variants are covered and linked by long-range Hi-C contacts; the sparse linkage cannot produce contiguous haplotypes (Edge et al., 2017) except for genomes with an exceptionally high variant density (~1 per 150 bp) (Selvaraj et al., 2013). The sparsity of long-range Hi-C linkage also limits its utility for phasing *de novo* mutations.

Although Hi-C contacts are too sparse to link single genetic variants, they generate sufficient linkage to concatenate haplotype blocks each consisting of hundreds of variants. This idea has led us to design a two-tier approach to determine complete whole-chromosome haplotypes combining long-range sequencing and Hi-C sequencing (Figure 1). This approach first determines local haplotype blocks consisting of 10^2^-10^3^ variants using long-range sequencing and then merge these blocks into a single haplotype using Hi-C contacts between blocks. We formulate both local haplotype inference and haplotype block concatenation as a minimization problem that can be efficiently solved by steepest descent methods. Applying our approach to two diploid human samples with reference haplotype data, we demonstrate that the computational inference produces the haplotypes of parental chromosomes with high accuracy (99%) and completeness (98%) and having no long-range switching error. We then describe strategies to analyze structural alterations of chromosomes using the parental haplotype, including the assembly of derivative chromosomes in aneuploid genomes using chromosome-specific Hi-C contacts.

**Figure 1:**
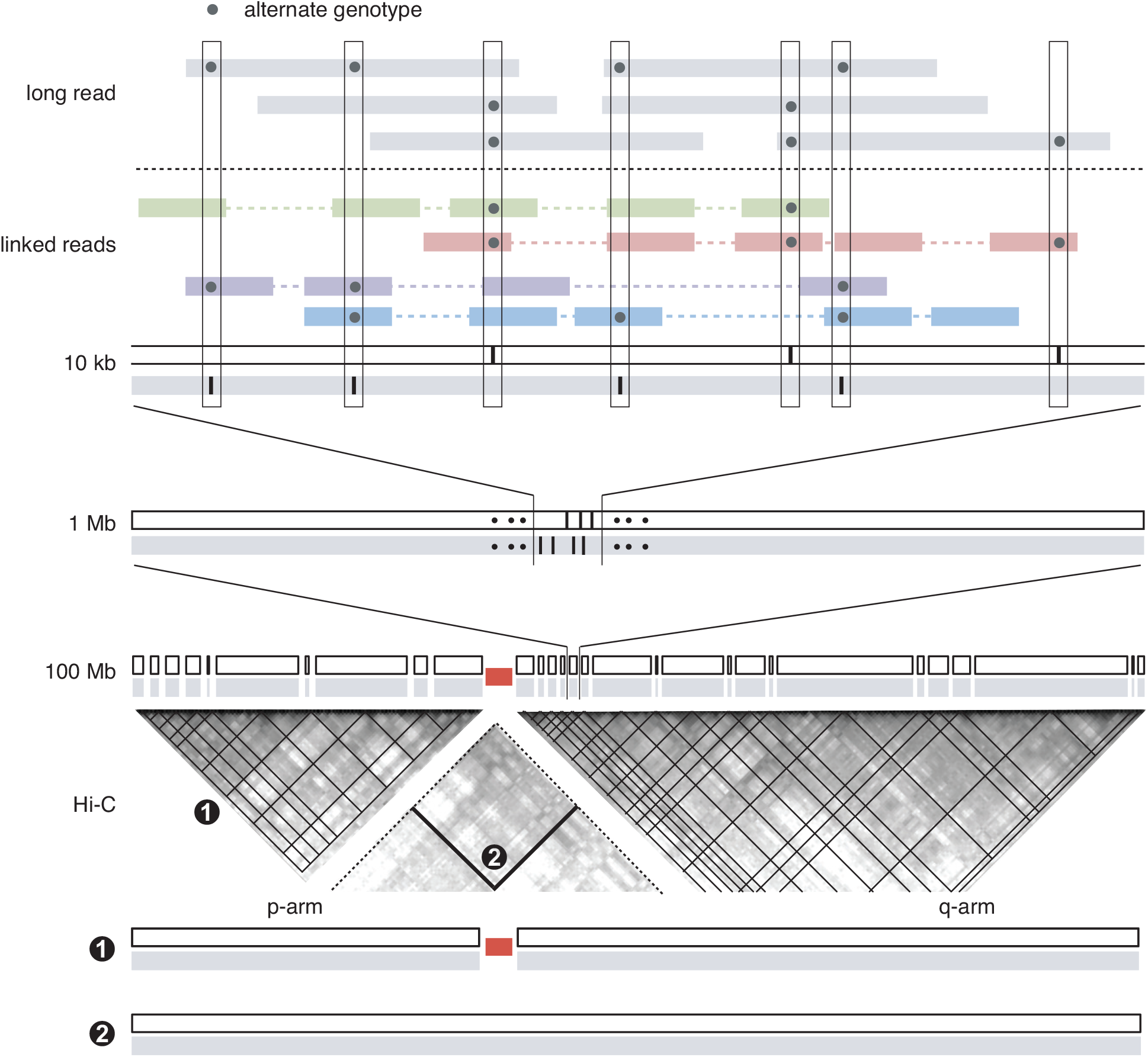
A hierarchical strategy for determining whole chromosome haplotypes. Megabase-scale haplotype blocks are determined using linkage information from linked-reads or long-read sequencing and then concatenated using Hi-C contacts, first within each chromosome arm and then between the p- and q-arms.

## Results

### Data generation and processing

For the computational inference and benchmarking of parental haplotypes in diploid genomes, we used published and newly generated datasets of the NA12878 cell line and the retinal pigment epithelium (RPE-1) cell line. Data sources are listed in Table 1. For the analysis of altered chromosomes in aneuploid genomes, we used bulk whole-genome sequencing data of aneuploid RPE-1 cells from Ref. (Maciejowski et al., 2015) and published cytogenetic (Gribble et al., 2000; Naumann et al., 2001) and sequencing (linked-reads and Hi-C) data of K-562 cells (Table S1). A detailed description of the experimental and data-processing protocols is provided in SI Materials and Methods.

**Table 1:** Sources of data for parental haplotype inference and benchmarking.

| Sample | Data type | Data source | Read count | Mean depth | Contacts (1 Mb) | Application |
| --- | --- | --- | --- | --- | --- | --- |
| RPE-1 | bulk WGS | Zhang et al. (2015) | 228,708,769<sup>a</sup> | 13× |  | variant calling |
| RPE-1 | linked reads | new | 941,518,426<sup>b</sup> | 60× |  | variant calling & local phasing |
| RPE-1 | Hi-C | Darrow et al. (2016) | 281,285,484<sup>d</sup> |  | 48,124,211 | long-range phasing |
| RPE-1 | single cell with monosomies | new |  |  |  | hi-conf variants & reference haplotypes |
| NA12878 | linked reads v.1 | 10X Genomics<sup>e</sup> | 422,179,395<sup>f</sup> | 35× |  | local phasing |
| NA12878 | linked reads v.2 | 10X Genomics<sup>g</sup> | 423,854,243<sup>h</sup> | 35× |  | local phasing |
| NA12878 | Hi-C | Rao et al. (2014) | 486,848,169<sup>i</sup> |  | 91,428,507 | long-range phasing |
| NA12878 | phased VCF | GIAB<sup>j</sup> |  |  |  | hi-conf variants & reference haplotypes |

<sup>a</sup>SRR1778442: median insert 243; 208,151,992 aligned in pair; 101 × 2; duplication rate 0.024.
<sup>b</sup>Mean molecular length 24.8 kb; median insert 551; 913,660,083 aligned in pair; 150 × 2; duplication rate 0.255.
<sup>c</sup>excluding the GEMcode sequence
<sup>d</sup>SRS1045722: median insert 364; 279,027,892 aligned in pair; 150 × 2; duplication rate 0.067.
<sup>e</sup>https://support.10xgenomics.com/genome-exome/datasets/2.1.0/NA12878_WGS_210
<sup>f</sup>Mean molecular length 68.7 kb; median insert 349; 407,015,530 aligned in pair; 150 × 2; duplication rate 0.062.
<sup>g</sup>https://support.10xgenomics.com/genome-exome/datasets/2.2.1/NA12878_WGS_v2
<sup>h</sup>Mean molecular length 85.6 kb; median insert 370; 418,283,435 aligned in pair; 150 × 2; duplication rate 0.079.
<sup>i</sup>SRR1658572: median insert 377; 484,211,662 aligned in pair; 101 × 2; duplication rate 0.028.
<sup>j</sup>https://ftp-trace.ncbi.nlm.nih.gov/ReferenceSamples/giab/release/NA12878_HG001/latest/GRCh38/

### Hi-C sequencing can reveal haplotype linkage across entire chromosomes

The basic unit of haplotype linkage evidence is a molecular link, which we use to denote a single DNA molecule providing genotype information at multiple variant sites. In long-read sequencing, a link consists of a single contiguous alignment; in Hi-C data, a link consists of multiple alignments of a Hi-C fragment; in linked-reads sequencing, a link consists of a group of sequencing fragments sharing the same molecular barcode. To demonstrate the utility of Hi-C sequencing for long-range haplotype inference, we calculate three metrics of haplotype linkage between heterozygous variant sites at different genomic distance: (1) percentage of linked variants; (2) average number of links between linked variants; and (3) percentage of links consistent with *cis* linkage according to the reference parental haplotype. These results are shown in Fig. 2 together with the same metrics of haplotype linkage from linked-reads data for comparison.

**Figure 2:**
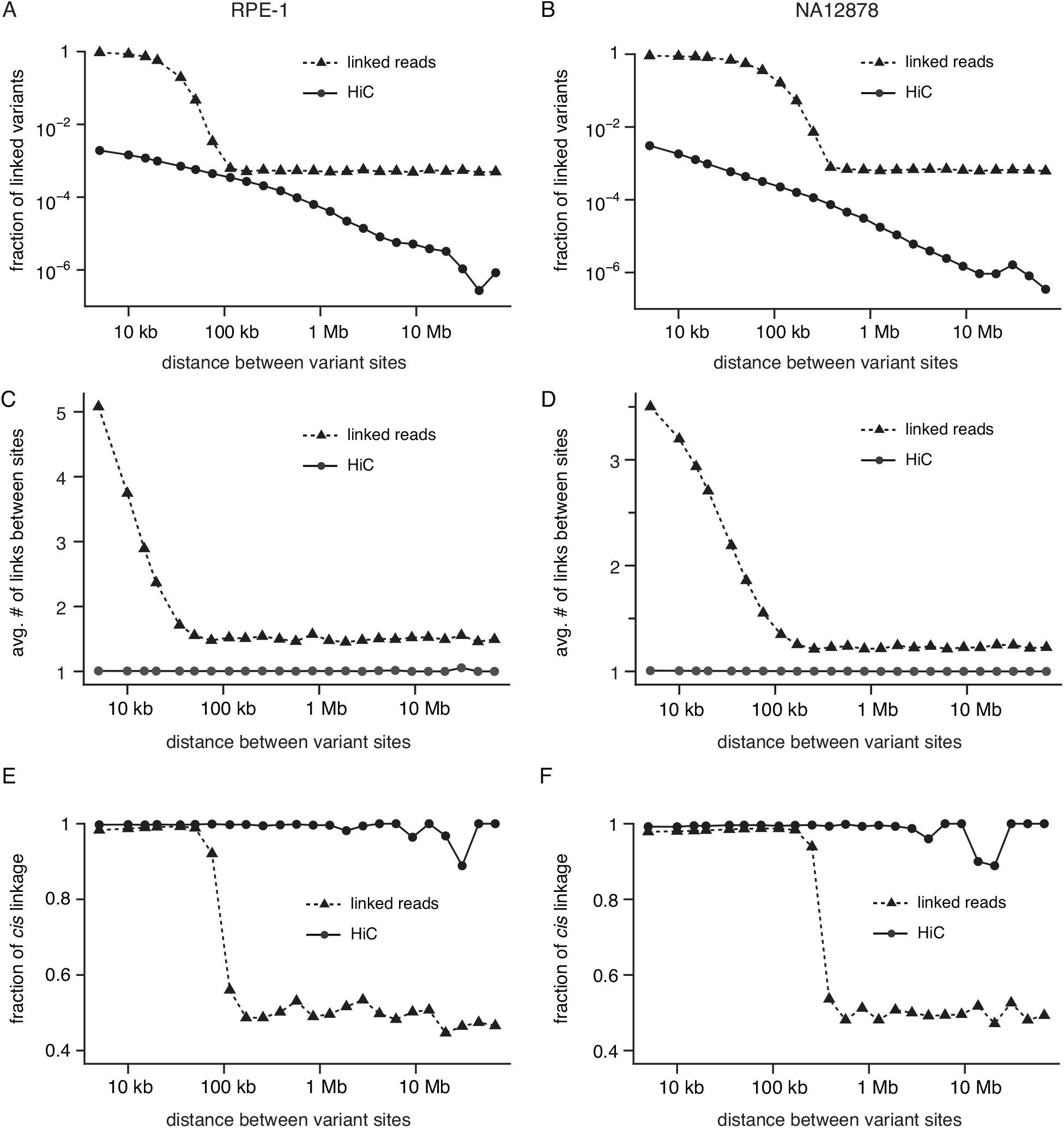
Statistical metrics of haplotype linkage between variants at different genomic distance extracted from linked-reads and Hi-C sequencing data of RPE-1 (A,C,E) and NA12878 (B,D,F) cells. A and B. Fraction of linked variants; C and D. Average number of links between linked variants; E and F. Fraction of links consistent with *cis* linkage. The residual *cis* linkage in linked-reads data (~50%) reflects unrelated DNA fragments from either parental homologs being tagged by identical molecular barcodes by chance.

The density of Hi-C links shows a power-law decay against genomic distance (Fig. 2A,B) that is similar to the probability of intra-chromosomal contacts (Lieberman-Aiden et al., 2009; Sanborn et al., 2015). The power-law dependence suggests that most Hi-C links result from intra-chromosomal contacts, as the presence of inter-chromosomal contacts (which should form largely at random) will cause deviations from the power-law decay. This is validated by the observation that more than 90% of all Hi-C links are consistent with *cis* contacts between loci in the same chromosome (Fig. 2E,F). By contrast, haplotype linkage from the linked-reads data has a limited range determined by the maximum size of input DNA molecules (~100kb in the RPE-1 library and ~300kb in the NA12878 libraries). The distance-independent linkage signal beyond the molecular size most likely arises from unrelated DNA fragments being tagged by the same molecular barcode by chance: As the unrelated DNA fragments can originate from either parental chromosomes, the average percentage of apparent *cis* or *trans* linkage is 50% (Fig. 2E,F), despite all links being intermolecular.

Although Hi-C links across the entire chromosome are dominated by intra-molecular Hi-C contacts, the sparsity of long-range Hi-C contacts limits its utility for haplotype inference. In both Hi-C data (RPE-1 and NA12878), the probability that two variant sites separated by 100kb are linked by Hi-C reads is less than 10^−3^ (Fig. 2A,B) and almost all linkage consists of only one link/read (Fig. 2C,D). Such weak linkage evidence is inadequate for accurate haplotype inference at all variant sites.

To take advantage of long-range Hi-C linkage without significantly increasing the number of Hi-C reads, we can aggregate Hi-C links between local haplotype blocks consisting of 10^2^ or more variants to generate a specific linkage signal between these blocks. To demonstrate this quantitatively, we calculate the average number of Hi-C links between 0.5, 1, and 2Mb segments at different genomic distance (Fig. 3A). This calculation shows that the Hi-C linkage signal between megabase-scale segments can extend well above 10Mb, which is sufficient to merge all haplotype blocks on each chromosome. As haplotype inference by either statistical phasing or long-range sequencing can conveniently generate megabase-scale haplotype blocks, we can use Hi-C sequencing only for linking these blocks across low-variant density regions, including gaps in the reference genome. We next describe a general computational strategy of haplotype inference that is applicable to both local haplotype inference and haplotype block concatenation (Fig. 3B).

**Figure 3:**
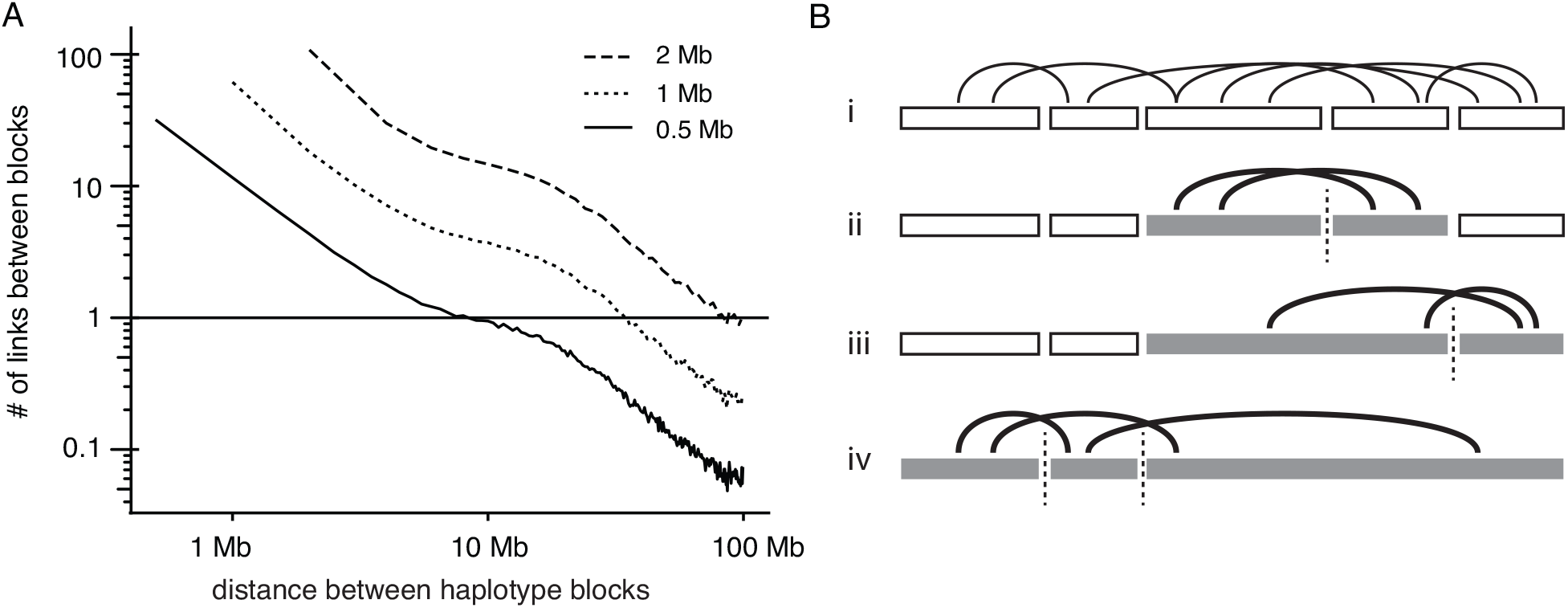
Linking haplotype blocks with Hi-C reads. A. Average number of Hi-C links between two segments of 0.5, 1, and 2 Mb at different genomic distance calculated using the RPE-1 Hi-C data. B. Concatenation of small haplotype blocks using Hi-C reads. (i) Local haplotype blocks (open rectangles) and Hi-C links (curves); (ii) and (iii) merging of adjacent haplotype blocks using Hi-C links; (iv) joint inference of the haplotype phase of all blocks using all Hi-C links.

### Haplotype inference from linkage evidence

We first introduce a binary numerical representation of genotypes at heterozygous variants as 1 for the reference base and 1 for the alternate base. A haplotype block consisting of *N* variant sites is represented as a vector

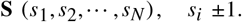

Similarly, a molecular link with genotype information at multiple variant sites is represented as

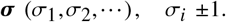

Using the binary genotype representation, we can simply four types of linkage between genotypes (for both σ_*i*_ and *s*_*i*_)

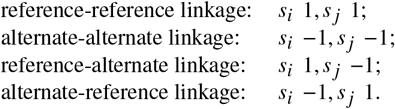

into two types of haplotype linkage

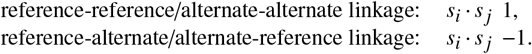

Moreover, a molecular link (σ_*i*_,*σ* _*j*_) between sites *i* and *j* is consistent with haplotype linkage (*s*_*i*_, *s*_*j*_) if and only if

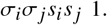

If the error probability of a molecular link is given by ϵ_*ij*_, then

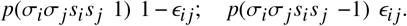

Assuming *p*(*s*_*i*_*s*_*j*_ 1) *p*(*s*_*i*_*s*_*j*_ −1) 1/2, we can re-write the above equation as

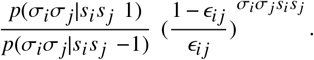

Extending this to a collection of links 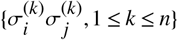, we have

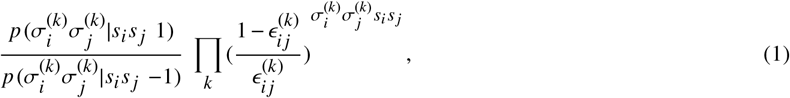

which leads to the following log-likelihood function

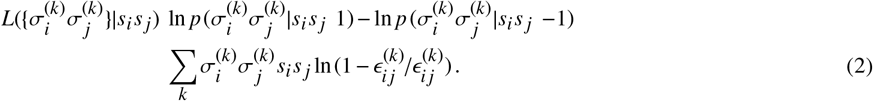

If we assume a constant error rate for all links, 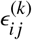, Eq. (2) is simplified to

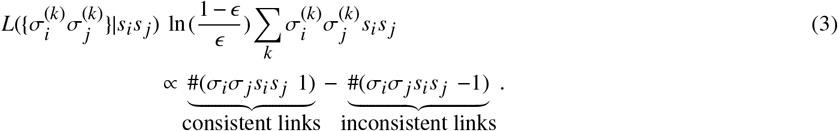

The haplotype linkage inferred from all molecular links is given by

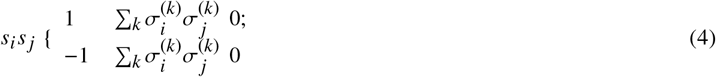

We can generalize Eq. (2) to *N* variants as

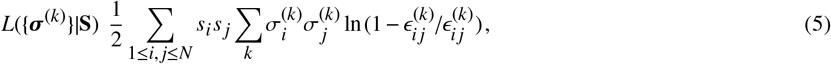

and solve for the optimal haplotype solution 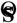 by maximizing Eq. (5) (assuming a uniform prior probability *p*(**S**)). We further assume the frequency of incorrect linkage is constant between different molecules but can vary between variant sites, 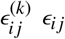. We can then rewrite Eq. (5) as

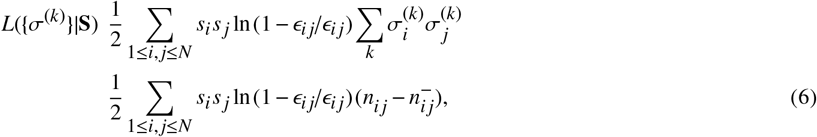

with the introduction of

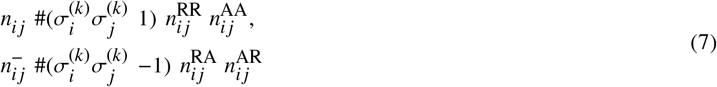

corresponding to the number of links supporting each type of haplotype linkage between site *i* and *j*. We estimate ϵ_*ij*_ using

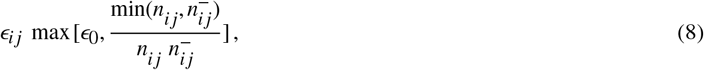

where 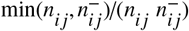 is the observed fraction of minor discordant haplotype linkage, and

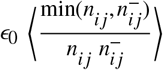

is the average fraction of observed discordant linkage.

The formalism of haplotype inference defined in Eqs. (5) and (6) has several advantages. First, the binary representation of haplotype phase and molecular linkage preserves the symmetry between parental haplotypes (**S** and **S**) or molecular links derived from parental chromosomes (***σ*** and −***σ***). This is convenient for performing haplotype inference in aneuploid genomes where one homolog may contribute dominant linkage evidence (*e.g.*, in hemizygous or trisomic regions).

Second, the formalism is directly applicable to haplotype block phasing. The phase of haplotype blocks **B**_1_, **B**_2_, …, **B**_*m*_ can be represented as a vector

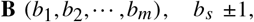

and determined from inter-block molecular linkage (similar to Eq. (7))

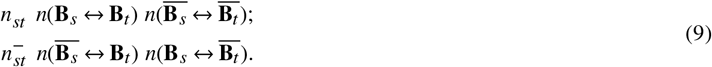

(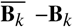 is the complementary phase block of **B**_*k*_.)

Finally, maximizing Eq. (6) is equivalent to minimizing the energy of a 1D Ising model (or spin glass model)

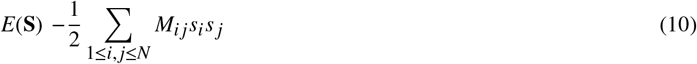

with finite range interactions 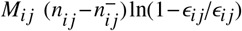, for which there are many existing approaches. Here we solve this problem by introducing two types of perturbations:

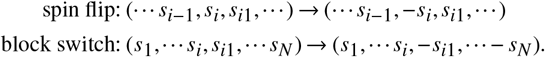

The changes to *E*(**S**) due to these perturbations are given by

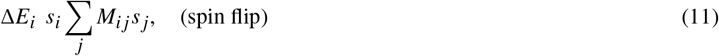

and

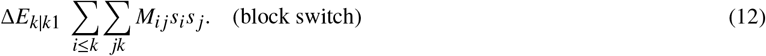

It can be shown that through iterations of spin flipping and block switching, one can always find (one of) the optimal haplotype solutions 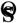 that minimizes Eq. (10) if the majority of molecular linkage is consistent with *cis* haplotype linkage (SI Discussion).

For two haplotype configurations **S** and **S**′, the energy difference is related to the likelihood ratio

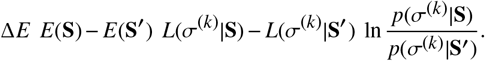

Thus, Δ*E*_*i*_ or Δ*E*_*i*|*i*1_ are related to the probability of phasing errors:

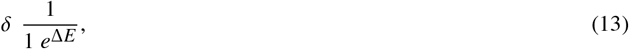

A low energy penalty score (Δ*E* ≈ 0) implies low phasing confidence (δ ≈ 0.5), and vice versa (Δ*E* » 0 ⇒ δ ≈ 0).

The spin-flipping energy penalty (Eq. (11)) can be rewritten as

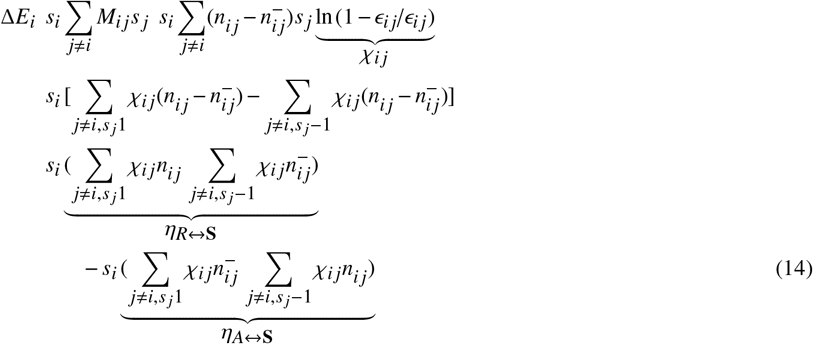

The two terms η_*R*↔**S**_ and η_*A*↔**S**_ in Eq. (14) measure the total linkage between the genotypes at site *i* (*R* for reference and *A* for alternate) and the haplotypes of parental chromosomes (**S** and −**S**). For true heterozygous variants, reference and alternate genotypes are phased to complementary haplotypes, *i.e.*, *R* ↔ **S** (and hence *A* ↔ −**S**), or *A* ↔ **S** (*R* ↔ −**S**). This implies that either η_*R*↔**S**_» η_*A*↔**S**_≈ 0 or η_*A*↔**S**_» η_*R*↔**S**_≈ 0. As the genotypes of false variants are generally not phased to complementary haplotypes, false variants tend to have low phasing confidence (Δ*E* ≈ 0) and can be excluded from the haplotype solution based on this feature. Moreover, linkage evidence from false variants is offset by the χ_*ij*_ factor due to the presence of significant discordant linkage evidence (ϵ_*ij*_ » ϵ_0_). These features make our haplotype inference method robust against the presence of ambiguous haplotype linkage from false variant sites and can implicitly exclude some of these variants based on haplotype linkage.

### Computational inference of the parental haplotypes of diploid NA12878 and RPE-1 genomes

We first performed local haplotype inference using the linked-reads data. We restricted haplotype inference to bi-allelic single-nucleotide variant (SNV) sites as they represent the majority of genetic variation and have better detection accuracy than more complex alterations (insertion/deletion/structural variants). For each chromosome, we accumulated all molecular linkage between SNVs within 100kb from each other and solved for the optimal haplotype by minimizing Eq. (10). The haplotype solution converged after 4-5 rounds of spin flipping and block switching. See SI Discussion and Data Table S1 for more technical details.

Phasing accuracy is assessed using the energy penalties calculated based on the final haplotype solution: The spin-flipping penalty (Eq. (11)) measures the probability of local (“short-switching”) phasing errors; the block-switching penalty (Eq. (12)) measures the probability of long-range switching errors. Both penalty scores show bimodal distributions (Fig. S1). Sites with low spin-flipping penalty scores are enriched with false variants (Fig. S1A,D); by contrast, a large fraction of sites with low block-switching penalty scores are found in regions of low variant density (Fig. S1C,F).

We can generate haplotype blocks with different levels of accuracy using different block-switching penalty cutoffs. This is illustrated in Fig. 4 using Chr.5 as an example for both samples. As expected, choosing lower block-switching cutoffs produces longer haplotype blocks with more intra-block switching errors than choosing higher cutoffs. The low-accuracy blocks (colored in red) usually contain only one or a few switching errors at sites with low block-switching penalty scores: these blocks are broken into two or more high-confidence blocks at a higher block-switching cutoff (red arrows). As the main goal of local haplotype inference here is to generate high-confidence blocks that can be concatenated by long-range Hi-C links, we have chosen conservative block-switching cutoffs to produce short haplotype blocks with high accuracy. The cutoffs are determined based on the location of the minimum in the block-switching penalty distribution (Fig. S1B,E): Δ*E* 1000 for the RPE-1 data and Δ*E* 5000 for the NA12878 data. The resulting haplotype blocks are much shorter than reported previously (Marks et al., 2019), but the uniform accuracy (98%) of each block ensures consistent accuracy across the entire chromosome once all the blocks are merged.

**Figure 4:**
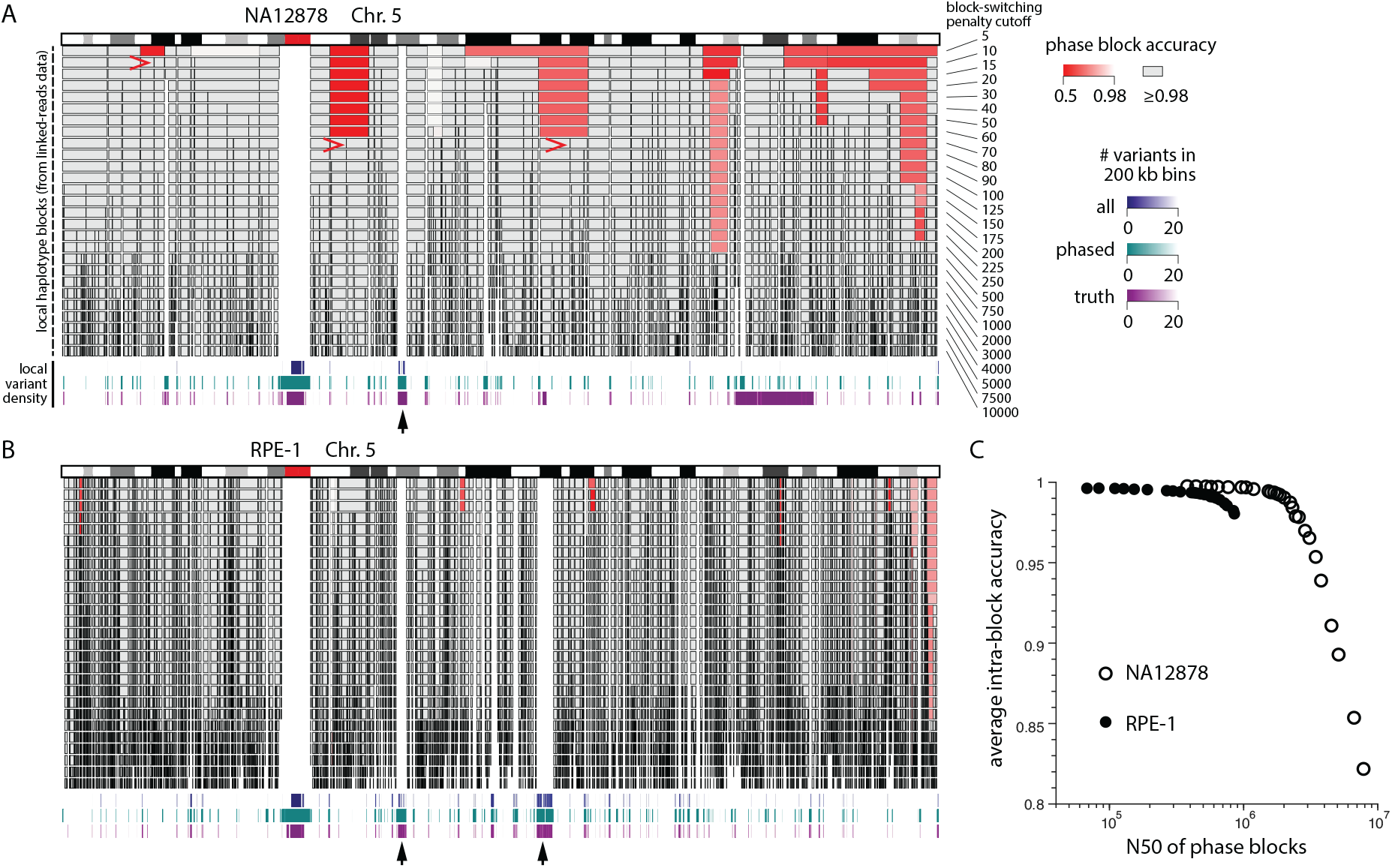
Haplotype blocks on Chr.5 inferred from the NA12878 and the RPE-1 linked-reads data. In panel 4A (NA12878) and B (RPE-1), each row represents haplotype blocks determined using a different switching-penalty cutoff (from Δ*E* 5 to Δ*E* 10, 000). Only blocks with 50 or more phased variants are shown. The accuracy of each block is estimated by the percentage of genotypes that are consistent with the majority haplotype of each block determined using the reference haplotype. Blocks with 98% accuracy are colored in grey; those with <98% accuracy are colored in red with brightness scaled by the accuracy. Three examples of low-accuracy blocks that are broken into two separate high-accuracy blocks at a higher block-switching cutoff are highlighted by red arrows in the NA12878 genome. Shown below the haplotype blocks are three tracks of regional variant density corresponding to the number of total detected variants (blue), phased variants in the final haplotype solution (green), and phased variants in the reference data (purple) in 200 kb bins. Bins with more than 20 variants (variant density more than 1 per 10 kb) are omitted. Black arrows highlight examples of large variant poor regions. Panel C shows the average intra-block accuracy (weighted by the number of variants in each block) and the N50 lengths of all haplotype blocks in each genome generated using different switching-penalty cutoffs. The NA12878 dataset produces longer haplotype blocks due to having longer input molecules (see Table 1).

**Figure 5:**
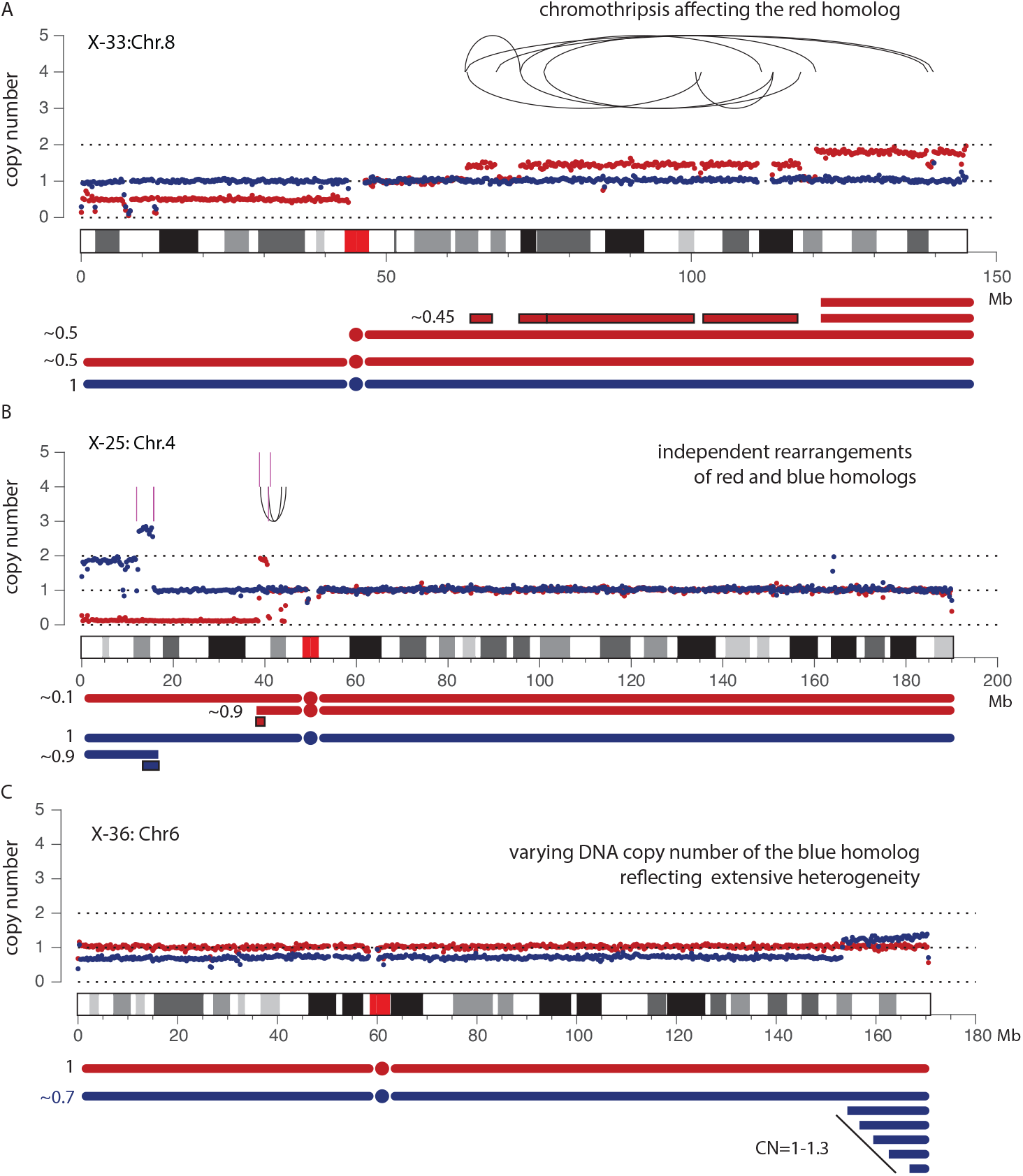
Haplotype-resolved DNA copy number and rearrangement analysis of post-crisis RPE-1 cells from Ref. Maciejowski et al. (2015). Schematic diagrams of alterations to each homolog and their clonal fractions are shown below the copy-number plots. Non-telomeric chromosomal fragments involved in complex rearrangements are outlined. A. Haplotype-specific DNA copy number (blue and red dots for each parental haplotype) and rearrangement (black arcs) of Chr.8 in the X-33 sample. Both the rearrangements and the copy-number alterations are restricted to the red homolog; the non-integer copy-number states indicate that the altered Chr.8 is present in a subclonal population (~45%). B. Haplotype-specific DNA copy number and rearrangement of Chr.4 (black arcs: intra-chromosomal; magenta vertical lines: inter-chromosomal to Chr.13) in the X-25 sample. Rearrangements and copy-number alterations affect both homologs. Both the gain of the blue homolog and the loss of the red homolog appear to be subclonal (~90%). C. Haplotype-specific copy number of Chr.6 in the X-36 sample. The q-terminus of the blue homolog shows non-constant copy number (1-1.3) that contrasts with the stable copy number of the red homolog or the rest of the blue homolog (∼0.7). We interpret this pattern as reflecting extensive genetic heterogeneity in the population.

We note that haplotype blocks are often terminated at regions of low-variant density (less than 1 per 10 kb). (Also see Figs. S1C,S1F,S2 and S3.) Two examples of large low-variant density regions are highlighted in the RPE-1 genome (black ar-rows in Fig. 4B). The first one in 5q13.2 is also seen in the NA12878 genome: This region, known as the spinal muscular atrophy (SMA) region, contains large segmental duplications (~200kb) with high sequence similarity (98%) (Schmutz et al., 2004) that cannot be resolved by short sequencing reads. Even though the standard variant caller (HaplotypeCaller) emits many variants in this region with variant density above 1 per 10kb (blue tracks), most of these variants are false and left out in the reference hap-lotype (purple tracks). They are largely excluded from the final haplotype solution (green tracks) due to inconsistent or missing haplotype linkage. By contrast, the second low-variant density region in the RPE-1 genome near 5p21.1 contains few variants and most likely reflects loss-of-heterozygosity. The exclusion of both regions in the final haplotype solution confirms the robustness of our haplotype inference algorithm against false variants with inconsistent haplotype linkage.

We merge haplotype blocks using Hi-C links in two steps. First, haplotype blocks within each chromosome arm are concate-nated using Hi-C links between variants separated by 10 Mb. Second, p- and q-arm haplotypes are joined using all Hi-C links between the arms. The consistency of haplotype solution in each step can be verified by comparing the number of *cis* and *trans* Hi-C links (Fig. S4). To expedite convergence of the haplotype solution, we exclude short haplotype blocks with 5 total Hi-C links to other blocks from the calculation. We refer to the concatenated haplotype blocks as the “scaffold haplotype.”

After generating the scaffold haplotype, we then calculate the linkage between variant genotypes and the scaffold haplotype using

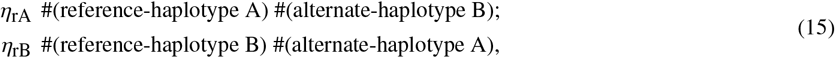

Here the counts reflect the number of unique molecules linking each genotype (Reference or Alternate) to either parental haplotype (A or B). We used the linkage to scaffold haplotypes to assign the haplotype phase at each variant site in the final haplotype solution.

### Variant filtration using haplotype linkage

For true heterozygous variants, we expect the variant genotypes to be phased to different parental haplotypes: η_rA_ » η_rB_ ≈ 0 or η_rB_ » η_rA_ ≈ 0. We implemented a “linkage filter” to exclude false variants in the final haplotype solution based on the following criteria of haplotype linkage:

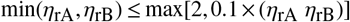

with at least one link to both genotypes (R and A) and both scaffold haplotypes (A and B).

### Benchmark of the haplotype solution

We evaluated the accuracy and completeness of the computationally inferred haplotypes using independent reference haplotype data (Table 1). For the NA12878 genome, the reference haplotypes were determined from the parental genomes and released by the Genome-In-A-Bottle consortium. For the RPE-1 genome, we determined the reference haplotypes from the sequencing data of monosomic RPE-1 cells. The NA12878 haplotype data have high specificity but leave out several large regions including the p-arms of Chrs.16 and 18. These regions were excluded from benchmarking. For the reference RPE-1 haplotype data, we additionally implemented a variant filter based on the average variant allele fraction in all the single-cell data.

We first benchmarked the scaffold haplotype solution constructed from large haplotype blocks (Table S2 for the NA12878 dataset and Table S3 for the RPE-1 dataset). For the NA12878 sample, the scaffold haplotype solution contained 94% of all phased variants in the reference haplotype data with 99.6% accuracy. (Chromosome 19 has the lowest accuracy of 98.5%.) For the RPE-1 sample, the scaffold haplotype solution contained 89% of all phased variants in the reference haplotype data and showed 98.3% agreement. (Chromosome 9 has the lowest percentage of agreement of 96.1%.) There is no long-range switching error in any chromosome in either sample.

We then benchmarked the final haplotype solution determined using the linkage between variant genotypes and the scaffold haplotypes. For the NA12878 sample, the final haplotype solution showed 99.7% accuracy and 98.0% completeness when compared to the reference haplotype (Table 2). The linkage filter removed 167,385 variants but did not affect the benchmarks as most of the false variants were already excluded from the reference haplotype.

**Table 2:** Comparison between the final haplotype solution and the reference haplotype of NA12878.

| Total variant sites | Phased from bulk data | Phased from parental data | Comparable sites | Agreed | Accuracy | Fraction of completion |
| --- | --- | --- | --- | --- | --- | --- |
| 2,652,381 | 2,319,027<sup>a</sup> | 1,861,941 | 1,824,401 | 1,818,042 | 0.997 | 0.980 |
|  | 2,151,642<sup>b</sup> | 1,815,212 | 1,815,197 | 1,809,886 | 0.997 | 0.975 |

<sup>a</sup>All phased variants without any filtering
<sup>b</sup>Filtered by haplotype linkage: $\geq 1$ link connecting ref, alt, HapA, and HapB, and minor linkage $\leq 2$ or minor linkage/total linkage $\leq 0.1$

For the RPE-1 sample, the final haplotype solution showed 98% agreement with the reference data before variant filtration (Table 3). We expected a large percentage of the discrepancy to be due to false variants. After excluding false variants based on either the variant allele fraction in the single cell data (100 samples) or the haplotype linkage in the linked-reads data, we saw 99% agreement between the haplotype solution and the reference haplotype. Although the linkage filter and the allele fraction filter are completely independent, they have good consistency: 2,054,859 variants pass both filters and represent 95% of variants passing each individual filter. Among variants passing both filters, the percentage of agreement between the haplotype solution and the reference haplotype is 99.6% and comparable to the NA12878 haplotype solution.

**Table 3:** Comparison between the final haplotype solution and the reference haplotype of RPE-1.

| Filter | Total variant sites | Phased from bulk data | Phased from monosomies | Agreed | Discordant | Fraction of discordance |
| --- | --- | --- | --- | --- | --- | --- |
| None | 2,475,311 | 2,242,237 | 2,320,153 | 2,101,195 | 40,006 | 0.019 |
| Allele fraction<sup>a</sup> | 2,172,689 | 2,087,188 | 2,109,589 | 2,018,906 | 12,903 | 0.006 |
| Linkage | 2,156,423 | 2,156,346 | 2,071,674 | 2,054,006 | 17,616 | 0.009 |
| Combined<sup>b</sup> | 2,054,859 | 2,054,832 | 2,001,674 | 1,993,552 | 8,098 | 0.004 |

<sup>a</sup>From single-cell data: minor allele fraction $\geq 0.3$ in disomic regions and $\in [0.18, 0.48]$ in the trisomic region in Chr.10q.
<sup>b</sup>With both the allele fraction (a) and the linkage (b) filter

### Resolving chromosome-specific alterations using haplotype copy number

To demonstrate this application, we use the parental RPE-1 haplotype information to resolve large structural alterations of chromosomes in RPE-1 cells generated by telomere fusions (Maciejowski et al., 2015). In this study, the authors performed bulk whole-genome sequencing on the progeny populations of single RPE-1 cells that underwent telomere crisis. We downloaded and reprocessed the sequencing data using our standard workflow and calculated haplotype coverage in 250kb bins using the RPE-1 haplotype data as

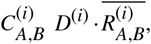

where *D*^(*i*)^ is the normalized average sequence coverage in the *i*th bin (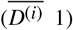 1) and 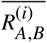 is the mean haplotype fraction (A or B) across all variants within the *i*th bin. We then calculated haplotype DNA copy number by normalizing *C*_*A*,*B*_ by its median value across the whole genome (from both haplotypes). The median value corresponds to the expected (normalized) coverage of a single chromosome (haplotype) assuming the genome is mostly diploid.

Figure 6 shows three examples of complex chromosomal alterations, each taken from a different sample that underwent telom-ere crisis. Haplotype-specific copy number is shown using red and blue dots; chromosomal rearrangements accompanying the copy-number alterations are shown as black arcs (intra-chromosomal events) and magenta vertical lines (breakpoints of inter-chromosomal translocations). The first example (Fig. 6A) shows a chromothripsis event affecting the 8q arm of the red homolog. Based on the non-integer copy-number states of the red haplotype and the near diploid karyotype of this sample (Maciejowski et al., 2015), we infer that the altered Chr.8q is present in a subclonal population (~45%). The second example (Fig. 6B) shows alterations to both Chr.4 homologs on the p-arm: Both the gain of the blue haplotype and the loss of the red haplotype are sub-clonal (~90%); the broken ends on both homologs are linked to Chr.13 (magenta lines), suggesting a complex event involving three separate chromosomes. The last example (Fig. 6C) shows an intriguing pattern of copy-number variation. The varying copy number of the blue haplotype at the q-terminus contrasts with the constant copy number of the red haplotype or the rest of the blue haplotype, excluding the possibility that the variation is due to technical artifacts. We interpret this varying copy number pattern as the outcome of varying terminal segments present in different cells in the population (Umbreit et al., 2020).

**Figure 6:**
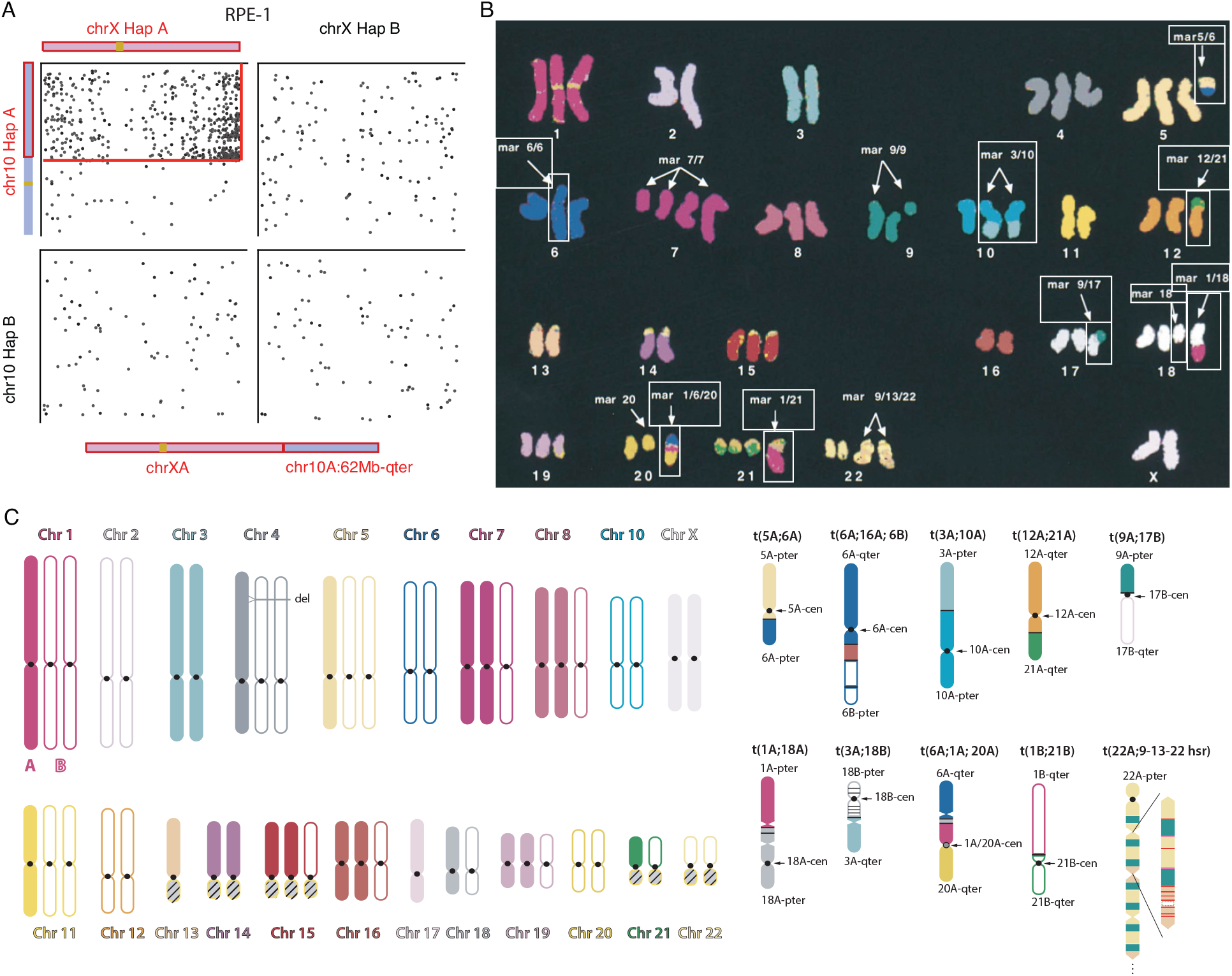
Haplotype-resolved structure of derivative chromosomes and aneuploid karyotypes. A. Walking the translocated X chromosome in the RPE-1 genome using phased Hi-C links (dots) between different homologs of Chr.10 and Chr.X. A significant enrichment of Hi-C links is seen in only one haplotype combination reflecting the translocation between Chr.10 and Chr.X. The enrichment of Hi-C links throughout the entire X chromosome suggests the composition of the derivative chromosome as shown at the bottom. B. The cytogenetic K-562 karyotype reported in Ref. (Gribble et al., 2000) with outlined marker chromosomes resolved by sequencing data and shown in C. C. The digital K-562 karyotype determined from linked-reads and Hi-C sequencing data (Table S1), with haplotype assignment to both normal (left) and marker (right) chromosomes. The digital karyotype mostly agrees with the cytogenetic karyotype and the differences may be attributed to additional alterations during cell culture. Among all the marker chromosomes listed in Panel B, we are able to determine the syntenic structure of the following: mar5/6, mar6/6, mar3/10, mar12/21, mar9/17, mar1/18, mar1/6/20, and mar1/21, and resolve most translocation junctions at the base pair level. Multiple junctions contained local fold-back rearrangements (opposite arrows in t(1A;18A), t(3A;18B), t(6A;1A;20A)) that are consistent with local DNA copy number gains; these events cannot be resolved by cytogenetic analyses. The mar18 described in Ref. (Gribble et al., 2000) is probably related/similar to t(3A;18B). The *BCR-ABL* amplification is contained in a homogeneously staining region (hsr) of a marker chromosome t(22A;9-13-22hsr). We infer the structure of the amplicon from DNA copy number and rearrangements but cannot validate the inferred structure due to technical limitations. We are further able to partially determine the structures of the altered Chr.7 and Chr.9 and resolve three additional marker chromosomes described in Ref. (Naumann et al., 2001) but not in Ref. (Gribble et al., 2000): t(2A;22A), t(3A;10A;17A), t(9A;13A). These results are summarized in Additional Dataset 2.

The haplotype copy-number analysis demonstrates that the progeny populations of single cells passing through telomere crisis can be highly heterogeneous. The feature of non-constant haplotype copy number is of particular interest and may be used as a signature to infer ongoing genome instability in a cell population directly from bulk DNA sequencing.

### Walking derivate chromosomes using haplotype-specific Hi-C contacts

Hi-C sequencing has previously been used for detecting long-range chromosomal rearrangements (Rao et al., 2014; Dixon et al., 2018; Zhou et al., 2019). The formation of new junctions between distal loci (separated by 1 Mb genomic distance or located on different chromosomes) creates new *cis* contacts with a significantly higher density than *trans* contacts in a normal genome. As each rearrangement breakpoint is generated on one parental chromosome, the newly formed *cis* contacts at the rearrangement junction should be phased to one haplotype on either side of the junction. For inter-chromosomal rearrangements, *cis* contacts between the parter chromosomes should be observed in one out of four possible haplotype combinations (AA, AB, BA, or BB); for intra-chromosomal rearrangements, newly formed *cis* contacts should be observed in one out of three possible combinations (AA, AB, or BB). Combining haplotype-specific connectivity from Hi-C contacts with haplotype DNA copy number from linked-reads data enables us to infer the syntenic structure of derivative chromosomes and generate phased karyotypes (Fig. 6).

We first illustrate this approach using a simple example in the RPE-1 genome (Fig. 6A). RPE-1 cells contain an extra copy of a segment from Chr.10q (62 Mb-qter). The DNA sequence near the breakpoint on Chr.10q shows repeats whose origin cannot be determined; cytogenetic analysis indicates that this segment is fused to the q-terminus of Chr.X. In the phased Hi-C contact map, this translocation is easily recognized from the enrichment of contacts near the q-terminus of Chr.X and the breakpoint on Chr.10q (~62Mb) that is restricted to one haplotype combination (arbitrarily denoted as A for both chromosomes). Importantly, the enrichment of Hi-C contacts extends throughout Chr.X to the p-terminus, indicating that the 10q segment joins a complete X chromosome and confirming the result from cytogenetic analysis.

Using a similar strategy, we are able to generate a “digital karyotype” of the K-562 genome using published sequencing data (Table S1). The K-562 genome is highly aneuploid (Zhou et al., 2019) and contains multiple marker chromosomes (Gribble et al., 2000; Naumann et al., 2001) and large regions of loss-of-heterozygosity (LOH). We determined the parental haplotypes in heterozygous regions and then calculated haplotype-specific DNA copy number using phased links in the linked-reads data and generated haplotype-specific Hi-C maps using Hi-C contacts that are phased on at least one end. The digital karyotype was determined by a joint analysis of haplotype-specific DNA copy number, rearrangements, and Hi-C contacts (SI Additional Data). The digital karyotype is schematically shown in Fig. 6C and shows excellent agreement with the cytogenetic karyotype reported in Ref. (Gribble et al., 2000) (Fig. 6B) and (Naumann et al., 2001). The digital karyotype shows excellent agreement with the cytogenetic karyotype but provides haplotype resolution of the syntenic blocks and base-pair resolution of the translocation junctions in 9 marker chromosomes reported in Ref. (Gribble et al., 2000) (outlined in Fig. 6B and schematically shown in Fig. 6C) and 3 additional marker chromosomes in (Naumann et al., 2001). We are also able to partially resolve the structure of the complex amplicon containing the BCR-ABL fusion in t(22A;9-13-22hsr) combining sequencing and cytogenetic data. Details of the reconstructed marker chromosomes are provided in SI additional Data.

## Discussion

Here we describe a computational method that can accurately determine complete chromosomal haplotypes from a combination of linked-reads sequencing (30-60× mean depth) and Hi-C sequencing (≥50 million long-range contacts). The computationally inferred haplotypes show high accuracy (99%) and completeness (98%) when compared to independent reference haplotypes.

Our method offers several advantages over previous methods. First, both linked-reads and Hi-C sequencing data can be generated on standard sequencing platforms and the construction of sequencing libraries does not involve special experimental techniques required for single-chromosome isolation (Ma et al., 2010; Fan et al., 2010; Yang et al., 2010), single-cell sequencing (Zhang et al., 2015), or similar techniques such as “Strand-Seq” (Falconer et al., 2012; Porubsky et al., 2017). Second, the computational algorithm implicitly excludes inconsistent linkage evidence from false variants based on the specificity of haplotype linkage. This contrasts with previous methods (Edge et al., 2017) that require high-quality variants as input. Our method further enables a variant-filtering strategy based on haplotype linkage that can be used to exclude false variants due to alignment errors and validate complex variants such as small insertions and deletions, or large structural variants.

Our formalism of haplotype inference as a minimization problem also has several unique features. The symmetric representation of binary genotypes and haplotypes simplifies the inference of complementary haplotypes into one minimization problem based on linkage evidence from both parental chromosomes. The haplotype inference algorithm is not affected by allelic imbalance, including loss-of-heterozygosity, and is directly applicable to aneuploid tumor genomes. We demonstrate that a simple iteration strategy can efficiently solve the parental haplotypes of diploid genomes but it is straightforward to incorporate more sophisticated minimization algorithms (*e.g.*, Monte-Carlo methods) when necessary. One useful extension of our method is to perform joint haplotype inference using population genotypes and Hi-C data. Population-based statistical phasing (Loh et al., 2016a) can produce long haplotype blocks (1Mb) that contain random but rare switching errors. It should be possible to correct these errors using Hi-C data and extend the statistical haplotype phase to the entire chromosome.

Knowledge of chromosomal haplotypes can be used to directly relate variations in the DNA sequence, histone marks, chromatin structure, and gene expression on each chromosome. This is especially useful for the analysis of cancer genomes where homologous chromosomes often acquire independent alterations in the DNA sequence that can cause differential changes in chromatin folding and gene expression (Spielmann et al., 2018). We demonstrate the capability to resolve the composition of derivative chromosomes and aneuploid genomes by constructing a digital karyotype of the K-562 genome from linked-reads and Hi-C sequencing data. We expect this strategy to be generally applicable to complex cancer genomes. The capability to determine the large-scale structure of derivative chromosomes shall enable us to further investigate the connection between 2D chromosomal structural alterations and 3D chromatin reorganization.

## Supporting information

Extended method description and supplementary discussion

Supplementary figures related to the analysis of K-562 karyotype.

