## Extended method description and supplementary discussion for "Determination of complete chromosomal haplotypes by bulk DNA sequencing"

June 16, 2020

### Materials and Method

#### Data generation

The sources of public data used in this study are listed in Table 1 (RPE-1 and NA12878) and in Table 1 (K-562).

##### Bulk linked-reads sequencing data of RPE-1 cells

The RPE-1 linked reads data were generated at the Yale Center for Genome Analysis (`https://medicine.yale.edu/keck/ycga/`). High-molecular weight DNA from RPE-1 cells was extracted using the RevoluGen PuriSpin Fire Monkey kit following the protocol provided by the vendor with the following modifications: Cells were lysed at 56°C for 2 hours, followed by addition of ~100 ng RNase A and additional incubation for 15 minutes at 56°C. A single linked-reads library was constructed using the Chromium Genome Library Kit v2 from 10X Genomics following the standard protocol. The library was then sequenced on the Illumina NovaSeq platform to generate 941,518,426 read pairs with 60× mean depth of coverage. The mean size of input DNA was estimated to be 25 kb by the LongRanger software. See Table 1 for additional metrics of the sequencing data.

##### Sequencing data of monosomic RPE-1 cells

Monosomic RPE-1 cells were generated using three different strategies: (1) Nocodazole block and release (Zhang et al., 2015); (2) Induction of dicentric chromosome bridges (Umbreit et al., 2020); (3) Treatment with Paclitaxel, a spindle toxin that prevents micrutubulin disassembly. All three strategies significantly increase the frequency of chromosome missegregation and the generation of monosomic daughter cells. Monosomic cells were first selected based on the arm-level DNA copy number estimated from low-pass ( 0.1×) whole-genome sequencing and then sequenced to 5-30× on either the Illumina HiSeq 2500 or the Illumina NovaSeq platforms at the Broad Institute of MIT and Harvard. We then identified and validated completely monosomic chromosomes using the deep sequencing data based on the observed heterozygosity normalized by the mean allelic coverage (Zhang et al., 2015):

$$\frac{\text{observed heterozygosity}}{(\text{observed allelic coverage})^2} = \frac{p_{\text{het}}}{(p_{\text{ref}} + p_{\text{alt}})^2/4}.$$

The **observed heterozygosity** $p_{\text{het}}$ is determined as the fraction of parental heterozygous sites that show heterozygous coverage in a single-cell genome; the **observed allelic coverage** is estimated using the average of the fraction of heterozygous sites with reference coverage $p_{\text{ref}}$ and the fraction of heterozygous sites with alternate coverage $p_{\text{alt}}$, which approximates the average coverage of each parental chromosome in disomic regions in a single cell genome (Zhang et al., 2015). Heterozygous variants in the parental genome were detected using the bulk sequencing data as described below in Variant calling and filtering. To eliminate false heterozygosity due to sequencing or amplification errors, we consider a variant site to show reference or alternate coverage only when the number of sequencing reads showing either genotype exceeds a threshold set as $d^* = \max(2, 1 + 0.1 \times \text{mean sequencing depth of chromosome})$: $d^* = 2$ if the mean sequencing depth is $\leq 10\times$ (most samples) and $d^* = 4$ if the mean sequencing depth is 30×. The minimum threshold of 2 reads is used to eliminate sequencing errors; the threshold of 0.1× mean sequencing depth serves to exclude low frequency (<10%) amplification errors. Complete monosomies are selected based on the criteria that the observed heterozygosity is less than 0.1× the median from all the cells ($\approx 1$). For the current study, we selected 39 cells with one or multiple monosomic chromosomes (32 from nocodazole release, 5 from bridge induction, and 2 from Paclitaxel treatment), containing 98 monosomic chromosomes in total. The sample names, mean sequencing depths, and the estimated heterozygosity of monosomic chromosomes are listed in Additional Data Table 3.

#### Data processing

All the sequencing data listed in Table 1 and 1 were re-processed starting from unmapped sequencing reads. For the linked-reads data, we used the LongRanger software from 10X Genomics to extract the molecular barcode of each sequencing fragment that is then preserved in the "BX" tag in the BAM record. The molecular barcode information was only used as molecular linkage evidence but not for sequence alignment. Alignment and post-alignment processing of all sequencing data except the K-562 linked-reads data were completed using the same pipeline as described below. For the K-562 linked-reads data, we used the output from LongRanger for downstream analysis.

##### Sequence data alignment

We aligned all sequencing data (both linked reads and Hi-C) using a standard short-read aligner (`https://github.com/lh3/bwa`) with default parameters ("`bwa mem`"). Using a barcode-agnostic aligner ensures better specificity of linkage information (and therefore better phasing accuracy) than using a barcode-aware aligner such as `Lariat` (`https://github.com/10XGenomics/lariat`) that is included in the LongRanger pipeline. The rationale is explained below in Linkage evidence from molecular identifier and sequence alignment.

##### Post-alignment processing

We generated the insert size histogram of sequencing fragments from the insert size inferred from the alignment positions of 2,000,000 uniquely aligned fragments (both mates having mapping quality 60) with proper alignment positions (two mates are placed at the forward-reverse orientation with < 2000 bp separation). The lower and upper limits of the insert size was set to be the 0.1% and 99.9% percentile of the insert size histogram. When choosing the primary alignment positions of sequencing reads with multiple alignment positions (supplementary or secondary alignments), we gave preference to alignment positions consistent with the proper-pair configuration, *i.e.*, placing the two pairmates at the forward-reverse orientation and within the insert size range. We then used the `MarkDuplicates` program in Picard (`https://broadinstitute.github.io/picard/`) to infer sequencing reads corresponding to PCR duplicates based on the primary alignment positions, and adjusted the duplication tag of both primary and supplementary alignments accordingly.

#### Variant calling and filtering

We ran the `HaplotypeCaller` program from GATK (v4.0.12.0-6-gfef36e3-SNAPSHOT) in the discovery mode ("`--genotyping-mode DISCOVERY`") to detect genetic variants. We imposed the following read filters in addition to the standard parameters and read filters used by `HaplotypeCaller` to exclude reads with improper or inaccurate alignment or low mapping quality:

```
--read-filter PairedReadFilter \
--read-filter MateOnSameContigOrNoMappedMateReadFilter \
--read-filter FragmentLengthReadFilter --max-fragment-length 1000 \
--read-filter MateDifferentStrandReadFilter \
--read-filter MappingQualityReadFilter --minimum-mapping-quality 30 \
--read-filter OverclippedReadFilter --filter-too-short 25 \
--read-filter GoodCigarReadFilter --read-filter AmbiguousBaseReadFilter
```

For the RPE-1 genome, variant discovery was performed jointly on the new linked-reads data (60×) and the previously published standard whole-genome data (13×) (Zhang et al., 2015). For the NA12878 genome, variant discovery was performed on two public linked-reads datasets (35× each).

We selected bi-allelic single-nucleotide variant sites (one reference plus one alternate) as the input for haplotype inference, excluding sites in pericentric, acrocentric, and centromeric regions based on the standard chromosome banding annotation ("`acen`","`gvar`","`stalk`") provided by the UCSC genome browser. No other quality filter (such as variant quality score recalibration) was applied.

#### Software implementation of the haplotype inference algorithm

We have implemented a C++ package "`linker`" (`https://github.com/rwtourdot/linker`) that performs multiple tasks related to haplotype inference.

##### Extracting variant linkage information from long-range sequencing

Taking long-range sequencing data and a set of heterozygous variants as input, "`linker extract`" generates a hash map from variant genotypes to sequencing reads ("variant-to-read") by iterating over all sequencing reads overlapping each variant site. This module can be executed either on an entire chromosome or on specified regions of interest. The output are plain text files with each line listing the names/identifiers of sequencing reads showing a specific variant genotype. For example,

```
198801_A_C -> {read_identifier1, read_identifier2, read_identifier3};
```

represents the following

```
198801: variant position;
A: reference base;
C: genotype observed in the supporting reads (read_identifier1, etc.).
```

For linked-reads data, `linker` uses the 16-bp molecular barcodes (usually stored in the "BX" Tag) as read identifiers. For Hi-C data, `linker` uses read names as read identifiers. For long-read data, the default read identifier is constructed by concatenating the read name with the start and end positions of alignment; this definition ensures that only linkage between variants within a single contiguous alignment is preserved, but not between variants in non-contiguous (split) alignments of the same read. `linker` additionally applies the following read filters (these will be customizable in a future release):

| Data type | Mapping quality | Base Quality | Bam Flag |
|---|---|---|---|
| linked-reads | < 20 | < 20 | duplicate, non-primary |
| long-read | < 20 | < 8[a] | duplicate, non-primary |
| Hi-C | < 20 | < 20 | duplicate |

[a]This threshold applies to uncorrected PacBio reads but not to corrected PacBio reads.

As `linker` iterates over all variants to generate the variant-to-read map, it simultaneously creates the inverse map from reads to linked variants. Each line in the "read-to-variant" map lists all the variant genotypes linked by each read/molecule. For example,

```
read_identifier1 -> {198801_A_C, 198322_T_T, 196990_C_G};
```

indicates that the genotypes `C`, `T`, and `G` at positions 198801, 198322, and 196990 are linked by a single read or linked-read molecule with identifier `read_identifier1`. The variant-to-read and the read-to-variant maps are the only input data for downstream analysis.

#### Calculating haplotype-specific linkage between variants

The signal of haplotype linkage between two variant genotypes (*e.g.*, `198801_A_C` and `198322_T_G`) is measured by the number of unique reads (molecules) linking these genotypes, which is calculated by intersecting the list of reads associated with each variant genotype (using the variant-to-read map). The calculation of haplotype linkage is embedded in the "`linker solve`" module prior to haplotype solution. We have implemented a separate module"`linker matrix`" that calculates and outputs haplotype linkage between variants on each chromosome.

#### Solving haplotype phase by minimization

`linker solve` takes the variant-to-read and the read-to-variant hash maps as input and finds the optimal haplotype solution by alternately performing spin flipping or block switching to lower the energy function given by Eq. (10). During each round of minimization, the program first performs "spin flips" at sites with negative flipping energy penalties calculated using Eq. (11); it then performs block switches between sites with negative switching energy penalties calculated using Eq. (12). In calculating the block switching energy $\Delta E_{k|k+1}$, we haven taken advantage of the following recursive relationship

$$\begin{aligned}\Delta E_{k|k+1} - \Delta E_{k-1|k} &= \sum_{i\le k}\sum_{j>k} M_{ij} s_i s_j - \sum_{i<k}\sum_{j\ge k} M_{ij} s_i s_j \\ &= s_k\left(\sum_{j>k} M_{kj} s_j - \sum_{i<k} M_{ik} s_i\right) \\ &= s_k\left(\sum_{i>k} M_{ki} s_i - \sum_{i<k} M_{ki} s_i\right). \end{aligned} \tag{S1}$$

The last step uses the symmetric property of $M_{ik} = M_{ki}$. Introducing

$$h_i^+ = \sum_{j>i} M_{ij} s_j \quad \text{and} \quad h_i^- = \sum_{j<i} M_{ij} s_j, \tag{S2}$$

we can calculate the spin flipping energy $\Delta E_i$ and the block switching energy $\Delta E_{k|k+1}$ using

$$\Delta E_i = s_i\left(h_i^+ + h_i^-\right) \quad \text{and} \quad \Delta E_{i|i+1} = \Delta E_{i-1|i} + s_i\left(h_i^+ - h_i^-\right). \tag{S3}$$

Therefore, the calculation of both energy penalties over all variant sites has the same complexity as the calculation of $h_i^+$ and $h_i^-$, which is proportional to the total number of non-zero $M_{ij}$'s that is approximately $N \times L$, where $N$ is the total number of variants, $L$ is the average number of variants covered by each link.

The energy penalties are calculated iteratively from the first spin to the last using Eqs. (S2) and (S3) while the spin configuration is continuously updated. This asynchronous iteration scheme achieves faster convergence to an energy minimum than synchronous iteration in which all spins are updated at once. We illustrate the difference between these two iteration strategies using a toy example shown below.

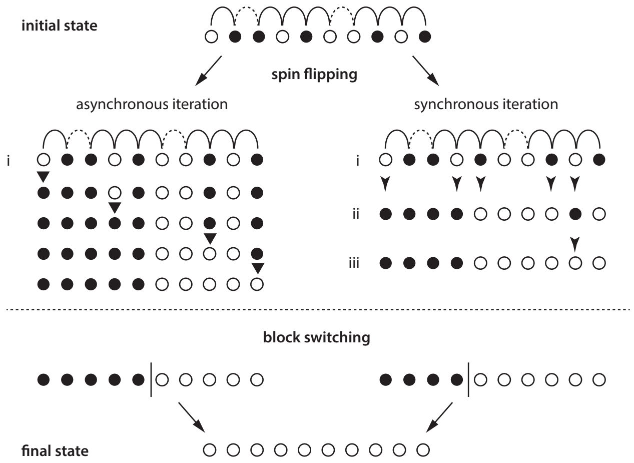


In this example, we start from a random initial configuration ($s_i^{(0)} = \pm 1$) with open and filled circles representing genotypes in complementary haplotypes. The interactions between adjacent genotypes are shown as arcs above the haplotype configuration. The interaction is positive (dashed arc) between genotypes in the same haplotype and negative (solid arc) between genotypes from opposite haplotypes. Spin flips are introduced if the net interaction with adjacent spins is negative. In asynchronous iteration (left), the energy penalty is calculated as spin flips are introduced: In one round of iteration, the 1st, 4th, 8th, and 10th spins are flipped due to negative interactions with the neighbor spins. In synchronous iteration (right), the 5th and the 9th spins are also flipped due to negative interactions with their neighbors in the initial state (ii); it will take another round of iteration to reverse the 9th spin to the optimal configuration (iii). The switching between two haplotypes at the end of spin flipping can only be resolved through a block switching move.

#### Assessing phasing accuracy and determining high-confidence haplotype blocks

At the end of iteration, "`linker solve`" outputs a haplotype solution consisting of a numerical genotype (+1 or −1) and two energy penalty scores at each variant site calculated using Eqs. (11) and (12). The energy penalty scores incorporate site-specific linkage error estimates (Eq. (6)) and measure the confidence of haplotype inference based on linkage evidence (Eq. (13)). High-confidence haplotype blocks are determined by disassembling the haplotype solution at sites with low block-switching penalty scores (high probability of switching errors).

#### Concatenating haplotype blocks using Hi-C links

We have implemented `linker scaffold` for this step. The program first determines high-confidence haplotype blocks from the haplotype solution generated by `linker solve` based on a block-switching penalty cutoff (default value $\Delta E = 700$) that can be specified as an optional input parameter. The program then constructs maps between variants and haplotype blocks:

```
198801 -> haplotype_block_1;
198322 -> haplotype_block_1;
haplotype_block_1 -> {198801, 198322, ...};
haplotype_block_2 -> {199005, 200142, ...}.
```

Phased genotypes within each haplotype block are represented as (1 for reference and -1 for alternate):

```
haplotype[haplotype_block_1]=[1, -1, -1, ...].
```

The phased linkage between haplotype blocks is calculated by iterating over all Hi-C reads spanning variants in different blocks (using the read-to-variant map of Hi-C data generated by `linker extract`). For example, the number of links supporting *cis*-linkage between two haplotype blocks $\mathbf{B}_s$ and $\mathbf{B}_t$ is given by

$$n_{st}^{+} = n(\mathbf{B}_s \leftrightarrow \mathbf{B}_t) + n(\overline{\mathbf{B}}_s \leftrightarrow \overline{\mathbf{B}}_t) = \#\left[\sigma^{(m)}(x_m)\sigma^{(m)}(y_m)\mathbf{B}_s(x_m)\mathbf{B}_t(y_m) = 1\right], \tag{S4}$$

where the count runs over all Hi-C links $\{\boldsymbol{\sigma}^{(m)}|m = 1,2,\cdots\}$ with variant positions $x_m$ and $y_m$ in the two haplotype blocks $\mathbf{B}_s$ and $\mathbf{B}_t$. Similarly,

$$n_{st}^{-} = n(\mathbf{B}_s \leftrightarrow \overline{\mathbf{B}}_t) + n(\overline{\mathbf{B}}_s \leftrightarrow \mathbf{B}_t) = \#\left[\sigma^{(m)}(x_m)\sigma^{(m)}(y_m)\mathbf{B}_s(x_m)\mathbf{B}_t(y_m) = -1\right], \tag{S5}$$

gives the signal of *trans*-linkage. These are used for calculating inter-block linkage given by Eq. (9).

When concatenating haplotype blocks, `linker scaffold` first uses Hi-C links between variants within 10 Mb to solve the linkage between blocks within each chromosome arm using the same strategy as described in Solving haplotype phase by minimization. In this step, it also drops short haplotype blocks with ≤ 5 total Hi-C links to other blocks to expedite convergence. The p- and q-arm haplotypes are then joined by evaluating all phased Hi-C links between the two arms. If the haplotypes of both arms are solved correctly, then the Hi-C linkage between arms should show a strong bias (>10:1) dominated by intra-chromosomal linkage (Fig. S4). The output of `linker scaffold` is referred to as the "scaffold haplotype solution".

##### Refining the scaffold haplotype solution using haplotype linkage

The scaffold haplotype solution has left out short haplotype blocks consisting of variants with insufficient linkage evidence. We have implemented a "`linker recover`" module that can impute the haplotype phase at the eliminated variants using the linkage between these variants and the scaffold haplotype. The linkage between a variant genotype (*e.g.*, `198801_A_C`) and a haplotype solution (**S**) is calculated using two methods. The first method calculates haplotype linkage as the <u>sum of linkage evidence between the variant genotype of interest and all variant genotypes in the haplotype of interest</u> (Eq. (14)). This method counts a single read identifier (unique molecule) more than once if it covers more than two variant sites in the haplotype solution. The second method calculates haplotype linkage by intersecting the list of read identifiers associated to the variant genotype of interest with the union of read identifiers at all variant genotypes in the haplotype of interest (Eq. (15)), which is equivalent to <u>the number of unique molecules linking the variant genotype to the haplotype solution</u>. We use the second measure of haplotype linkage to determine the final haplotype solution and filter false variant sites with ambiguous linkage evidence.

### Supplementary Discussion

#### Linkage evidence from molecular identifier and sequence alignment

Molecular linkage between sequencing fragments is reflected in two features. First, sequencing fragments derived from the same DNA molecule should share the same molecular barcode:

$$\text{physical linkage} \Rightarrow \text{identical molecular barcode.}$$

Second, sequencing fragments derived from the same DNA molecule should map to proximal locations based on their sequences:

$$\text{physical linkage} + \text{correct alignment} \Rightarrow \text{proximity of alignment positions.}$$

The logic behind barcode-aware aligners (*e.g.*, Lariat) (Marks et al., 2019) is the following :

$$\left.\begin{array}{r}\text{identical molecular barcode} \dashrightarrow \text{physical linkage}\\ \text{sequence information}\end{array}\right] \rightsquigarrow \text{optimal alignment positions.}$$

This strategy can improve the placement of <u>sequence fragments with multiple possible alignment positions using the alignment positions of **uniquely aligned fragments** with the same molecular barcode</u>. For example, the alignment positions of sequence fragments derived from short interspersed repeats (typically <10 kb) may be anchored by the alignment positions of fragments in the flanking non-repeat regions. However, for sequence fragments derived from <u>regions of segmental duplications</u>, or from regions not represented in the reference genome (*e.g.*, centromeres), their true alignment positions cannot be uniquely determined or are not included in the reference after all. In these scenarios, barcode-aware aligners may reinforce false linkage evidence if these fragments are placed in proximity but at incorrect locations.

To ensure **the best specificity of linkage between sequence fragments**, we want to use molecular barcodes and alignment positions as independent evidence of linkage between sequence fragments. The alignment positions are determined solely based on the DNA sequence and do not use the molecular barcode information (*i.e.*, barcode-agnostic alignment). We consider sequence fragments to be linked in *cis* only when they both share the same molecular barcode and are aligned to proximal positions (< 100 kb) independent of the molecular barcodes:

$$\left.\begin{array}{r}\text{identical molecular barcode}\\ \text{proximity of alignment positions } (< 100\text{ kb})\end{array}\right] \Rightarrow \textit{cis}\text{ linkage between fragments.}$$

The 100 kb threshold is determined from the distance feature of linkage density and linkage accuracy in the linked-reads data (Fig. 2).

For sequence fragments derived from "difficult" regions such as segmental duplications, a barcode-agnostic aligner will place them at random locations (if multiple hits are found) and/or with low mapping quality scores. Linkage evidence from these ambiguously aligned fragments will be excluded based on the distance filter (100 kb) and by the mapping quality filter (see Extracting variant linkage information from long-range sequencing). The exclusion of ambiguous linkage evidence due to alignment inaccuracy ensures better accuracy of linkage evidence that is appropriate for **haplotype inference**. For *de novo* variant discovery, especially of **structural variants**, the benefit of improved alignment accuracy by barcode-aware alignment outweighs the compromise of linkage accuracy. For such applications, using barcode-aware alignment may be advantageous provided that the contributions to the mapping quality score from sequence alignment and from the molecular barcode can be accurately calibrated.

#### Haplotype inference and energy minimization of the 1D spin model

To better understand our strategy of haplotype inference, we introduce a different numerical representation of haplotype solutions using the parental haplotypes $\mathbf{H}_A$ and $\mathbf{H}_B = \overline{\mathbf{H}_A}$. We can represent the parental haplotype $\mathbf{H}_A$ as

$$h_{A,i} = \begin{cases} 1 & \text{if haplotype A has the reference genotype at site } i\text{ ,} \\ -1 & \text{if haplotype A has the alternate genotype at site } i\text{ .} \end{cases}$$

A numerical haplotype based on the reference/alternate representation $\mathbf{S}$ can be converted to $\mathbf{S}'$ in the paternal/maternal representation as

$$\mathbf{S} = \mathbf{S}' \cdot \mathbf{H}_A, \tag{S6}$$

where

$$s'_i = \begin{cases} 1 & \text{if the genotype at site } i \text{ agrees with haplotype A,} \\ -1 & \text{if the genotype at site } i \text{ agrees with haplotype B.} \end{cases}$$

Using this new representation, we can rewrite the coupling term in Eq. (10) as

$$M_{ij} s_i s_j = M_{ij} s'_i h_{A,i} s'_j h_{A,j} = M'_{ij} s'_i s'_j. \tag{S7}$$

with

$$M'_{ij} = M_{ij} h_{A,i} h_{A,j}. \tag{S8}$$

As $h_{B,i} = -h_{A,i}$, we also have

$$M'_{ij} = M_{ij} h_{B,i} h_{B,j}. \tag{S9}$$

It is straightforward to verify that $M'_{ij}$ is proportional to the difference between *cis* (A-A/B-B) linkage ($\mu_{ij}$) and *trans* (A-B/B-A) linkage ($\delta_{ij}$):

$$\begin{aligned} M'_{ij} &= M_{ij} h_{A,i} h_{A,j} = \chi_{ij} \sum_k \sigma_i^{(k)} \sigma_j^{(k)} h_{A,i} h_{A,j} \\ &= \chi_{ij} \Big[ \underbrace{\#(\sigma_i^{(k)} \sigma_j^{(k)} h_{A,i} h_{A,j} = 1)}_{\mu_{ij}} - \underbrace{\#(\sigma_i^{(k)} \sigma_j^{(k)} h_{A,i} h_{A,j} = -1)}_{\delta_{ij}} \Big] \end{aligned}$$

The energy function Eq. (10) now becomes

$$E'(\mathbf{S}') = -\frac{1}{2}\sum_{i,j}\chi_{ij}\left(\mu_{ij}-\delta_{ij}\right)s'_i s'_j. \tag{S10}$$

Finding the minimum of $E(\mathbf{S})$ is equivalent to finding $\mathbf{S}'$ that minimizes $E'(\mathbf{S}')$ as defined in Eq. (S10). If $\mu_{ij} > \delta_{ij}, \forall i, j$, it is straightforward to see that $E'(\mathbf{S}')$ has two global minima: $s'_i = 1$ or $s'_i = -1$. We next discuss how spin flipping and block switching can converge to these two minima starting from a random configuration

$$p(s'_{i,0} = 1) = p(s'_{i,0} = -1) = 1/2.$$

We first look at spin flips $s'_i \to -s'_i$. The associated energy changes are given by

$$\Delta E'_{i,0} = \sum_j \chi_{jk}\left(\mu_{ij}-\delta_{ij}\right)s'_{i,0}s'_{j,0} = s'_{i,0}h'_{i,0}. \quad \text{(spin flip)} \tag{S11}$$

By accepting spin flips that lower the energy function, we have

$$h'_{i,0} > 0 \to s_{i,1} = -1;$$
$$h'_{i,0} < 0 \to s_{i,1} = 1.$$

The probability that two sites $s'_{i,1}$ and $s'_{j,1}$ are phased correctly relative to each other is given by

$$p(s'_{i,1}\cdot s'_{j,1} = 1) = p(h'_{i,0}\cdot h'_{j,0} > 0).$$

We have

$$\begin{aligned} h'_{i,0}\cdot h'_{j,0} &= \sum_{k,l} M'_{ik}M'_{jl}s'_{k,0}s'_{l,0} \\ &= \sum_k M'_{ik}M'_{jk} + \sum_{k\neq l} M'_{ik}M'_{jl}s'_{k,0}s'_{l,0}. \end{aligned} \tag{S12}$$

For a random configuration, we have

$$p(s'_{k,0}\cdot s'_{l,0} = 1) = p(s'_{k,0}\cdot s'_{l,0} = -1) = 1/2. \tag{S13}$$

When $M'_{ij} \geq 0$, we have

$$\begin{aligned} E(h'_{i,0}\cdot h'_{j,0}) &= \sum_k M'_{ik}M'_{jk} + \sum_{k\neq l} M'_{ik}M'_{jl}E(s'_{k,0}\cdot s'_{l,0}) \\ &\approx \sum_k M'_{ik}M'_{jk} > 0, \end{aligned} \tag{S14}$$

which implies

$$p(s'_{i,1}\cdot s'_{j,1} = 1) = p(h'_{i,0}\cdot h'_{j,0} > 0) > p(h'_{i,0}\cdot h'_{j,0} < 0) = p(s'_{i,1}\cdot s'_{j,1} = -1).$$

In other words, more sites are phased correctly relative to each other after one round of spin flipping due to the positive coupling term. Equation (S14) further implies that as $p(s'_i\cdot s'_j = 1)\uparrow$ (*i.e.*, more sites are phased correctly),$E(s'_i\cdot s'_j)\uparrow$ and $E(h'_i\cdot h'_j)\uparrow$. Therefore, the iteration of spin flips can gradually improve phasing accuracy.

Although Eq. (S13) is true globally, it can be violated locally. An obvious example is at sites of long-range switching, *e.g.*,

$$s'_i = 1,\, i \leq k;\, s'_i = -1,\, i > k \Rightarrow s'_i s'_j = -1,\, i \leq k < j.$$

These errors are most efficiently removed by block switching moves that lower the energy by

$$\Delta E_{k|k+1} = \sum_{i\leq k}\sum_{j>k}\chi_{ij}\left(\mu_{ij}-\delta_{ij}\right)s'_i s'_j \approx -\sum_{i\leq k}\sum_{j>k}\chi_{ij}\left(\mu_{ij}-\delta_{ij}\right) < 0. \tag{S15}$$

Therefore, as long as the signal of true haplotype linkage is much stronger than background noise ($\mu_{ij} \gg \delta_{ij}$, $M'_{ij} > 0$), the iteration of spin flipping and block switching moves will converge **any initial state** to $\mathbf{S}' = 1$ or $\mathbf{S}' = -1$, and accordingly $\mathbf{S} \to H_A$ or $H_B$.

We can introduce more sophisticated minimization algorithms when the energy landscape contains many local minima. This happens when the linkage signal is sparse ($\mu_{ij} = 0$) or contains more errors $\delta_{ij} > \mu_{ij}$. Due to the presence of local minima, the final solution will depend on the initial haplotype configuration. We can use the same "steepest descent" strategy as described above but perform multiple simulations starting from different initial states to obtain a pool of haplotype solutions $\{\mathbf{S}^{(\alpha)}, \alpha = 1, 2, \cdots\}$ and then determine the haplotype linkage between site $i$ and $j$ from the pool of final states. For example, the haplotype linkage between site $i$ and $j$ in solution $\mathbf{S}^{(\alpha)}$ is given by $s_i^{(\alpha)} s_j^{(\alpha)}$, and its confidence can be estimated using

$$p_{i,j}^{(\alpha)} = \frac{1}{1 + e^{-\Delta E_{i,j}^{(\alpha)}}} \approx \begin{cases} 1 & \Delta E_{i,j}^{(\alpha)} \gg 0 \\ 1/2 & \Delta E_{i,j}^{(\alpha)} \sim 0 \\ 0 & \Delta E_{i,j}^{(\alpha)} \ll 0 \end{cases}$$

where

$$\Delta E_{i,j}^{(\alpha)} = \min\left(\Delta E_i^{(\alpha)}, \Delta E_j^{(\alpha)}, \Delta E_{i|j}^{(\alpha)}\right)$$

measures the energy changes of flipping ($\Delta E_i$, $\Delta E_j$) and switching ($\Delta E_{i|j}$) perturbations. We can infer the optimal haplotype linkage $\overline{s_i s_j}$ from the likelihood ratio that is similar to Eq. (1):

$$\frac{p(\overline{s_i s_j} = 1)}{p(\overline{s_i s_j} = -1)} = \prod_{s_i^{(\alpha)} s_j^{(\alpha)} = 1} \frac{p_{ij}^{(\alpha)}}{1 - p_{ij}^{(\alpha)}} \prod_{s_i^{(\beta)} s_j^{(\beta)} = -1} \frac{1 - p_{ij}^{(\beta)}}{p_{ij}^{(\beta)}}. \tag{S16}$$

To enable more efficient sampling of the configuration space in each simulation, one can also use the Metropolis algorithm for spin flipping and block switching moves.

### Determination of the K-562 karyotype by haplotype-specific genomic analysis

In this subsection we discuss results presented in the Additional Data Document ("**Complete karyotype analysis of the K-562 genome using Hi-C**"). All genomic coordinates are according to the GRCh38 human genome reference.

#### Cytogenetic karyotypes

Pages 2 and 3 show the cytogenetic karyotypes of K-562 cells (page 2) and selective marker chromosomes (page 3) analyzed using MFISH by Gribble *et* al.(2000) (Ref.39) and Naumann *et* al.(2001) (Ref.40).

#### Chromosomal copy number

Pages 5-7 show the normalized DNA copy number of each parental homolog (100kb bins) calculated using allelic depths calculated from the linked-reads data in combination with parental haplotypes determined using the linked-reads data and the Hi-C data. Chromosomes 3,9,13,14 and X show complete loss-of-heterozygosity and the DNA copy number is calculated using the total sequencing depth. Segmental deletion/loss-of-heterozygosity affects Chr.2A (blue, q-terminus), Chr.10A (red, q-terminus), Chr.12A (blue, p-terminus), Chr.17B (blue, p-arm), Chr.20A (blue, p-arm) and Chr.22A(red, q-terminus). These regions are omitted in the allelic copy-number plots. We also assign large segmental copy-number alterations to derivative chromosomes that are labelled on top of the copy-number plots. The syntenic structures of these derivative chromosomes are discussed next. The only incompletely resolved segmental copy-number changes are gains of Chr.7q (blue homolog) and Chr.9q (outlined with magenta boxes).

#### Digital karyotype

Pages 9 and 10 summarize the haplotype-resolved syntenic structures of normal (page 5) and marker/derivative chromosomes (page 6). Among the structurally normal chromosomes, both copies of Chr.4B contain a deletion between 159,570,188 and 162,695,549 (annotated); Chr.16q contains a tandem duplication of DNA sequence between 88,525,011 and 88,794,607 that is inferred to affect the B homolog by DNA copy number (not annotated). The only major discrepancy between the sequencing-derived karyotype and the cytogenetic karyotypes is seen in Chr.7. Both Gribble *et al.* (2000) and Naumann *et al.* (2001) reported a single normal Chr.7 plus three rearranged (marker) Chr.7. Naumann *et al.* inferred that one Chr.7 marker chromosome (M4) contained a paracentric inversion of p11-p22 and two Chr.7 marker chromosomes (M5 and M6) had rearrangements of both p- and q-arms resulting in a net gain on Chr.7q. Our DNA copy number analysis reveals a segmental gain of Chr.7q from ~117Mb to the q-terminus but we cannot detect any rearrangement either near the copy-number breakpoint or elsewhere on

Chr.7 from either linked-reads or Hi-C data. We think the normal Chr.7 in Ref.40 is the A homolog and the K-562 cells used to generate the linked-reads data (from which the DNA copy number is calculated) had gained an extra copy of Chr.7A. The marker chromosome (M4) with the paracentric inversion may have been lost. The two markers M5 and M6 are both derived from the B homolog, but the fusion breakpoints are likely located in heterochromatic regions (centromere/telomere) that cannot be resolved by shotgun sequencing. Among the marker chromosomes, the composition of t(22A;9-13-22hsr) is inferred using both sequencing and cytogenetic data (Page 3) and should be taken as a model.

**Marker chromosomes with completely resolved translocations**

Pages 11-16 summarize the data and analysis that completely determines all the syntenic blocks and junctions in five marker chromosomes: t(5A;6A), t(12A;21A), t(3A;10A), t(3A;10A;17A), and t(6A;16A;6B). All of these marker chromosomes were also identified in Refs. 39 and/or 40. For each chromosome, we first identify the syntenic blocks from the haplotype-specific DNA copy number and Hi-C contacts; we then refine the junction breakpoints first using the barcode map generated from linked-reads data and then by sequencing reads with split alignments. Each phased Hi-C contact map contains 9 panels: the bottom left panel shows the density map of unphased Hi-C contacts; the upper left and bottom right panels show the density maps of Hi-C contacts where one end of the contact is phased (A-U, B-U, U-A, and U-B); the upper right panels are scatter plots of all Hi-C contacts with both ends phased (A-A, A-B, B-A, and B-B). The density map of molecular barcodes shows the number of unique molecular barcodes with sequencing fragments mapping to loci on both axes. A fusion between distal loci results in a sharp increase in the density of off-diagonal contacts in both Hi-C and molecular barcode maps; the breakpoints are annotated using red lines meeting at the apex with the highest density of contacts/barcodes. The barcode density map is generated with 10kb bins, which on average contains about 100 unique molecular barcodes per chromosome. We can therefore infer the copy number of the translocated chromosome. For example, t(5A;6A), t(12A;21A), and t(6A;16A;6B) all have one copy per genome; the junction t(3A;10A) has two copies per genome, although one copy has an additional translocation to Chr.17 to make t(3A;10A;17A).

There are a few notable examples. In t(3A;10A;17A), we infer the fusion between Chr.10A and Chr.17A is at the centromeres; we have omitted the molecular barcode map as the junction is not represented in the human genome reference and therefore cannot be resolved by shotgun sequencing. t(6A;16A;6B) is a special example as it involves both Chr.6 homologs. The resolution of the structure of this chromosome is possible because the parental Chr.6 haplotype can be determined from Hi-C contacts within the normal Chr.6B, even though there is no normal Chr.6A. The proximity of breakpoints on Chr.6A at 37,856,135 in t(5A;6A) and at 38,139,520 in in t(6A;16A;6B) suggests that these two breakpoints/translocations may have resulted from a single DNA break on Chr.6A near 38Mb.

**Marker chromosomes with incompletely resolved translocations**

Pages 17-23 summarize the analysis of derivative/marker chromosomes with one or more junctions not completely resolved. Nost of the incompletely resolved junctions contain DNA sequence with ambiguous origin (most likely derived from heterochromatic regions).

We infer t(2A;22A) (Page 18) to contain two inverted duplication (“foldback”) events on Chr.2 followed by a translocation to Chr.22. This chromosome corresponds to M1 in Ref. 40. Naumann *et al.* reported that the added Chr.22 segment also contains Chr.9 material that forms the *BCR-ABL* fusion. Consistent with this result, we observe an increase in Hi-C contact between Chr.2 and Chr.9 (data not shown) but cannot determine the telomeric end of this chromosome.

We infer t(1B;21B) (Page 19) to be generated by a translocation between Chr.1B at 162,379,862 and the acrocentric arm of Chr.21B, but cannot determine the breakpoint location on Chr.21.

We determine the composition of t(6A;1A;20A) (Pages 20 and 21) from the Hi-C maps and further detect a small piece from Chr.18 inserted between Chr.6A and Chr.1A (Page 22). For the Chr.1:Chr.18 junction, both breakpoints are resolved by sequencing reads. For the Chr.18:Chr.6 junction, the breakpoint on Chr.6 is almost certainly located at 135,580,256 and the breakpoint on Chr.18 is likely located at 27,330,533, but we cannot assemble the complete junction sequence as these breakpoints are joined by DNA sequence with ambiguous origin (likely derived from heterochromatic regions). We are unable to detect an increase in the molecular barcode density between Chr.6 and Chr.18 breakpoints in the linked reads data, indicating that the inserted sequence may be longer than 100kb.

Besides t(6A;1A;20A), we infer both Chr.18 homologs to be involved in complex rearrangements giving rise to t(1A;18A) and t(3A;18B) (Pages 22-26). Page 22 shows the density map of unphased Hi-C contacts between Chr.18 and Chr.1 and between Chr.18 and Chr.3. The breakpoints in t(1A;18A) are resolved to be Chr.1:54,726,344 and Chr.18:25,907,722 (Page 23). The breakpoint on Chr.3 in t(3;18A) is resolved to be Chr.3:138,669,875 (Page 24) but it is joined with DNA sequence with ambiguous origin. The partner breakpoint on Chr.18 cannot be determined with certainty. Interestingly, both the Chr.1A junction and the Chr.3A junction contain inverted duplications (opposite facing arrows) near the breakpoint.

To completely resolve the alterations to both Chr.18 homologs, we jointly analyze rearrangement breakpoints and haplotype-specific DNA copy number of Chr.18 (Page 26). The Chr.18A homolog contains a foldback rearrangement at 42Mb and 3 other breakpoints (24,413,290, 24,822,840, and 41,843,305) whose fusion partners cannot be identified. An inspection of the Hi-C contact map between Chr.1 and Chr.18 reveals that the foldback rearrangement at 42Mb is part of t(1A;18A). Given that there are two p-arms of Chr.18A but only one normal q-terminus of Chr.18A, we infer that there is one normal Chr.18A and the other Chr.18A p-arm is connected to the rearranged segments on Chr.18q and then joins Chr.1A to form a stable chromosome with two telomeres.

We identify 18 breakpoints on Chr.18B. A close inspection of the Hi-C contact map between Chr.3 and Chr.18 (Page 24) reveals that the rearranged Chr.18B is connected to Chr.3. The fusion between breakpoints at 39,768,382 and 40,287,508 results in a tandem duplication and appears to be unrelated to the others. Among the remaining 16 breakpoints on Chr.18B, 12 are paired and result in little or no DNA copy number change; the breakpoint at 26.97Mb is a foldback rearrangement; three breakpoints (10,860,074, 21,920,738, and 27,509,642) are accompanied with DNA copy-number changes. 14 breakpoints are connected by rearrangements (a-e on Page 25) detected from linked reads and further validated on the Hi-C contact map. The junction between Chr.18:8,110,157 and 26,776,296 is not visible on the Hi-C map. It is evident from the barcode map that this junction is linked to another breakpoint near 26,776,296, which is identified to be located at 26,770,719 but with an unidentifiable fusion partner. From the Hi-C contact map we infer the breakpoint at 8,110,157 eventually joins the breakpoint at 27,509,642 that also has an unidentified partner. Since the only other breakpoint on the B homolog (blue) with an unidentified partner is at 23,730,830, we infer that Chr.18:23,730,830 is connected to Chr.3A at 138,669,875 with additional sequence insertion.

Chromosome 9 has by far the most alterations and was inferred to participate in four marker chromosomes (M7, M8, M15 and M16 in Ref.40) in addition to the *BCR-ABL* amplicon present in t(2A;22A) and t(22;hsr). We are able to partially resolve three marker chromosomes involving Chr.9 (Pages 27-31).

We first infer that three Chr.9 fragments are partitioned to separate marker chromosomes from the Hi-C map (Page 27). The p-terminal segment (0-20.75Mb) joins Chr.17q (B homolog) at the centromere, as is determined from the Hi-C contact map between Chr.9 and Chr.17 (Page 28). The t(9;17B) chromosome is present at two copies and match M16 in Ref. 40.

The two segments Chr.9:26.58-28.55Mb and Chr.9:31.62-38.43Mb show few contacts with the rest of Chr.9 (Page 27) but form extensive contact throughout Chr.13q (Page 29), suggesting a derivative chromosome t(9;13). The only copy-number breakpoint with an unidentified fusion partner is Chr.9:32,239,218, which most likely joins Chr.13 (Page 30). The t(9;13) matches M15 reported in Ref. 40.

The remaining Hi-C contacts are consistent with a marker Chr.9 with deletion from the p-terminus to ~37Mb. We additionally identify a foldback rearrangement at 37.08Mb that marks the distal end of this chromosome. This chromosome matches M8 reported in Ref.40 that is annotated as del(9)(:p12-qter). We also detect a gained segment on Chr.9q from 91,321,259 to the q-terminus but cannot map the fusion partner of the breakpoint. Naumann *et al.* reported that this segment is fused to the q-terminus of Chr.9 and forms M7.

Finally, we attempt to infer the structure of the amplicon in the t(9;13;22) chromosome that contains multiple copies of the *BCR-ABL* fusion gene. We start with the amplified segment on Chr.9. This segment is flanked by three breakpoints: 130,731,760 at the 5'-end (fusion 1), 131,199,197 (fusion 2) and 131,280,137 (fusion 3) on the 3'-end.

The breakpoint at Chr.9:131,280,137 joins Chr.13:108,009,063 (fusion 3) and is connected to multiple amplified segments on Chr.13q (Pages 31 and 32). The amplicons on Chr.13q have roughly the same DNA copy number ~12 (Page 33); the rearrangements further suggest that the individual segments are joined together in a single chunk and then amplified as a single unit. Because there is only one junction connecting the Chr.13 fragments to Chr.9, we infer that the Chr.13 fragments form a linear palindromic structure with a foldback junction at Chr.13:91,821,757><91,823,504 and both ends of the palindrome join Chr.9 through fusion 3.

The breakpoint at Chr.9:131,199,197 joins Chr.22:16,819,349 (fusion 2) and its copy number is estimated to be ~8 (Page 33); the breakpoint at Chr.9:130,731,760 joins Chr.22:23,290,555 and creates the *BCR-ABL* fusion gene (fusion 1) with an estimated DNA copy number ~20. In contrast to Chr.9 and Chr.13, Chr.22 shows multiple copy-number states that are consistent with breakage-fusion-bridge cycles interspersed with chromothripsis. We further identify two rearrangement junctions on Chr.22 with similar DNA copy number as fusion 2. One event is a fusion between Chr.22:18,965,610 and Chr.22:22,592,903, the other is a foldback rearrangement at chr22:16,243,545-16,245,357. The other copy-number breakpoints on Chr.22 all have less DNA copy number and therefore most likely occurred after the initial amplification. Putting all the information together, we infer that the Chr.9:Chr.22 amplicon is generated from four segments: Chr.22:16.24-16.81Mb, Chr.9:130.73-131.20Mb, Chr.22:22.59-23.59Mb, and Chr.22:18.96-23.29Mb; these segments are joined in tandem and then form a palindrome with a foldback junction at Chr22:16,243,545><16,245,357. The ends of the palindrome join Chr.9 through fusion 1.

We further infer that the 9:13 amplicon and the 9:22 amplicon are joined together in a single amplicon that is amplified in an inverted tandem array structure. The tandem array configuration is consistent with MFISH analysis in Ref. 39 indicating that the 9:13:22 amplicon creates a homologous staining region (HSR) in two marker chromosomes each containing 4-5 copies of the Chr.13 probe (Page 32). We speculate that the slightly higher DNA copy number of the 9:13 amplicon (~12) than the 9:22 amplicon (~8) is due to extra copies of the 9:13 amplicon in the t(2A;22A) derivative chromosome. The basic unit of the tandem array is a palindrome formed by inverted duplication of a single 9:13:22 amplicon; the palindrome then undergoes multiple rounds of duplications in tandem. Interestingly, the tandem array only consists of foldback rearrangements but all DNA sequence is amplified uniformly. This contrasts with amplification by the breakage-fusion-bridge cycles that are also generated by foldback rearrangements but usually result in varying DNA copy number. We think the foldback rearrangements in the 9:13:22 HSR were first generated by breakage-fusion-bridge cycles, but the creation of almost perfect palindromic sequences (Chr.13:91,821,757><91,823,504 and Chr22:16,243,545><16,245,357) leads to more frequent amplifications by homology-dependent DNA recombination or replication errors. We speculate that the formation of palindromic sequence by foldback rearrangements between breakpoints in close proximity may be a hallmark features of HSRs.

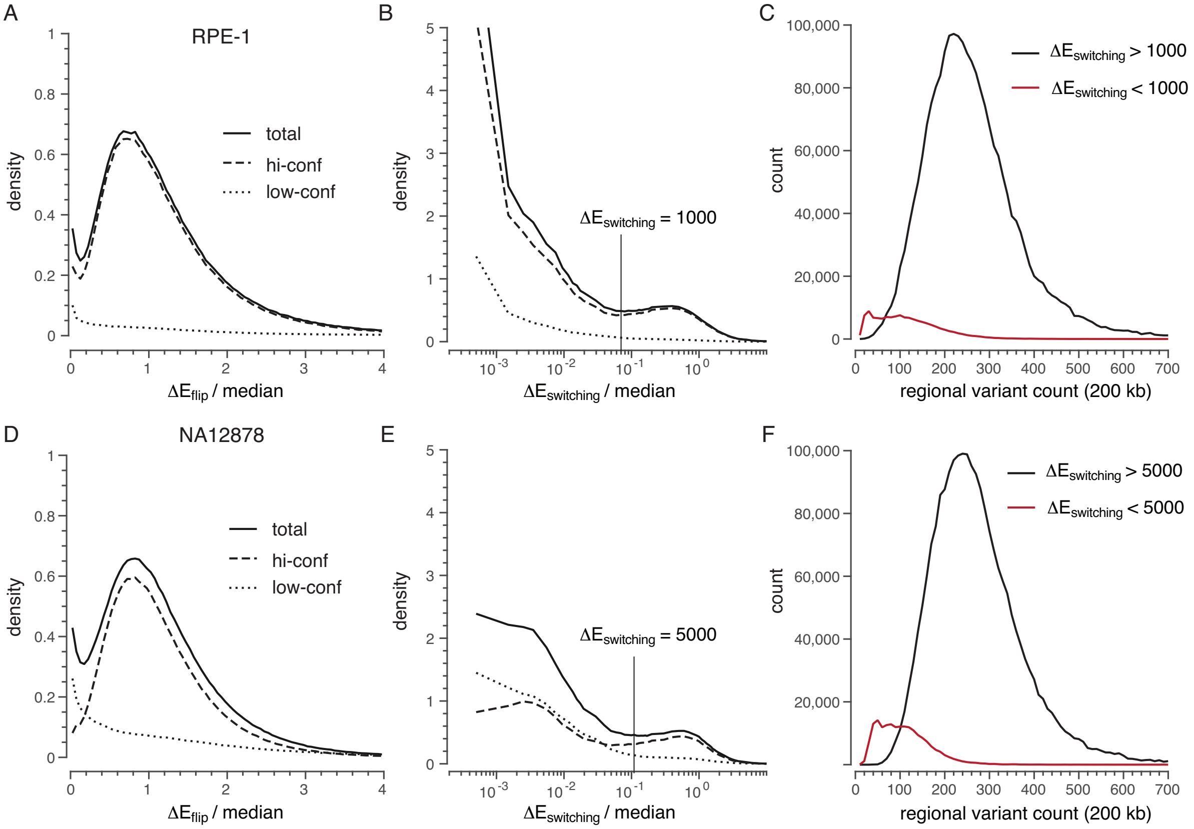


Figure S1: Distributions of the energy penalty scores in the haplotype solution of the RPE-1 (A-C) and NA12878 (D-F) linked-reads data. The energy penalty scores measure the confidence of haplotype inference with respect to single base (A and D) and block-switching (B and E) errors. For each type of phasing error, the distribution of penalty scores is shown for all variants (solid line) and separately for high-confidence variants (dashed line) and low-confidence variants (dotted line). In the distribution plots, the energy penalty is normalized by the median number of unique molecular links at each variant site. Panel C and F show the distributions of local variant density near each variant site (estimated by the number of high-confidence variants within a 200 kb neighborhood) with high (black) and low (dark red) switching penalty scores in the RPE-1 genome (C) and in the NA12878 genome (F).

Table 1: Data used for haplotype inference and karyotype reconstruction of the K-562 genome

| Sample | Data type | Data source | Read count | Mean depth | Application |
|---|---|---|---|---|---|
| K-562 | linked reads | ENCODE[a] | 736,246,361[b] | 60×[c] | local phasing |
| K-562 | Hi-C | Rao et al. (2014) (Rao et al., 2014) | 456,757,799[d]<br>591,854,553[e] | | long-range phasing,<br>karyotype construction |

[a] `https://www.encodeproject.org/experiments/ENCSR053AXS/`

[b] Mean Molecular length 58.2 kb; median insert 385; 661,037,851 aligned in pair; 151 × 2; duplication rate 0.012.

[c] excluding the GEMcode sequence

[d] SRR1658693: median insert 357; 428,963,615 aligned in pair; 101 x 2; duplication rate 0.05.

[e] SRR1658694: median insert 341; 578,201,812 aligned in pair; 101 x 2; duplication rate 0.17.

Table 2: Comparison between the scaffold haplotype solution and the reference haplotypes of NA12878

| ref_id | Total variant sites[a] | Phased from parents[b] | Sites for phasing[c] | Phased from bulk data[d] | Comparable sites | Fraction of agreement | Fraction of completion[e] |
|---|---|---|---|---|---|---|---|
| chr1 | 193,281 | 145,417 | 176,458 | 160,578 | 137,172 | 0.995 | 0.939 |
| chr2 | 208,196 | 147,477 | 189,810 | 174,664 | 139,562 | 0.999 | 0.946 |
| chr3 | 175,998 | 136,151 | 155,662 | 145,153 | 126,777 | 0.999 | 0.930 |
| chr4 | 179,803 | 115,678 | 163,757 | 153,465 | 110,687 | 0.999 | 0.957 |
| chr5 | 164.837 | 120,895 | 151,544 | 142,076 | 115,291 | 0.999 | 0.953 |
| chr6 | 174,820 | 132,898 | 161,825 | 152,261 | 126,250 | 0.986 | 0.937 |
| chr7 | 147,495 | 108,363 | 136,008 | 124,499 | 102,500 | 0.996 | 0.942 |
| chr8 | 136,150 | 95,067 | 126,479 | 117,763 | 90,290 | 0.999 | 0.949 |
| chr9 | 119,635 | 84,264 | 109,185 | 99,223 | 80,150 | 0.999 | 0.951 |
| chr10 | 133,655 | 97,420 | 119,667 | 111,518 | 91,867 | 0.992 | 0.935 |
| chr11 | 125,208 | 94,319 | 111,748 | 103,244 | 87,692 | 0.999 | 0.929 |
| chr12 | 120,828 | 85,751 | 110,074 | 102,101 | 80,984 | 0.999 | 0.944 |
| chr13 | 89,785 | 71,352 | 83,292 | 78,981 | 69,059 | 0.999 | 0.968 |
| chr14 | 81,921 | 63,767 | 76,081 | 70,468 | 60,787 | 0.999 | 0.952 |
| chr15 | 72,770 | 51,991 | 64,540 | 58,841 | 49,344 | 0.999 | 0.948 |
| chr16 | 80,402 | 36,736 | 73,735 | 67,472 | 35,018 | 0.994 | 0.947 |
| chr17 | 68,924 | 45,706 | 61,140 | 54,597 | 42,054 | 0.989 | 0.910 |
| chr18 | 73,239 | 44,077 | 66,670 | 62,328 | 41,705 | 0.999 | 0.946 |
| chr19 | 63,856 | 39,204 | 46,748 | 41,658 | 30,635 | 0.985 | 0.769[f] |
| chr20 | 68,144 | 42,736 | 53,616 | 49,585 | 40,104 | 0.998 | 0.937 |
| chr21 | 44,623 | 26,893 | 32,979 | 31,183 | 25,818 | 0.999 | 0.960 |
| chr22 | 44,187 | 24,229 | 33,498 | 30,971 | 23,024 | 0.990 | 0.941 |
| chrX | 84,624 | 51,550 | 74,634 | 61,529 | 45,244 | 0.999 | 0.877 |
| Total | 2,652,381 | 1,861,941 | 2,379,150 | 2,194,158 | 1,746,304 | 0.997 | 0.938 |

[a]single-nucleotide variant sites with one reference and one alternate genotypes detected by HaplotypeCaller without any filter

[b]phased variants in the Genome-In-A-Bottle release

[c]excluding sites in centromeric regions or not having linkage to both reference and alternate genotypes

[d]scaffold haplotype solution

[e]percentage of correctly phased sites

[f]The low percentage is because a large number of variants in the trio haplotype are located in the centromere region and excluded from the scaffold haplotype.

Table 3: Comparison between the scaffold haplotype and reference haplotypes of RPE-1

| ref_id | Total variant sites[a] | Phased from monosomies[b] | Sites for phasing[c] | Phased from bulk data[d] | Comparable sites | Agreed | Fraction of agreement |
|---|---|---|---|---|---|---|---|
| chr1 | 185,997 | 179,216 | 176,078 | 163,057 | 159,408 | 157,056 | 0.985 |
| chr2 | 194,641 | 188,297 | 186,201 | 175,383 | 172,093 | 170,210 | 0.989 |
| chr3 | 169,194 | 157,887 | 152,824 | 145,109 | 137,854 | 135,905 | 0.986 |
| chr4 | 168,626 | 161,562 | 161,915 | 154,564 | 150,795 | 148,588 | 0.985 |
| chr5 | 149,212 | 146,447 | 142,975 | 135,587 | 134,138 | 132,877 | 0.991 |
| chr6 | 156,723 | 155,379 | 149,049 | 143,371 | 142,649 | 141,478 | 0.992 |
| chr7 | 143,405 | 139,365 | 136,858 | 129,164 | 126,868 | 123,263 | 0.972 |
| chr8 | 124,489 | 120,140 | 120,905 | 114,559 | 111,458 | 110,321 | 0.990 |
| chr9 | 115,189 | 110,993 | 108,636 | 99,516 | 97,845 | 94,032 | 0.961 |
| chr10 | 119,524 | 112,758 | 112,078 | 106,059 | 102,486 | 100,767 | 0.983 |
| chr11 | 116,321 | 113,375 | 107,966 | 102,255 | 101,091 | 98,782 | 0.977 |
| chr12 | 111,225 | 106,331 | 105,314 | 99,033 | 96,115 | 94,095 | 0.979 |
| chr13 | 84,586 | 83,092 | 82,270 | 78,835 | 77,752 | 77,059 | 0.991 |
| chr14 | 74,463 | 71,195 | 71,637 | 67,289 | 65,023 | 64,021 | 0.985 |
| chr15 | 70,936 | 62,662 | 66,134 | 61,948 | 55,646 | 54,013 | 0.971 |
| chr16 | 78,049 | 70,375 | 72,948 | 67,658 | 62,217 | 61,154 | 0.983 |
| chr17 | 75,407 | 60,418 | 69,001 | 64,261 | 52,385 | 50,393 | 0.962 |
| chr18 | 64,006 | 61,622 | 60,708 | 57,620 | 57,037 | 56,692 | 0.994 |
| chr19 | 61,461 | 26,906[e] | 44,083 | 41,246 | 18,402 | 18,023 | 0.979 |
| chr20 | 61,029 | 55,063 | 49,742 | 46,735 | 43,738 | 42,999 | 0.983 |
| chr21 | 44,178 | 36,240 | 33,089 | 31,755 | 28,079 | 27,885 | 0.993 |
| chr22 | 40,635 | 36,154 | 30,115 | 28,140 | 26,142 | 25,802 | 0.987 |
| chrX | 66,015 | 64,676 | 62,476 | 52,503 | 51,926 | 51,441 | 0.991 |
| Total | 2,475,311 | 2,320,153 | 2,303,002 | 2,165,647 | 2,071,147 | 2,036,856 | 0.983 |

[a]single-nucleotide variant sites with one reference and one alternate genotypes detected by HaplotypeCaller without any filter
[b]whole-chromosome haplotypes determined from single-cell sequencing of monosomic RPE-1 cells
[c]excluding sites in centromeric regions or not having linkage to both reference and alternate genotypes
[d]scaffold haplotype solution
[e]generated from one monosomic cell and having a lower percentage of phased variants

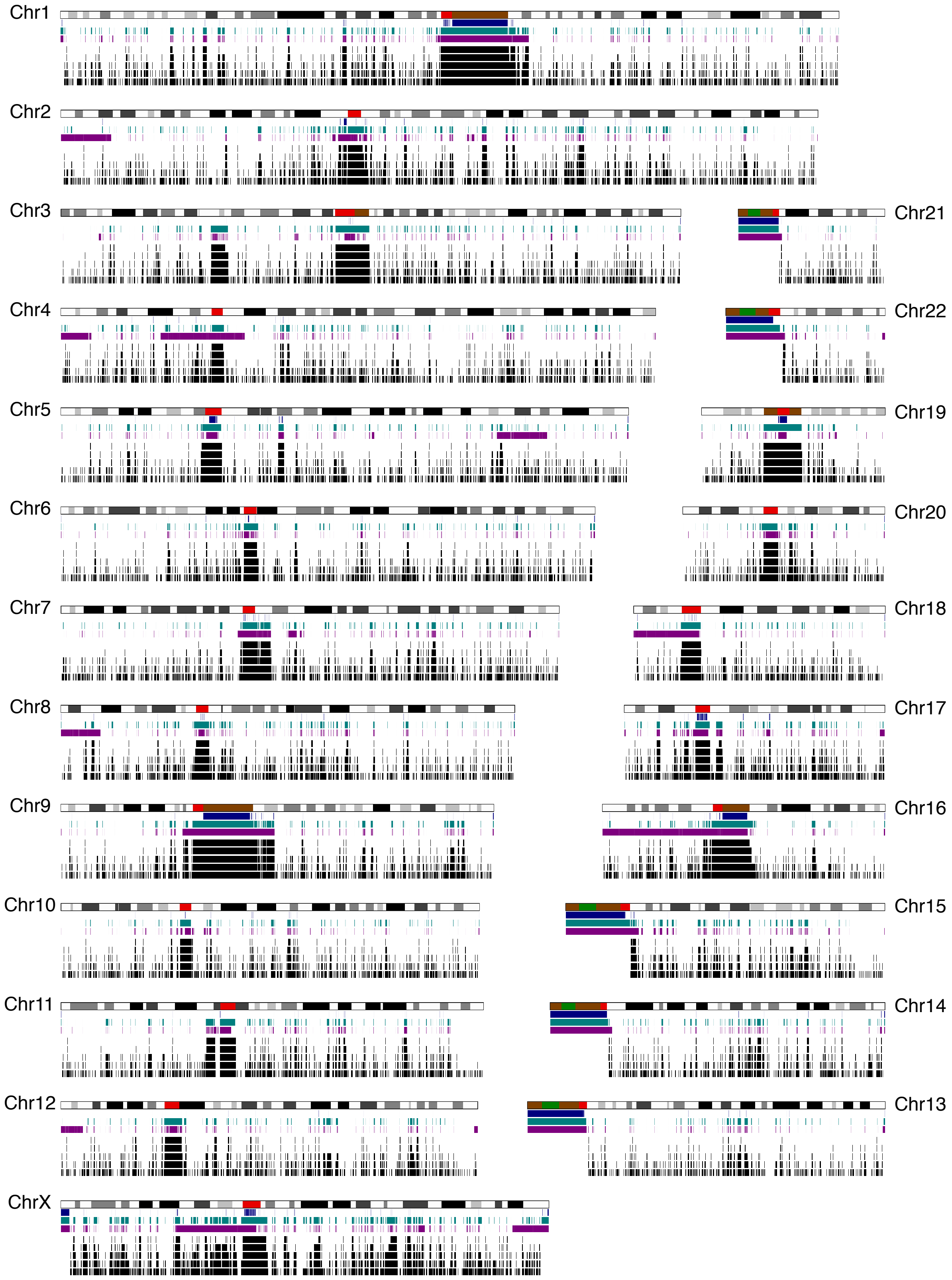


Figure S2: Low variant density regions in the NA12878 genome and boundaries of haplotype blocks inferred from the linked-reads data. The variant density tracks follow the same definitions as given in Fig. 4. The haplotype blocks are determined based on the following block-switching penalty cutoff (top to bottom): 5,20,100,500,5000. The last block-switching cutoff are used to generate high-confidence haplotype blocks in the scaffold haplotype.

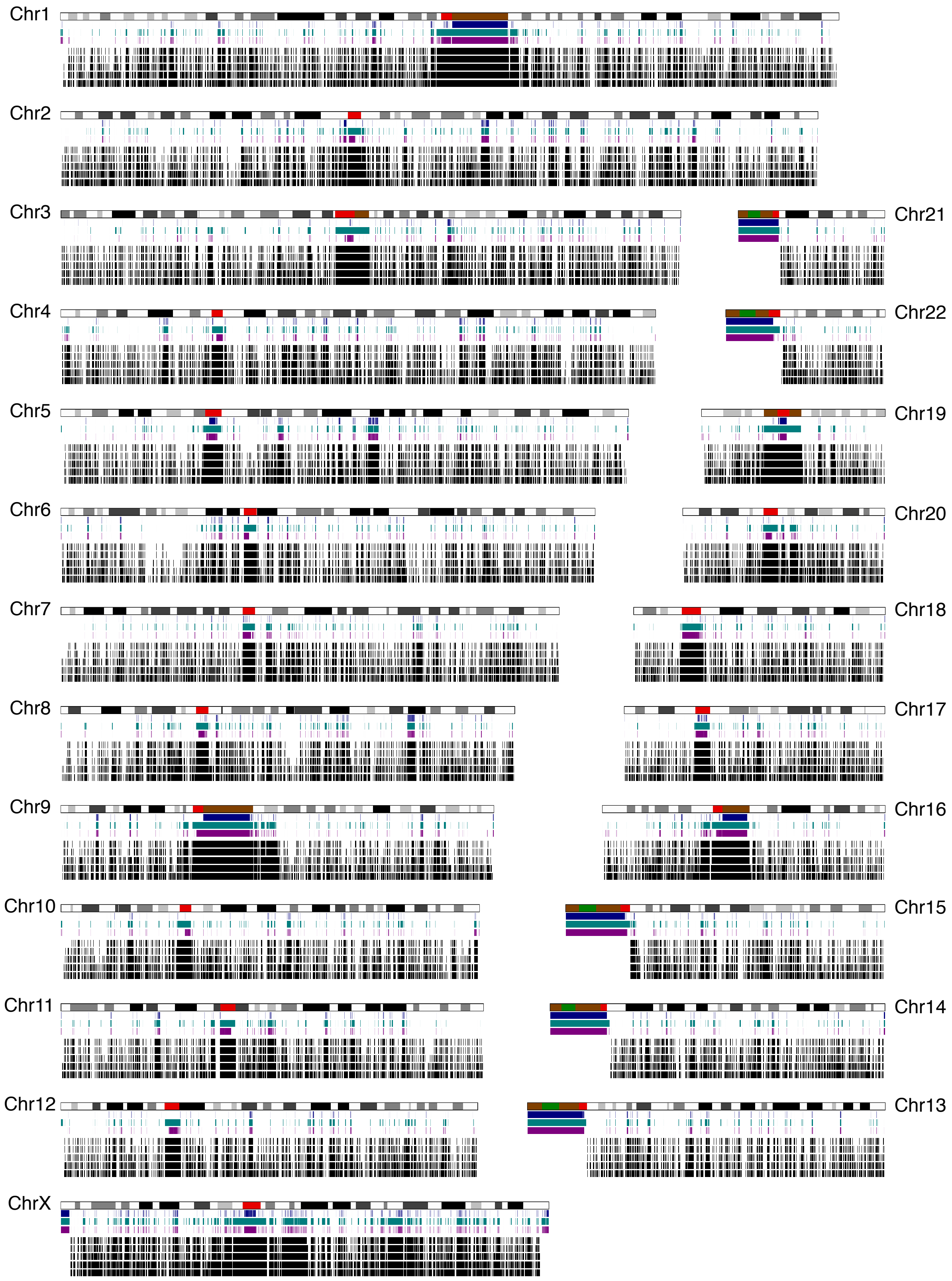


Figure S3: Low variant density regions in the RPE-1 genome and boundaries of haplotype blocks inferred from the linked-reads data. The haplotype blocks are determined based on the following block-switching penalty cutoff (top to bottom): 5,20,100,500,1000. The last block-switching cutoff are used to generate high-confidence haplotype blocks in the scaffold haplotype.

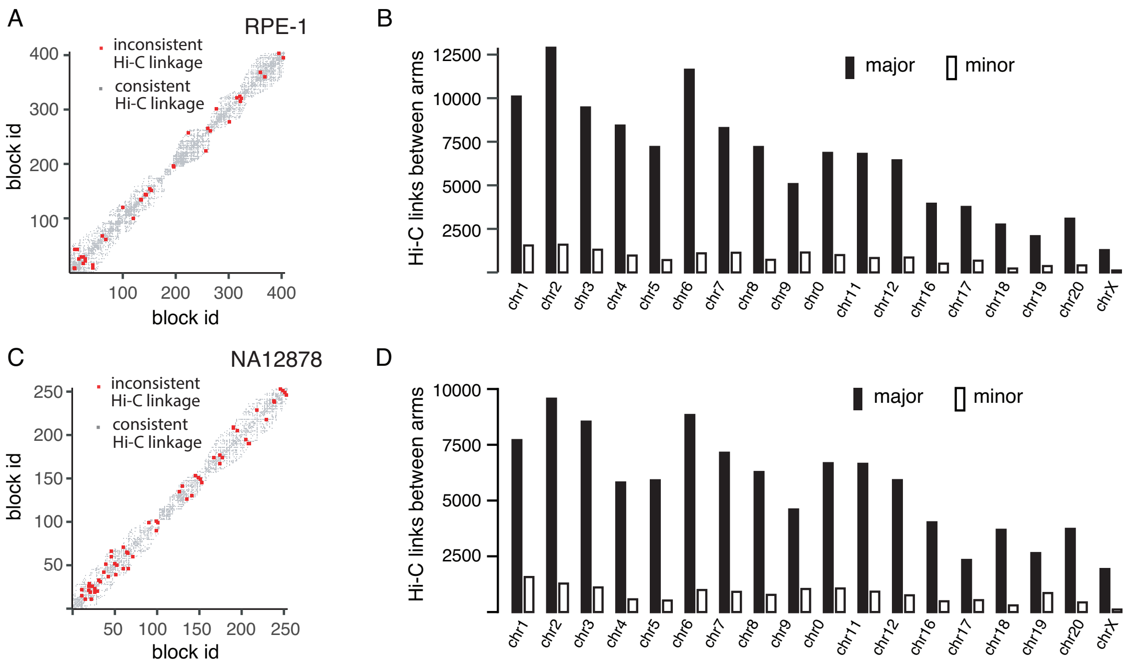


Figure S4: Concatenation of haplotype blocks using Hi-C links. A and C. Maps of *cis* (gray) and *trans* (red) Hi-C linkage between local haplotype blocks in the scaffold haplotype solution of Chr.X determined from the RPE-1 (A) and the NA12878 data (C). The centromere is located near block 200 in the RPE-1 solution and near block 100 in the NA12878 solution. B and D. Total number of *cis* and *trans* Hi-C links between p- and q-arms in the RPE-1 (B) and the NA12878 data (D).
