## Supplementary figures related to the analysis of K-562 karyotype. for "Determination of complete chromosomal haplotypes by bulk DNA sequencing"

### K-562: cytogenetic karyotypes

Gribble et al. Cancer Genetics and Cytogenetics, 2000.

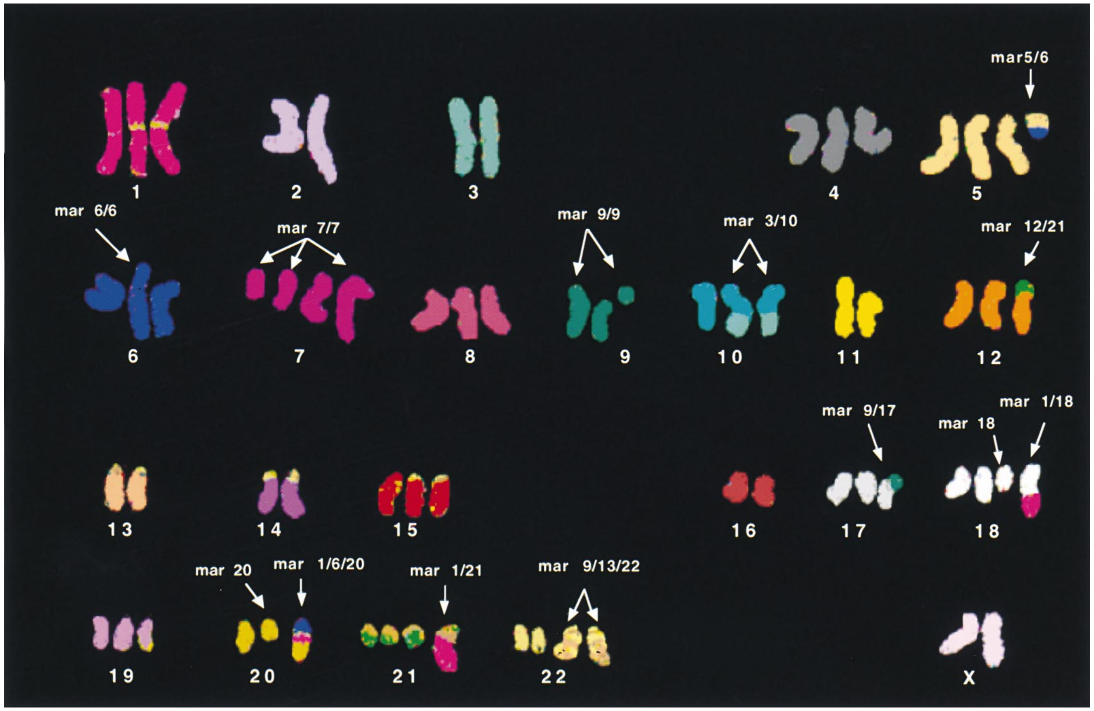


Naumann et al. Leukemia Research, 2001.

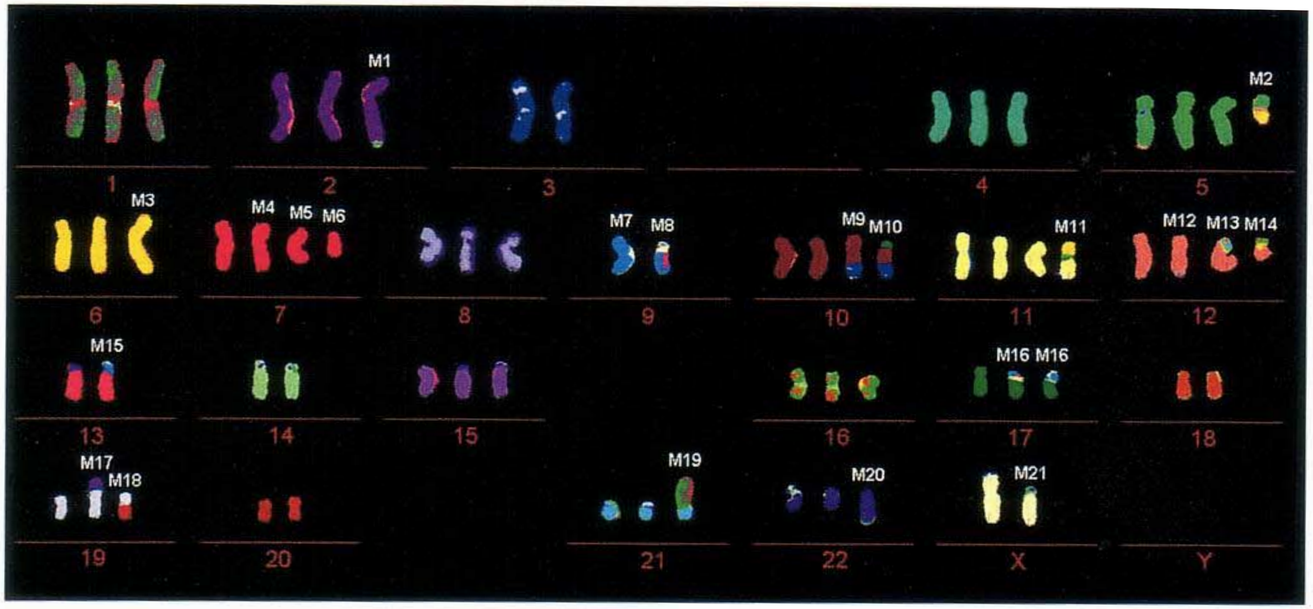

### Selected marker chromosomes

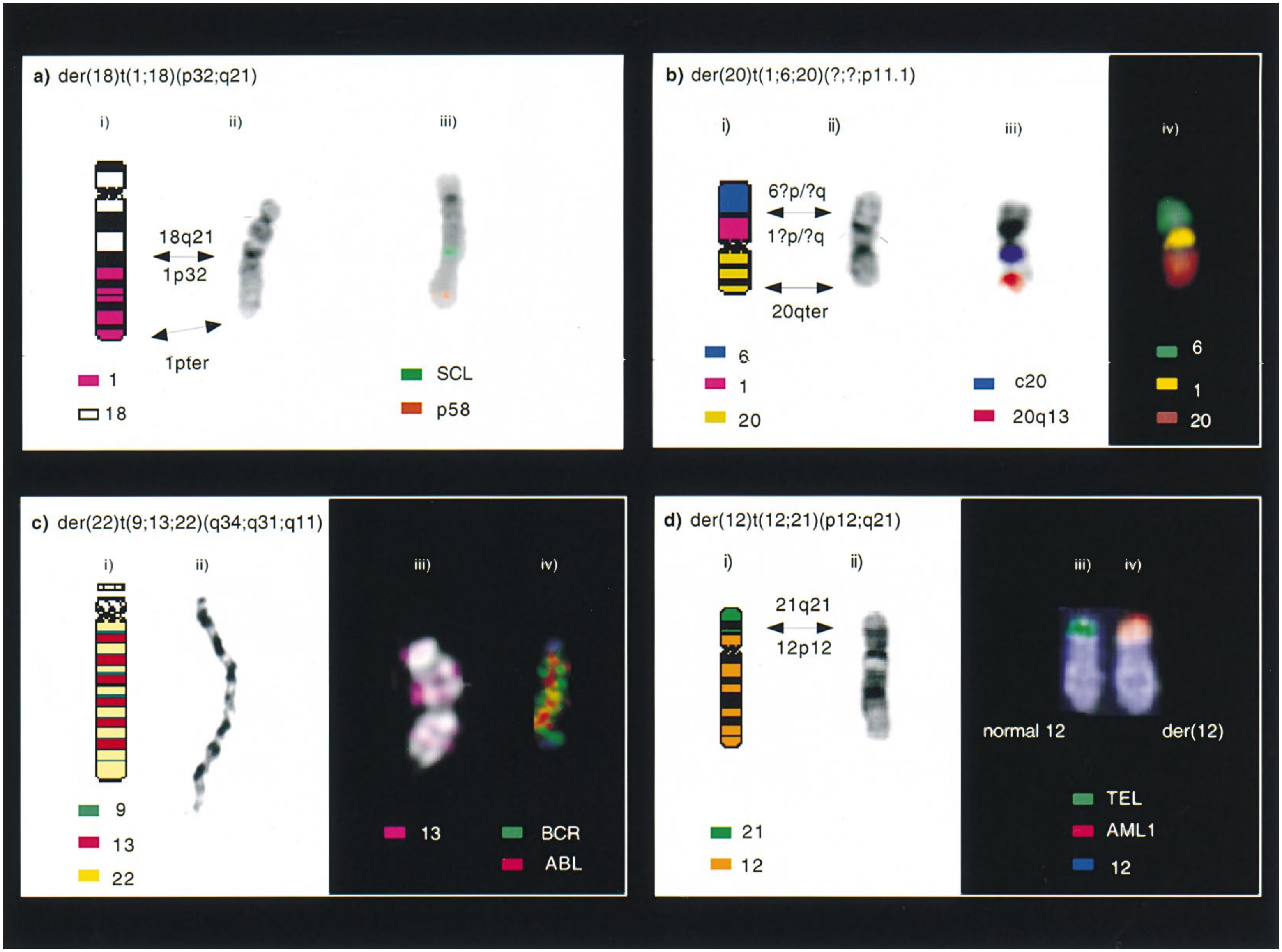


**Gribble et al. Cancer Genetics and Cytogenetics, 2000.**

### Chromosomal copy number

### Chromosomal copy number

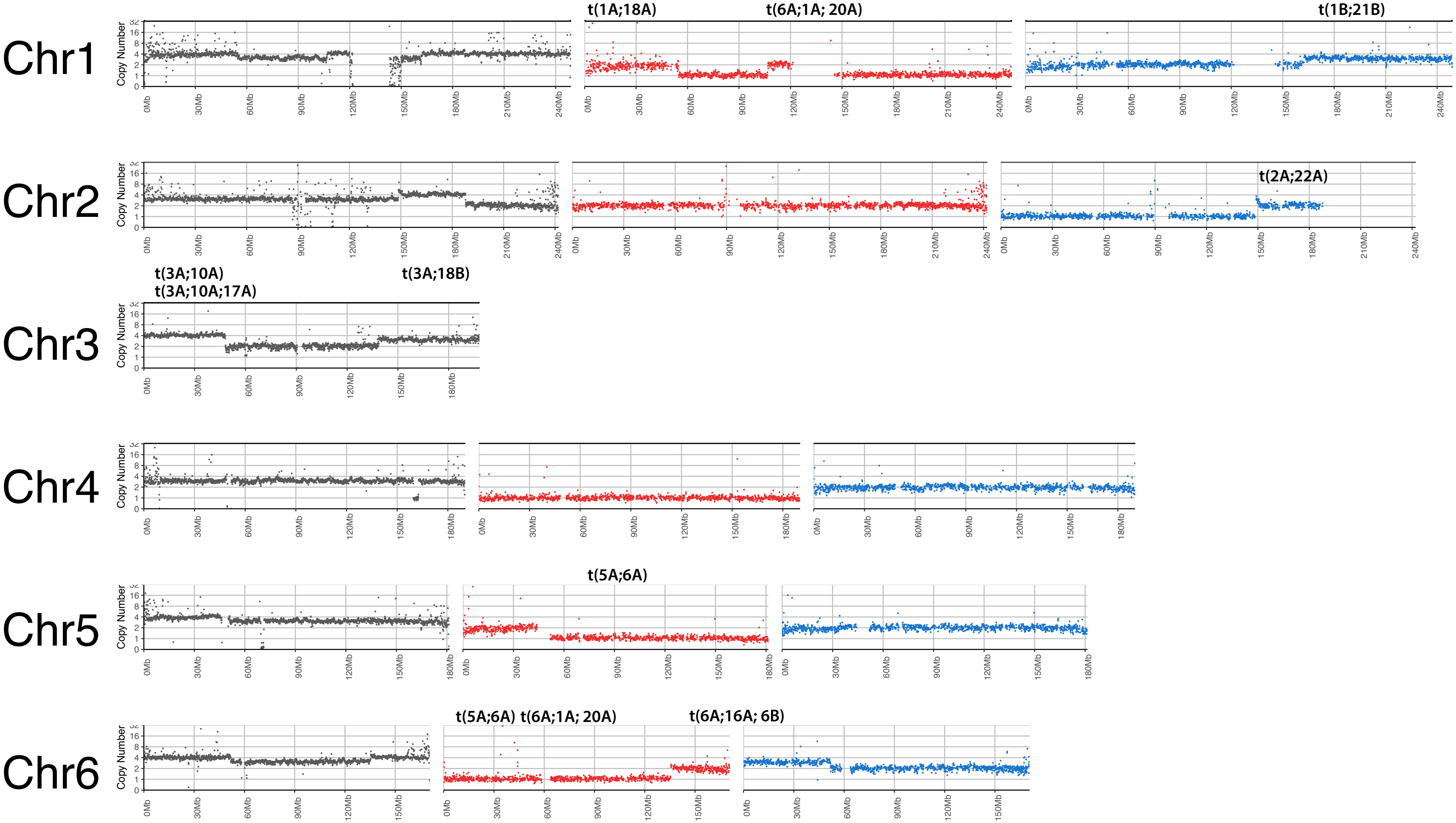

### Chromosomal copy number

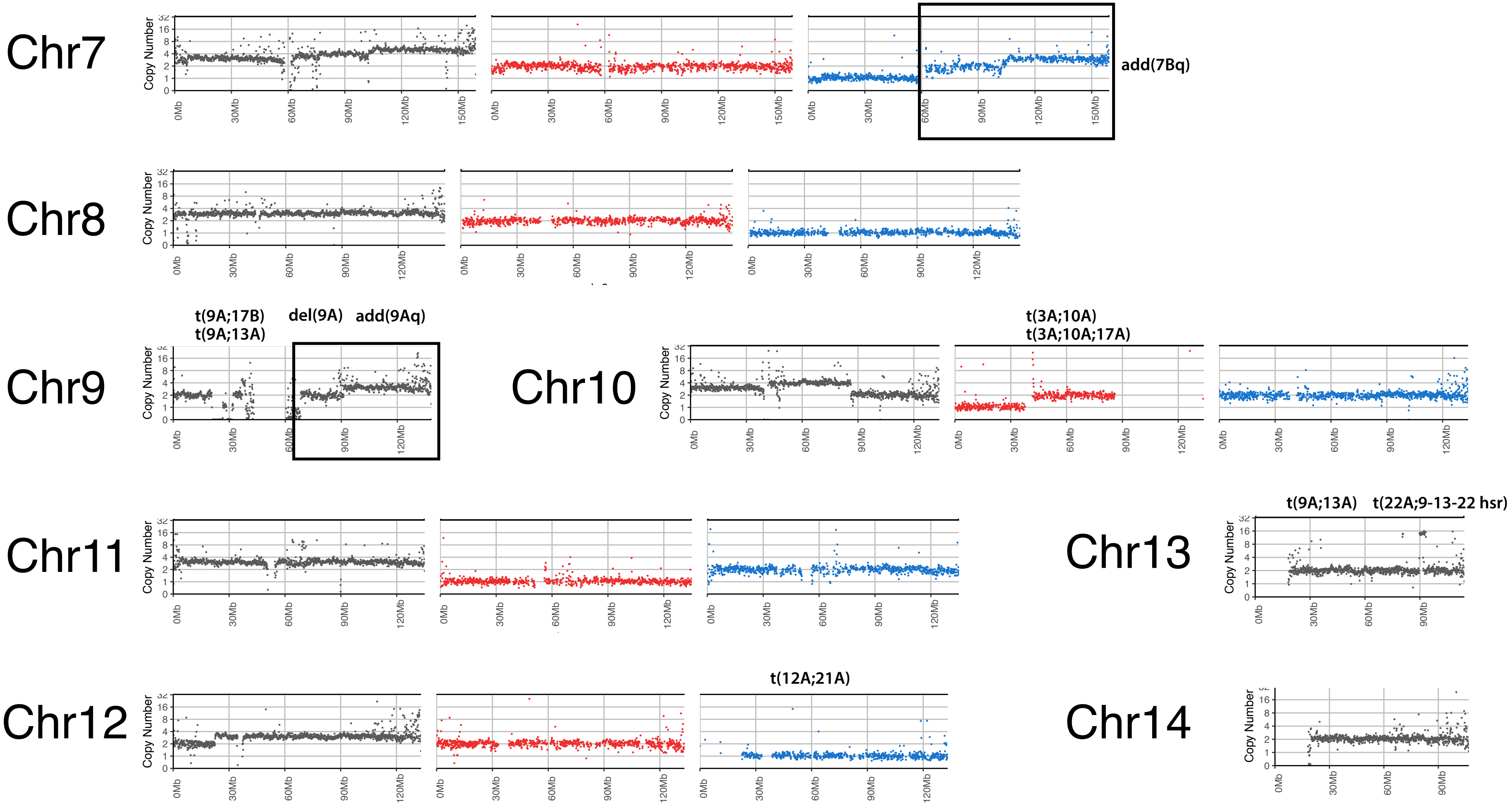

### Chromosomal copy number

Chr15

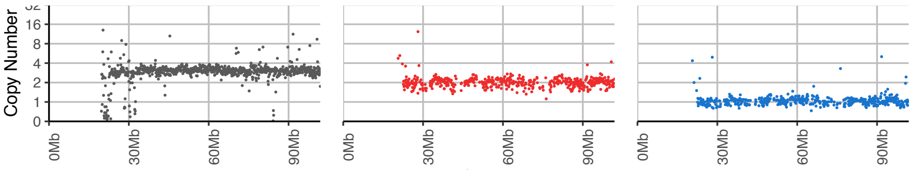


Chr19

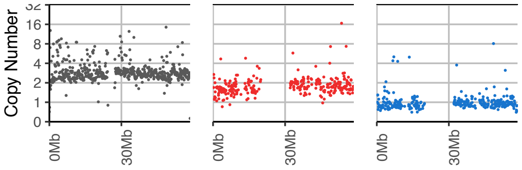


Chr16

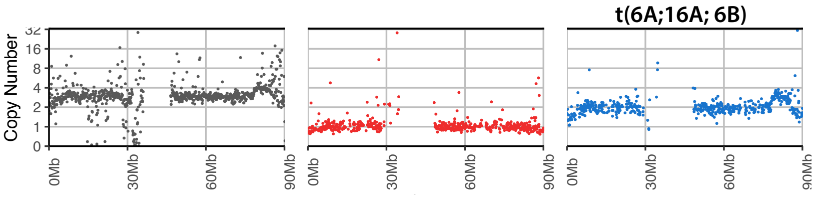


Chr20

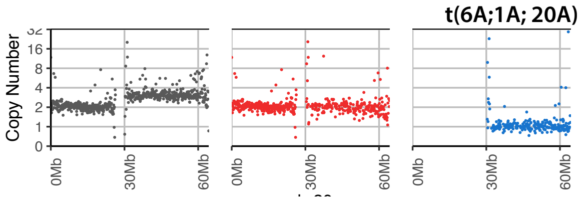


Chr17

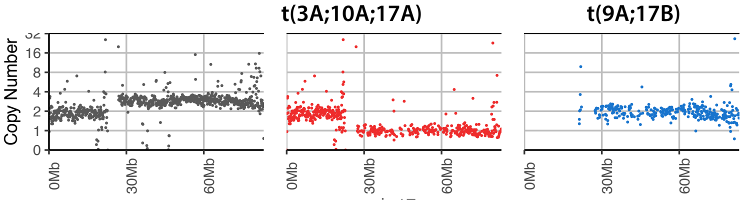


Chr21

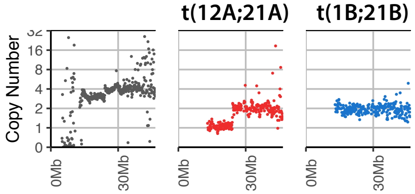


Chr18

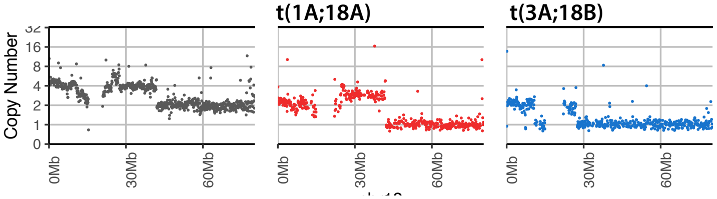


Chr22

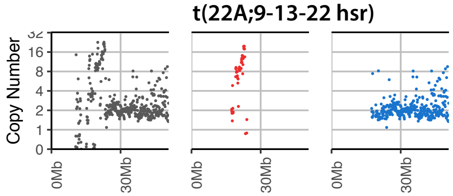


ChrX

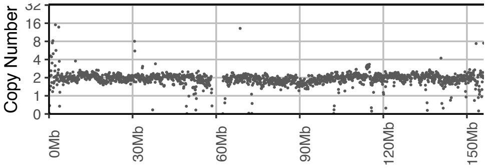

### Digital karyotype

### Structurally normal chromosomes

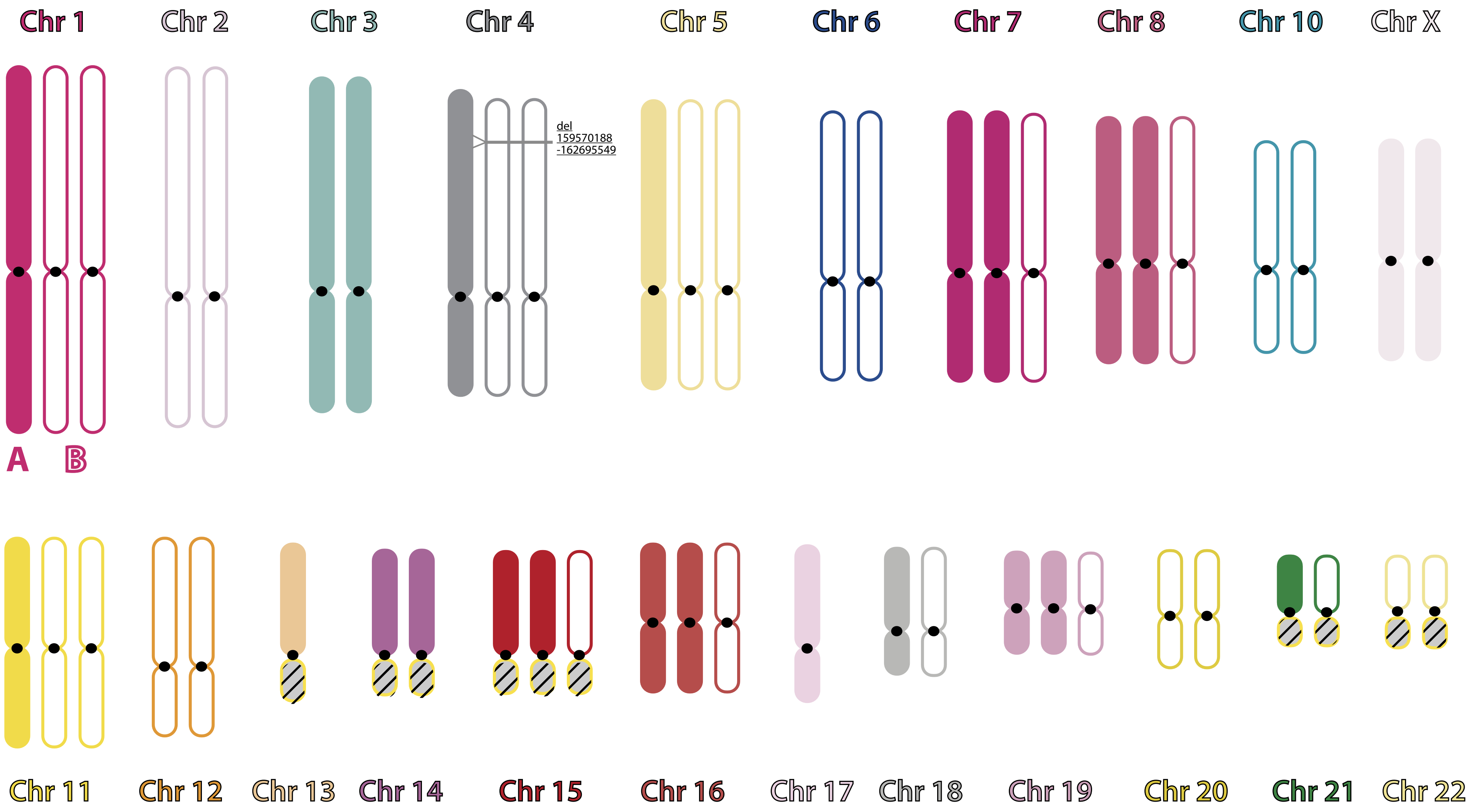

### Marker chromosomes

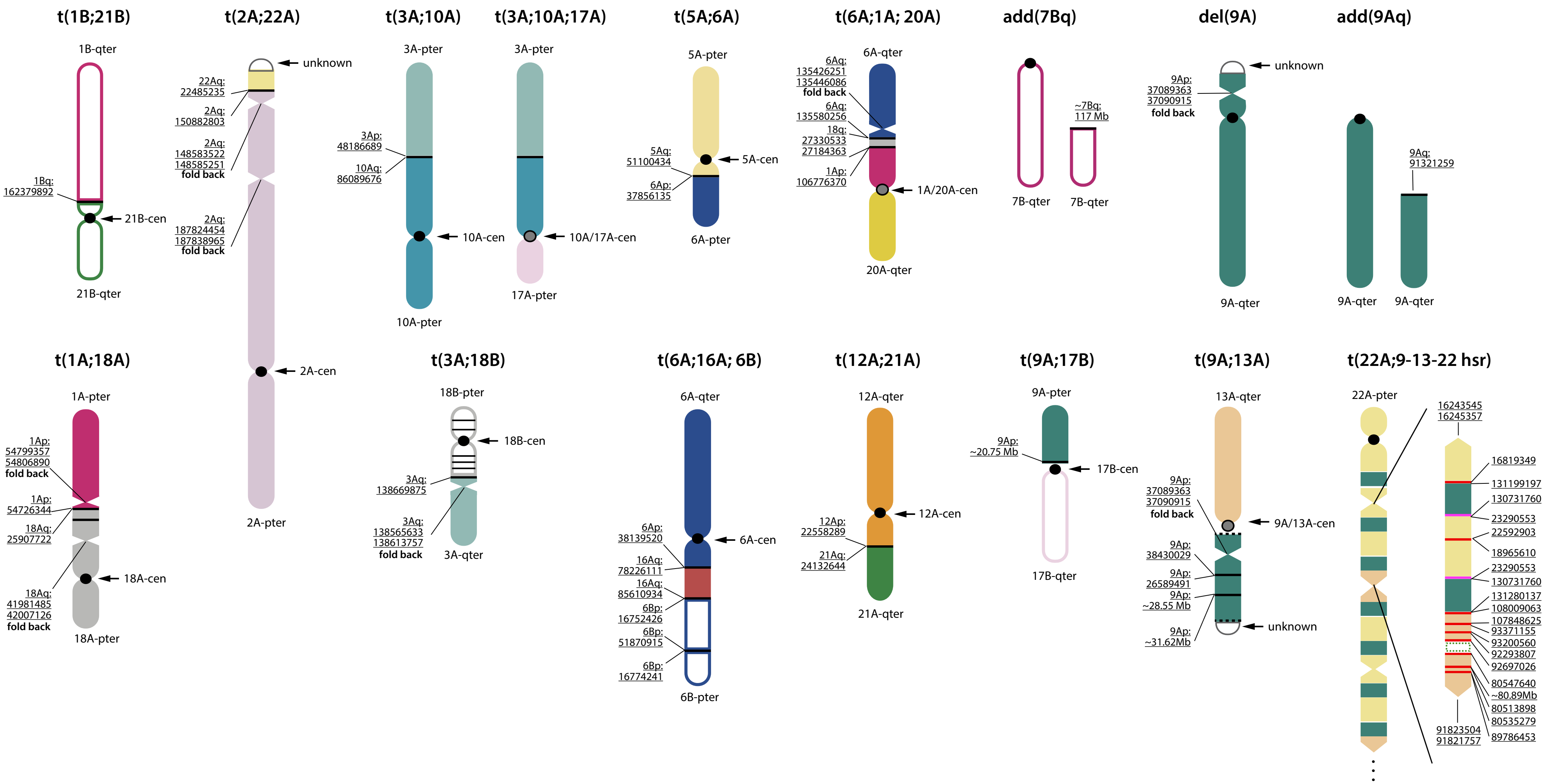

### Marker chromosomes with completely resolved translocations

**t(5A;6A)**

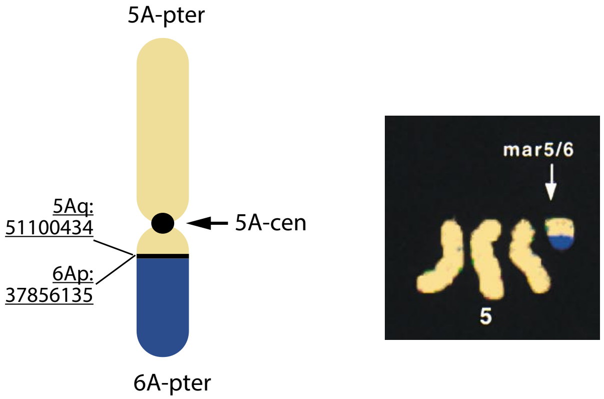


# 10X

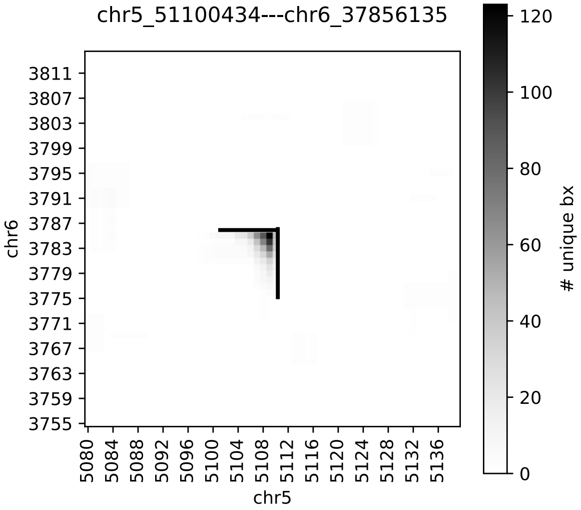


### HiC

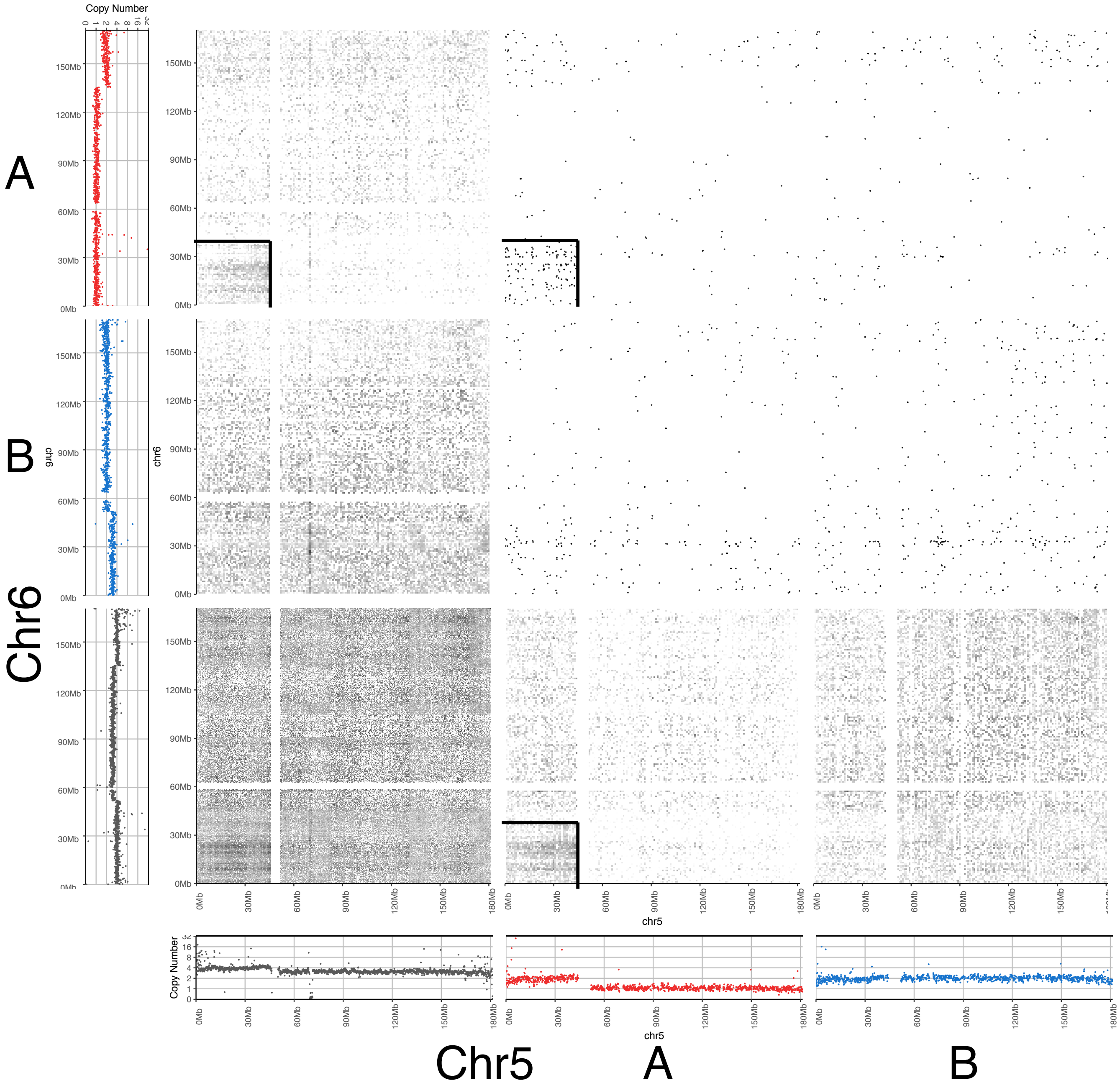

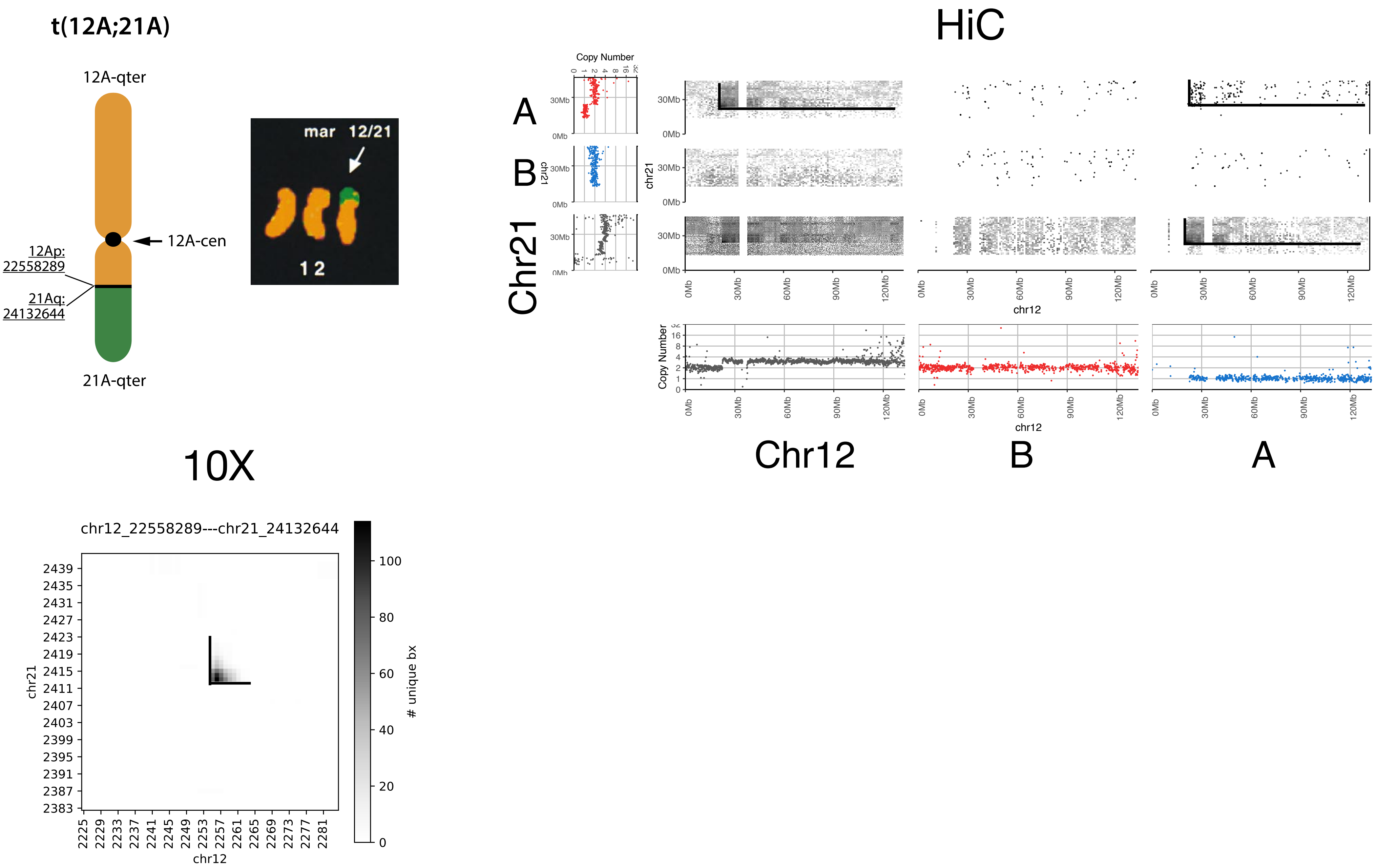
t(12A;21A)
12A-qter
12A-cen
12Ap: 22558289
21Aq: 24132644
21A-qter
mar 12/21
12
HiC
Copy Number
A
B
Chr21
chr21
chr12
Chr12
B
A
10X
chr12_22558289---chr21_24132644
chr21
chr12
### unique bx

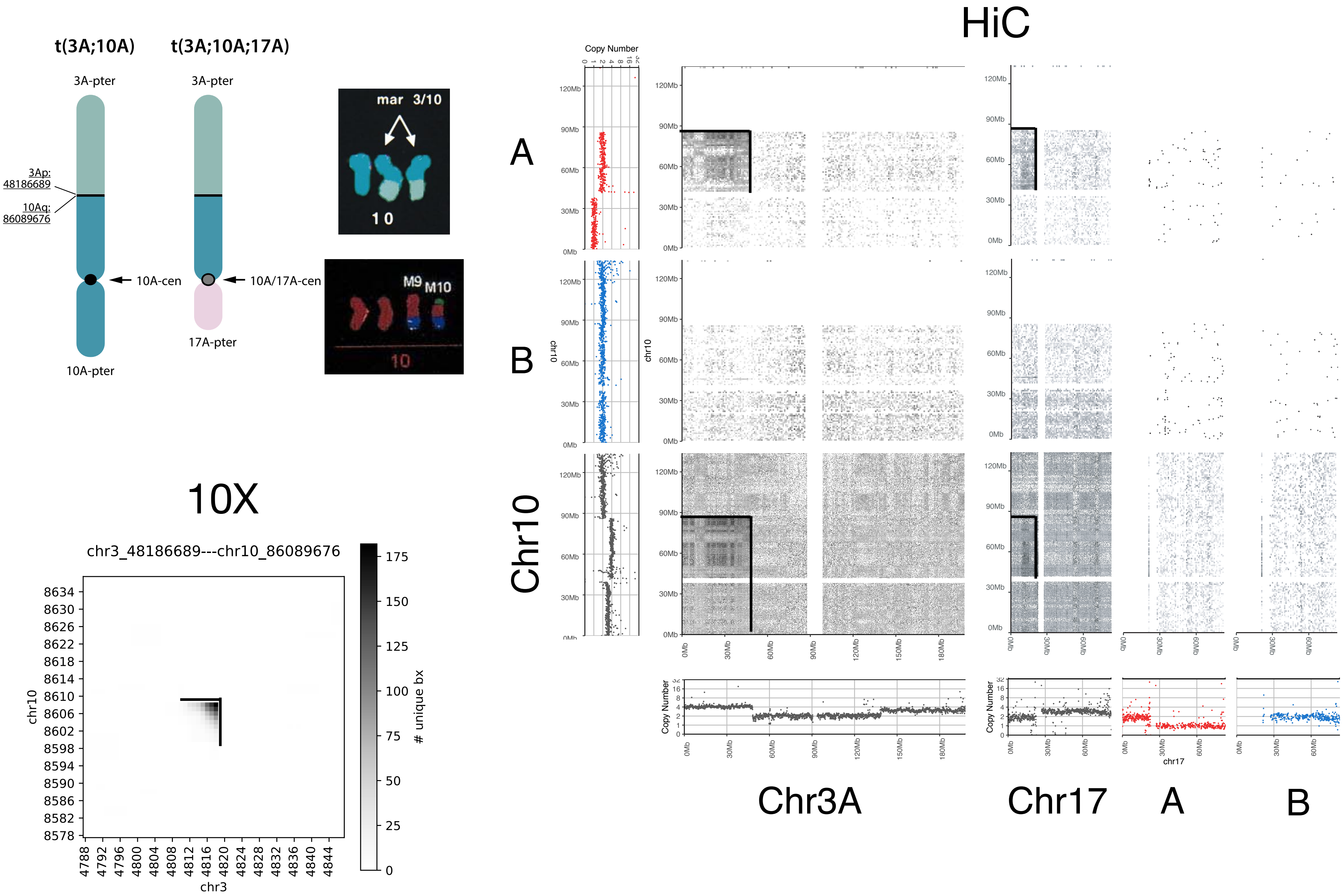

t(3A;10A)
t(3A;10A;17A)
3A-pter
3Ap: 48186689
10Aq: 86089676
10A-cen
10A/17A-cen
17A-pter
10A-pter
mar 3/10
10
M9 M10
10X
chr3_48186689---chr10_86089676
chr10
chr3
### unique bx
HiC
Copy Number
A
B
Chr10
Chr3A
Chr17
chr17

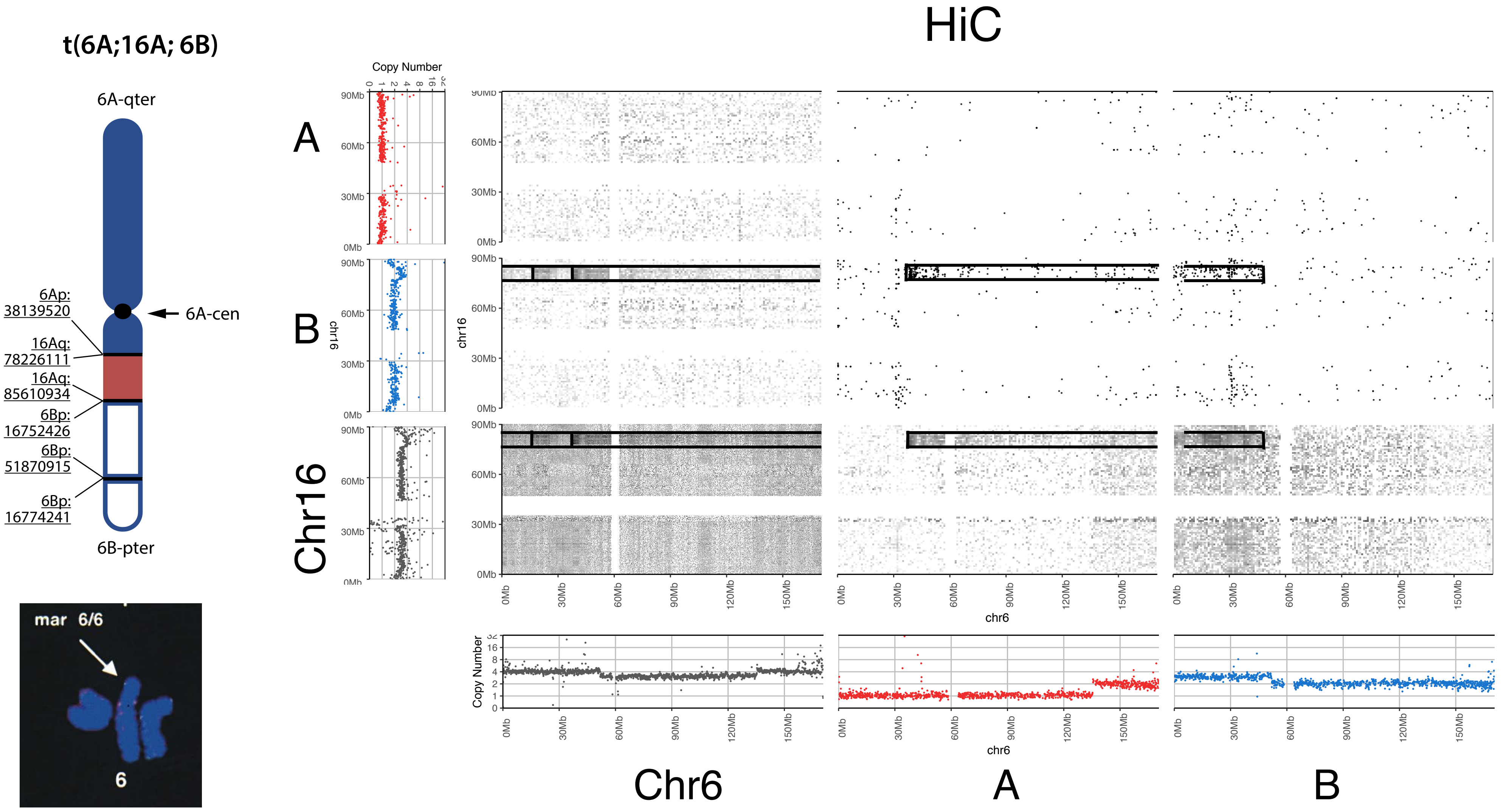
t(6A;16A; 6B)
6A-qter
6Ap:
38139520
16Aq:
78226111
16Aq:
85610934
6Bp:
16752426
6Bp:
51870915
6Bp:
16774241
6A-cen
6B-pter
mar 6/6
6
HiC
Copy Number
A
B
Chr16
chr16
chr6
Chr6
A
B

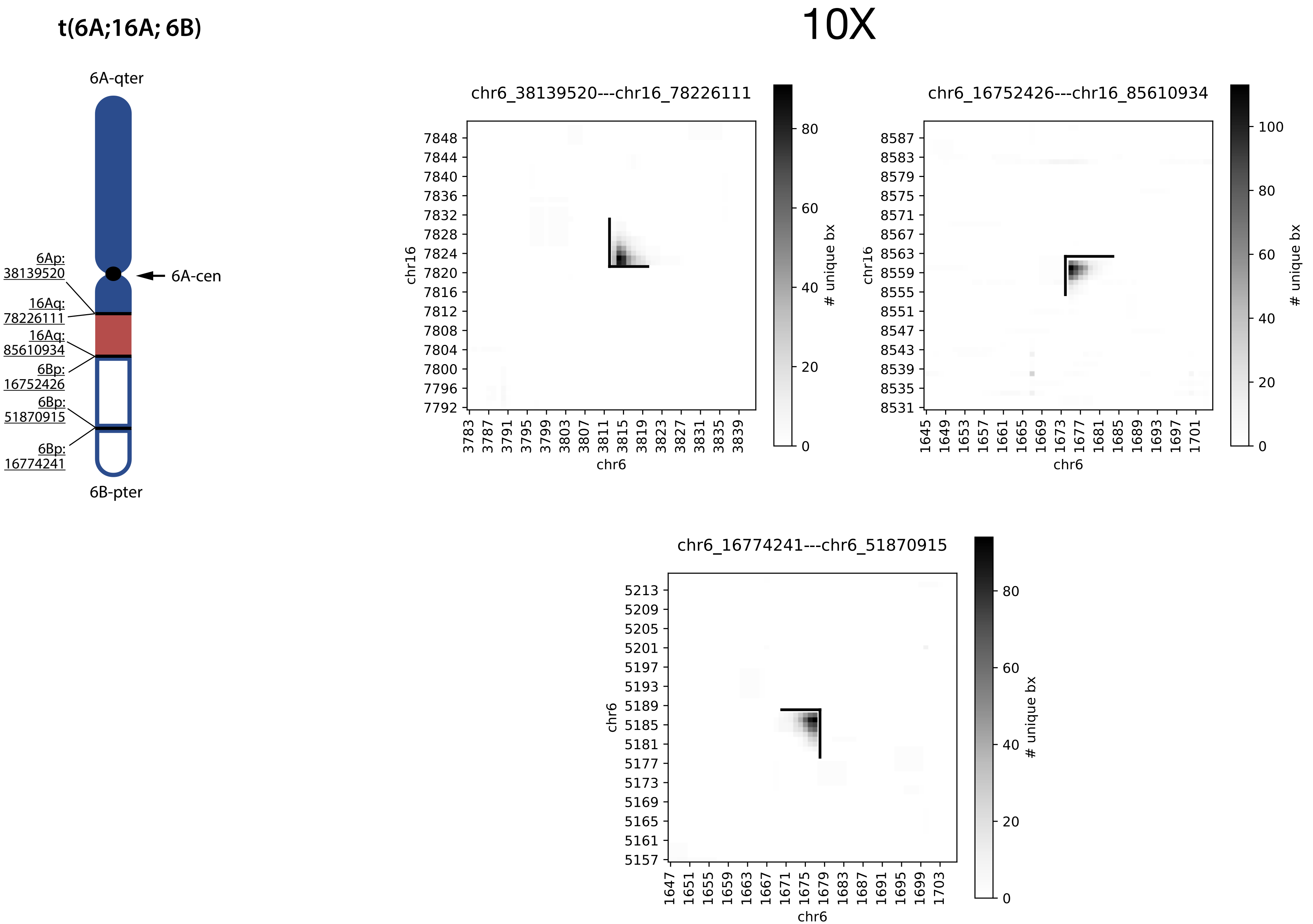
t(6A;16A; 6B)
10X
6A-qter
6Ap:
38139520
16Aq:
78226111
16Aq:
85610934
6Bp:
16752426
6Bp:
51870915
6Bp:
16774241
6A-cen
6B-pter
chr6_38139520---chr16_78226111
chr6_16752426---chr16_85610934
chr6_16774241---chr6_51870915
chr16
chr6
### unique bx

### Marker chromosomes with incompletely resolved translocations

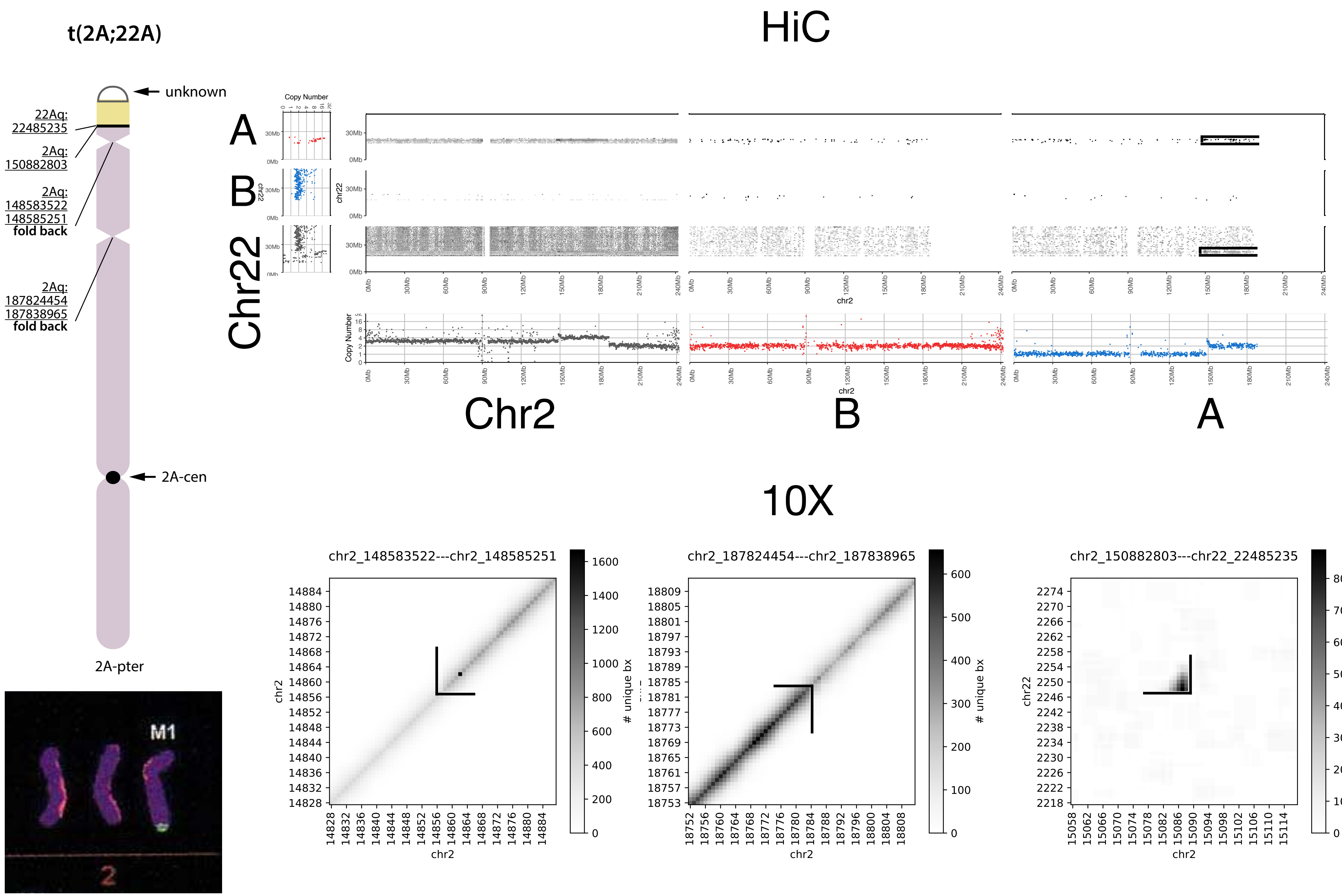
t(2A;22A)
unknown
22Aq:
22485235
2Aq:
150882803
2Aq:
148583522
148585251
fold back
2Aq:
187824454
187838965
fold back
2A-cen
2A-pter
M1
2
HiC
A
B
Chr22
Copy Number
chr22
chr2
Chr2
B
A
10X
chr2_148583522---chr2_148585251
chr2_187824454---chr2_187838965
chr2_150882803---chr22_22485235
### unique bx
chr2
chr22

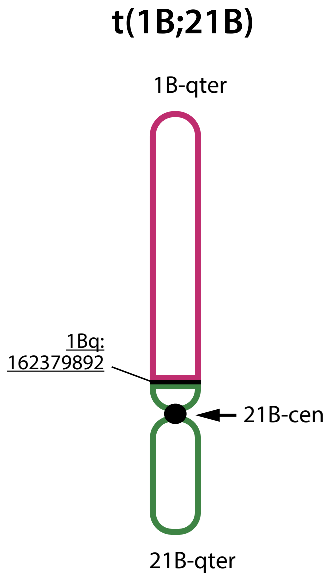
t(1B;21B)
1B-qter
1Bq:
162379892
21B-cen
21B-qter

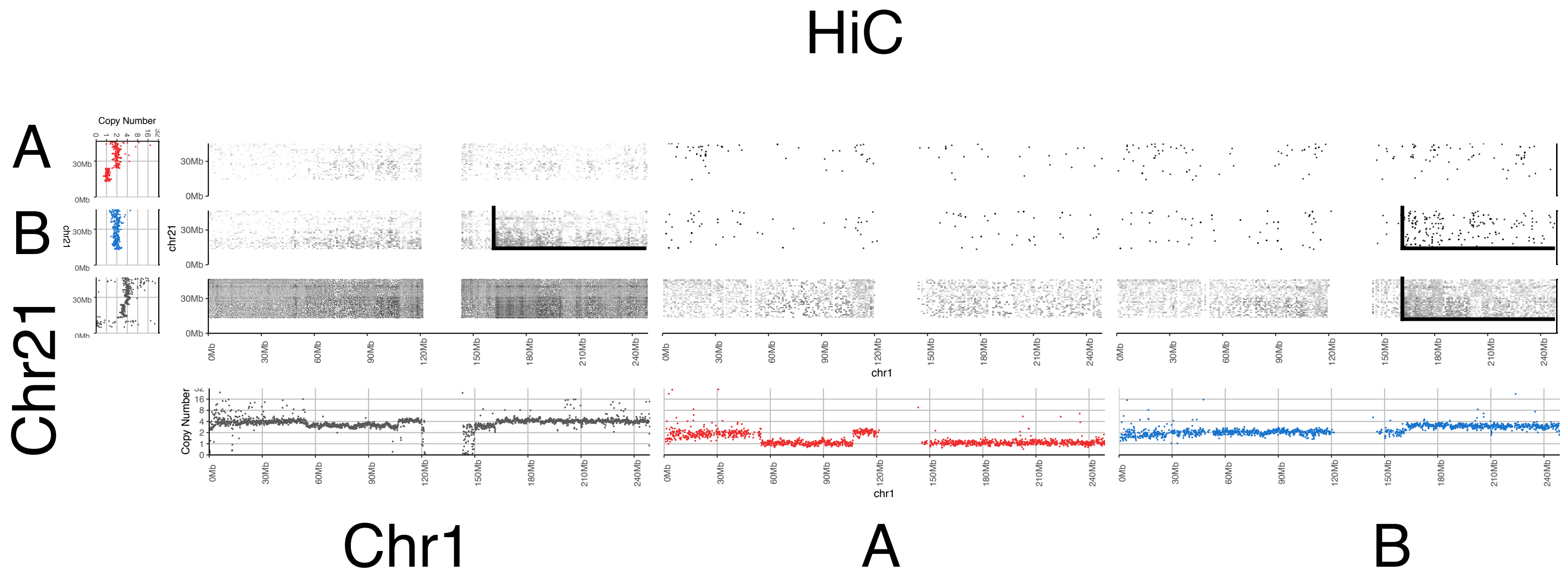
HiC
A
B
Chr21
Copy Number
chr21
chr1
Chr1
A
B

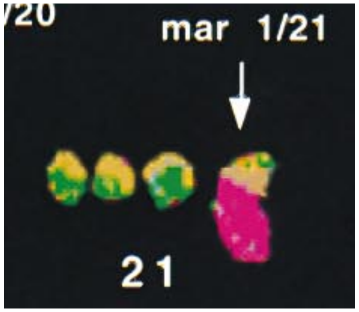
mar 1/21
21

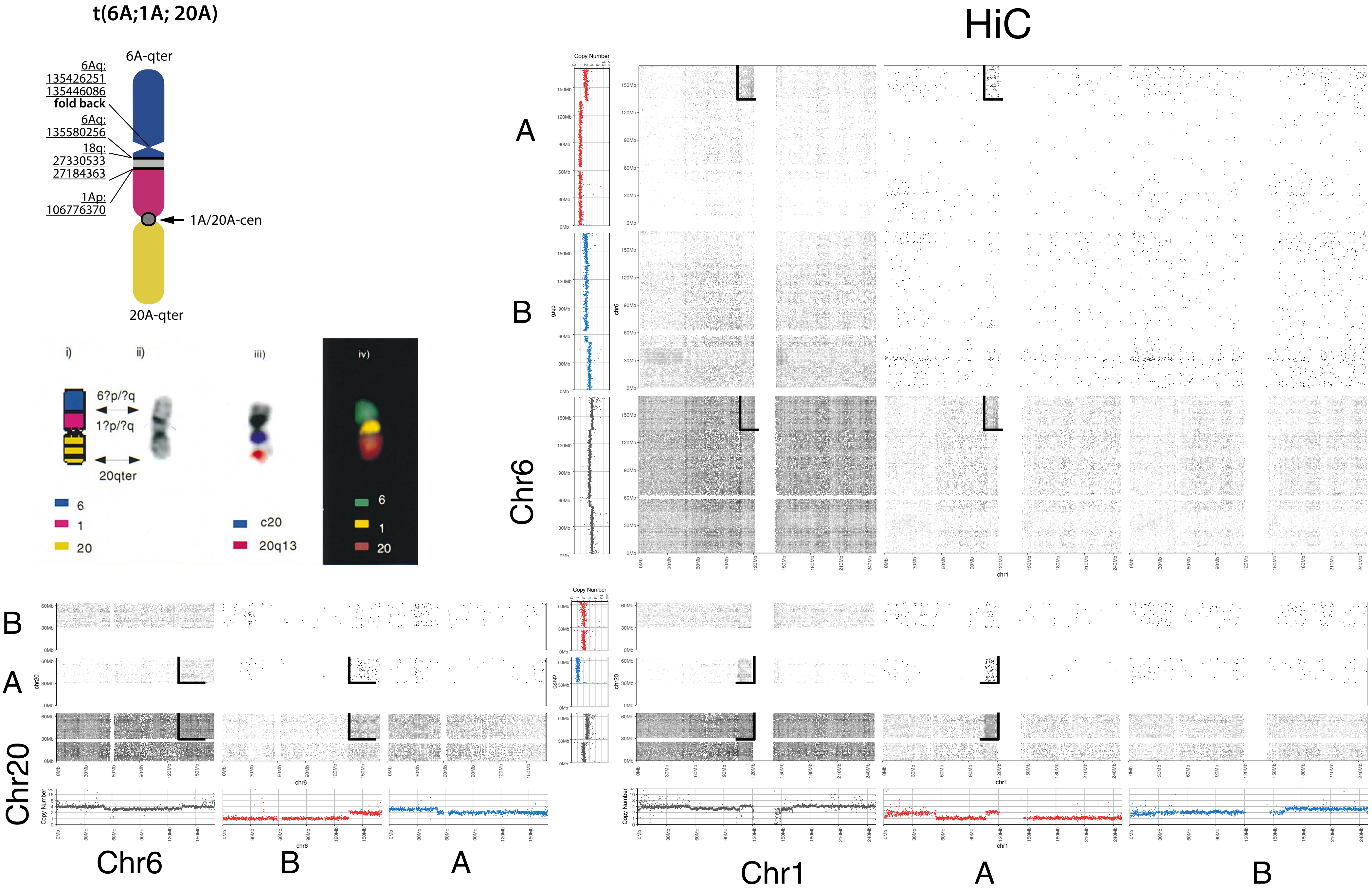

t(6A;1A; 20A)
6A-qter
6Aq:
135426251
135446086
fold back
6Aq:
135580256
18q:
27330533
27184363
1Ap:
106776370
1A/20A-cen
20A-qter
i)
ii)
iii)
iv)
6?p/?q
1?p/?q
20qter
6
1
20
c20
20q13
6
1
20
HiC
A
B
Chr6
Copy Number
B
A
Chr20
Chr6
B
A
Chr1
A
B

**t(6A;1A; 20A)**

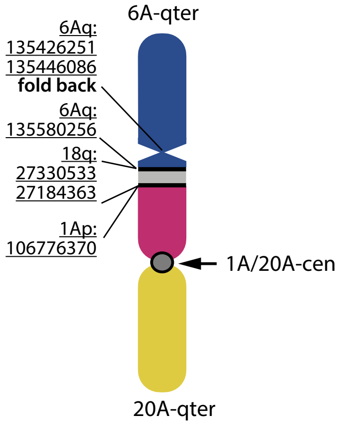


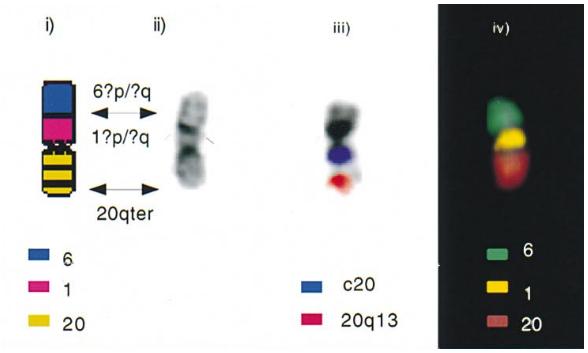


# 10X

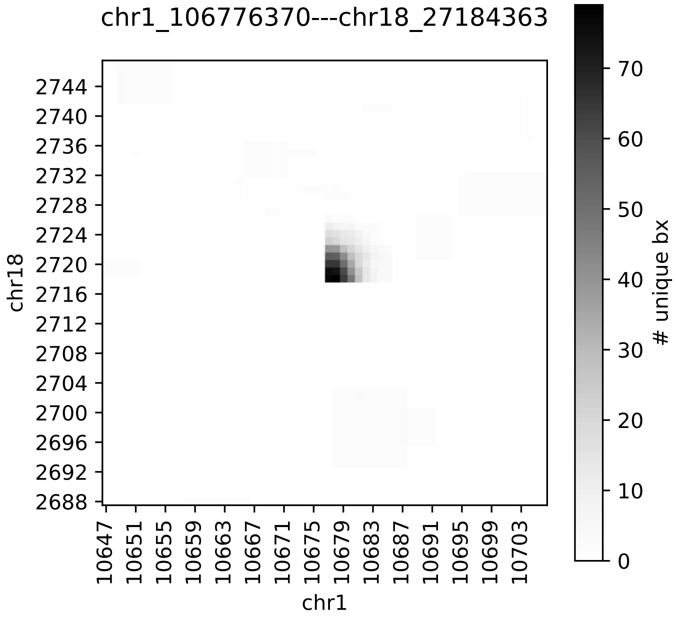


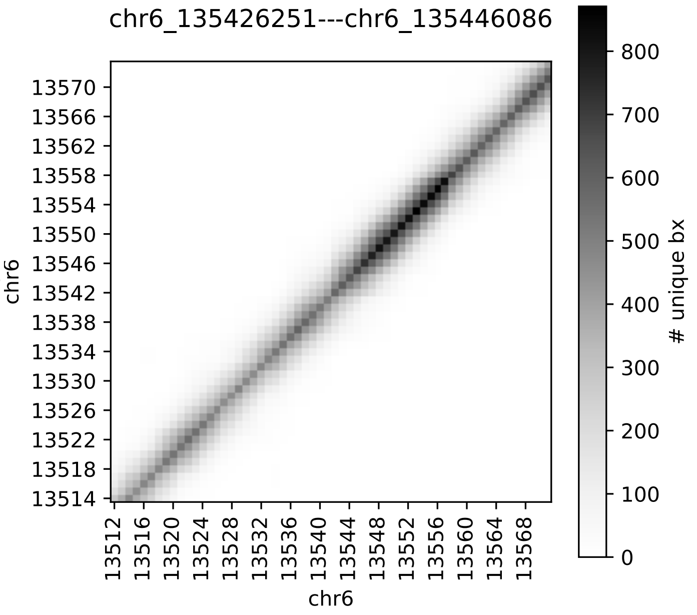

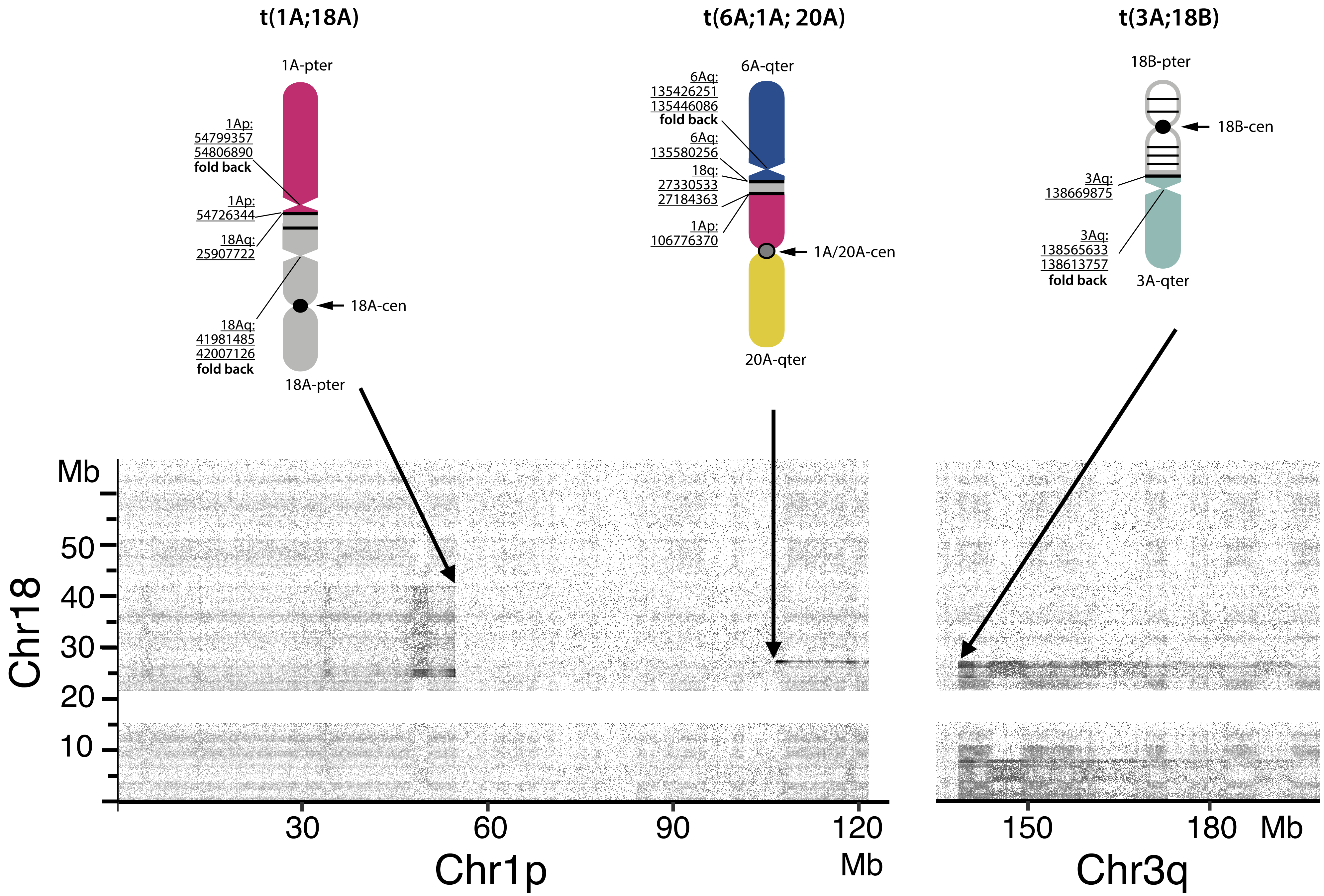
t(1A;18A)
1A-pter
1Ap:
54799357
54806890
fold back
1Ap:
54726344
18Aq:
25907722
18A-cen
18Aq:
41981485
42007126
fold back
18A-pter
t(6A;1A; 20A)
6A-qter
6Aq:
135426251
135446086
fold back
6Aq:
135580256
18q:
27330533
27184363
1Ap:
106776370
1A/20A-cen
20A-qter
t(3A;18B)
18B-pter
18B-cen
3Aq:
138669875
3Aq:
138565633
138613757
fold back
3A-qter
Mb
50
40
30
20
10
Chr18
30
60
90
120
Mb
Chr1p
150
180
Mb
Chr3q

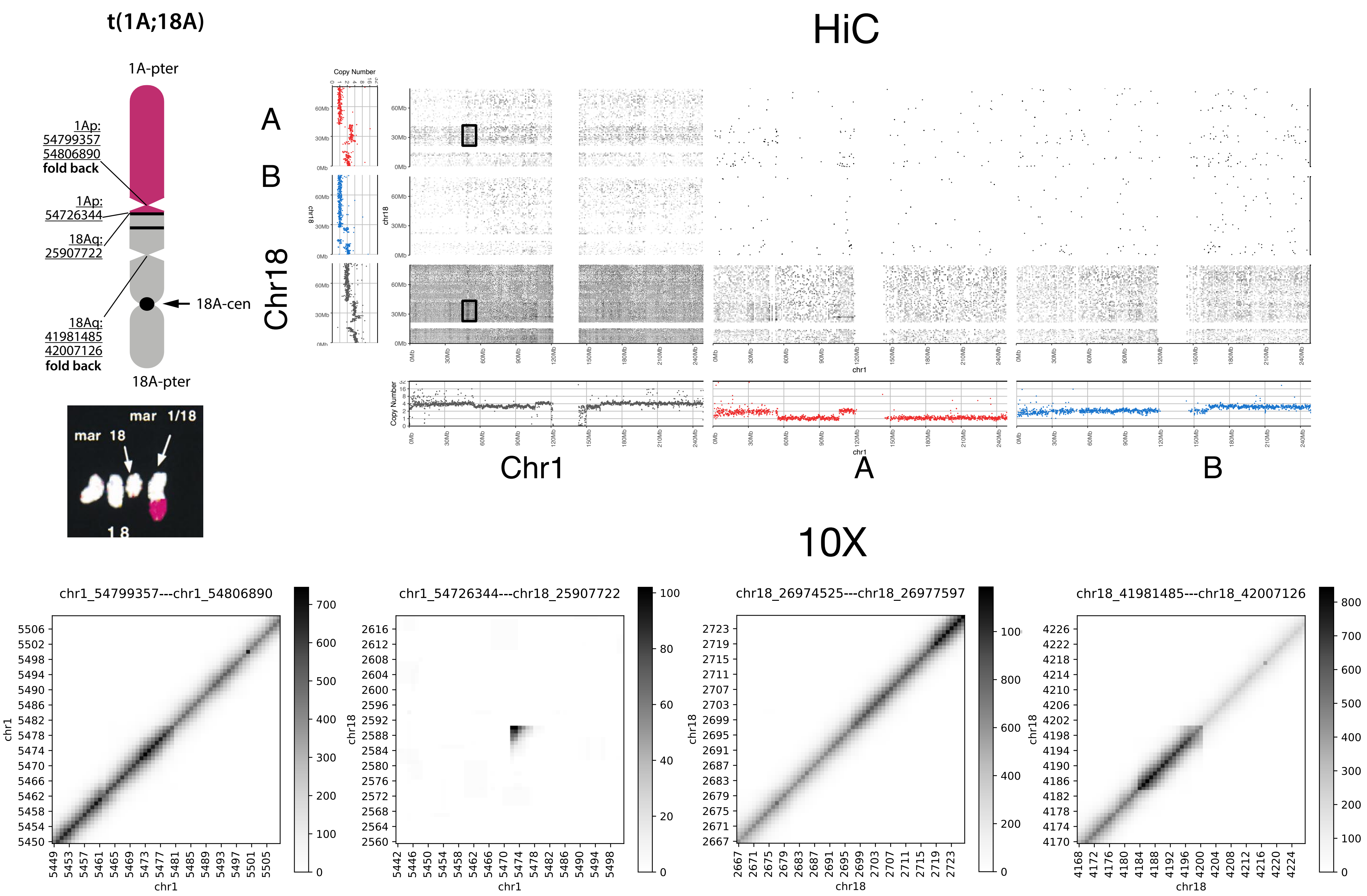
t(1A;18A)
1A-pter
1Ap:
54799357
54806890
fold back
1Ap:
54726344
18Aq:
25907722
18A-cen
18Aq:
41981485
42007126
fold back
18A-pter
mar 1/18
mar 18
18
HiC
A
B
Chr18
Copy Number
Chr1
A
B
10X
chr1_54799357---chr1_54806890
chr1_54726344---chr18_25907722
chr18_26974525---chr18_26977597
chr18_41981485---chr18_42007126
chr1
chr18

t(3A;18B)

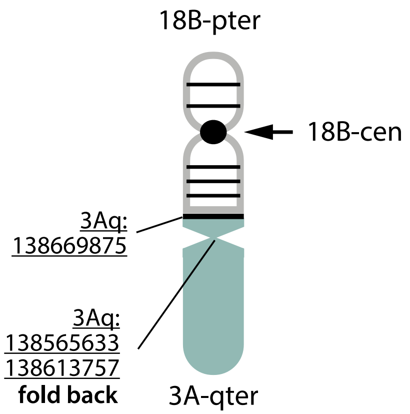


10X

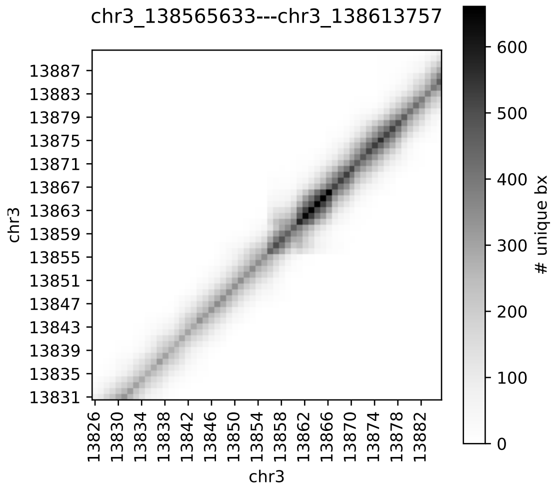


HiC

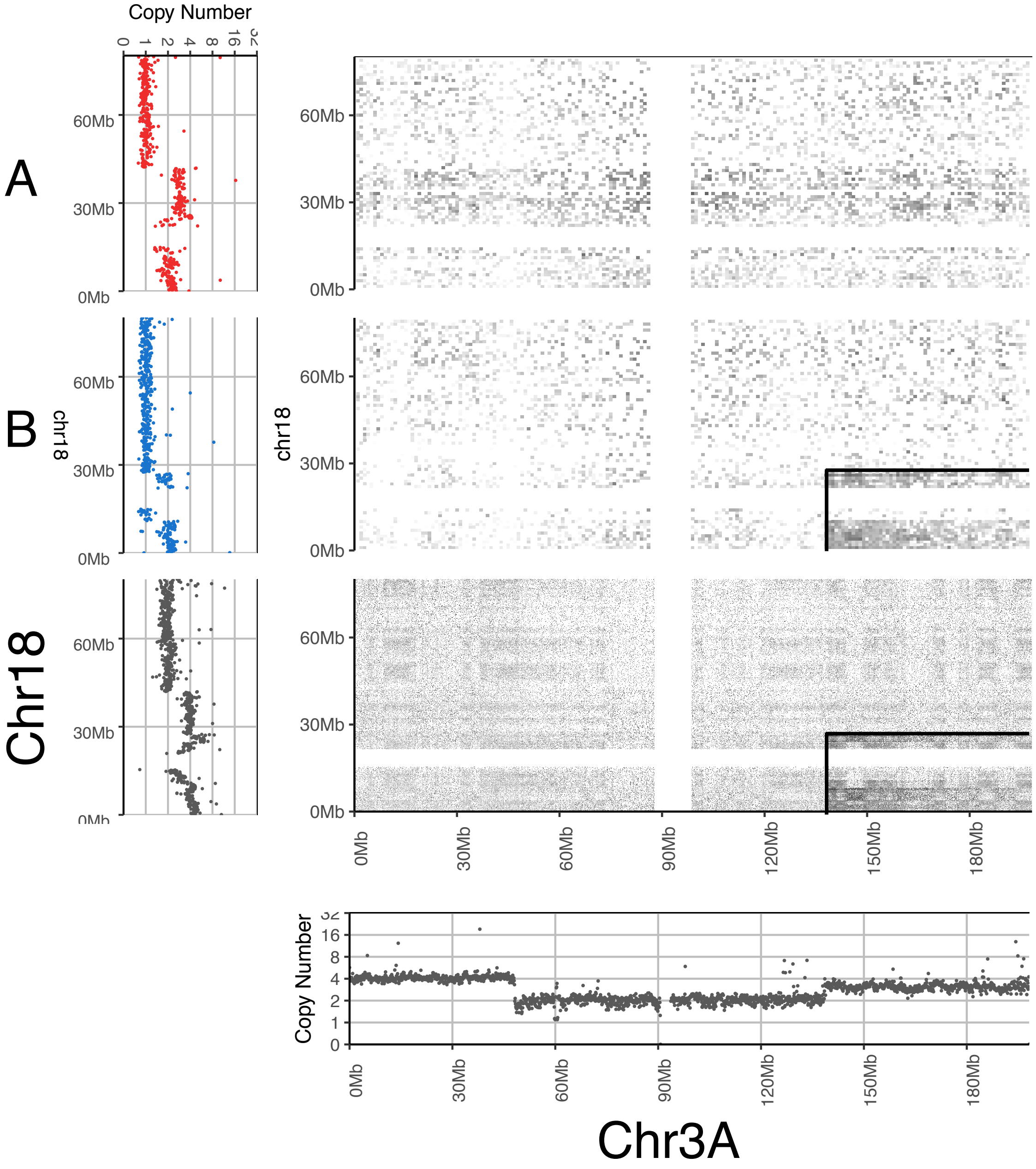

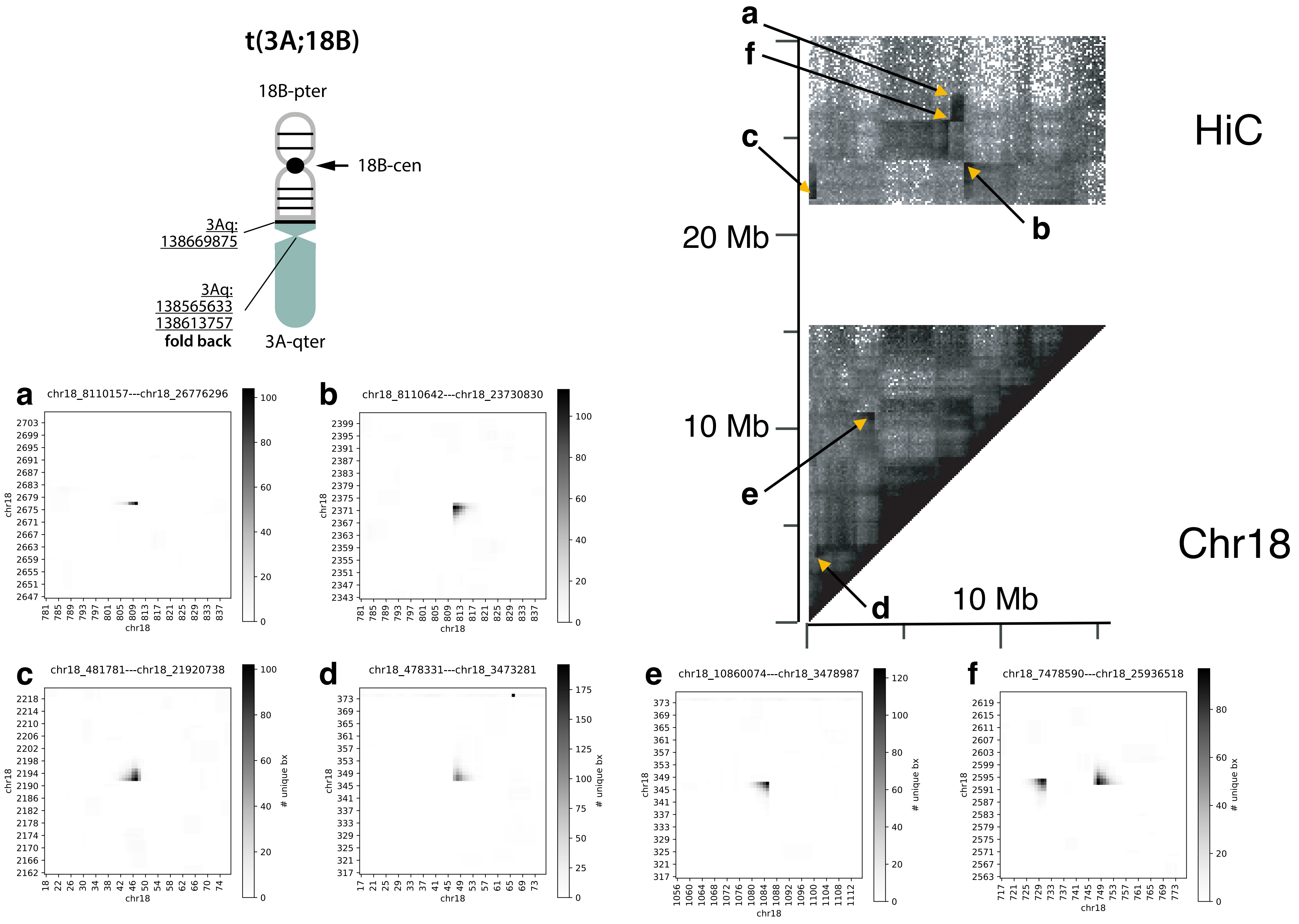
t(3A;18B)
18B-pter
18B-cen
3Aq: 138669875
3Aq: 138565633 138613757
fold back
3A-qter
a
f
c
b
e
d
20 Mb
10 Mb
10 Mb
HiC
Chr18
a chr18_8110157---chr18_26776296
b chr18_8110642---chr18_23730830
c chr18_481781---chr18_21920738
d chr18_478331---chr18_3473281
e chr18_10860074---chr18_3478987
f chr18_7478590---chr18_25936518
chr18
### unique bx

### Chromosomal copy number & rearrangements

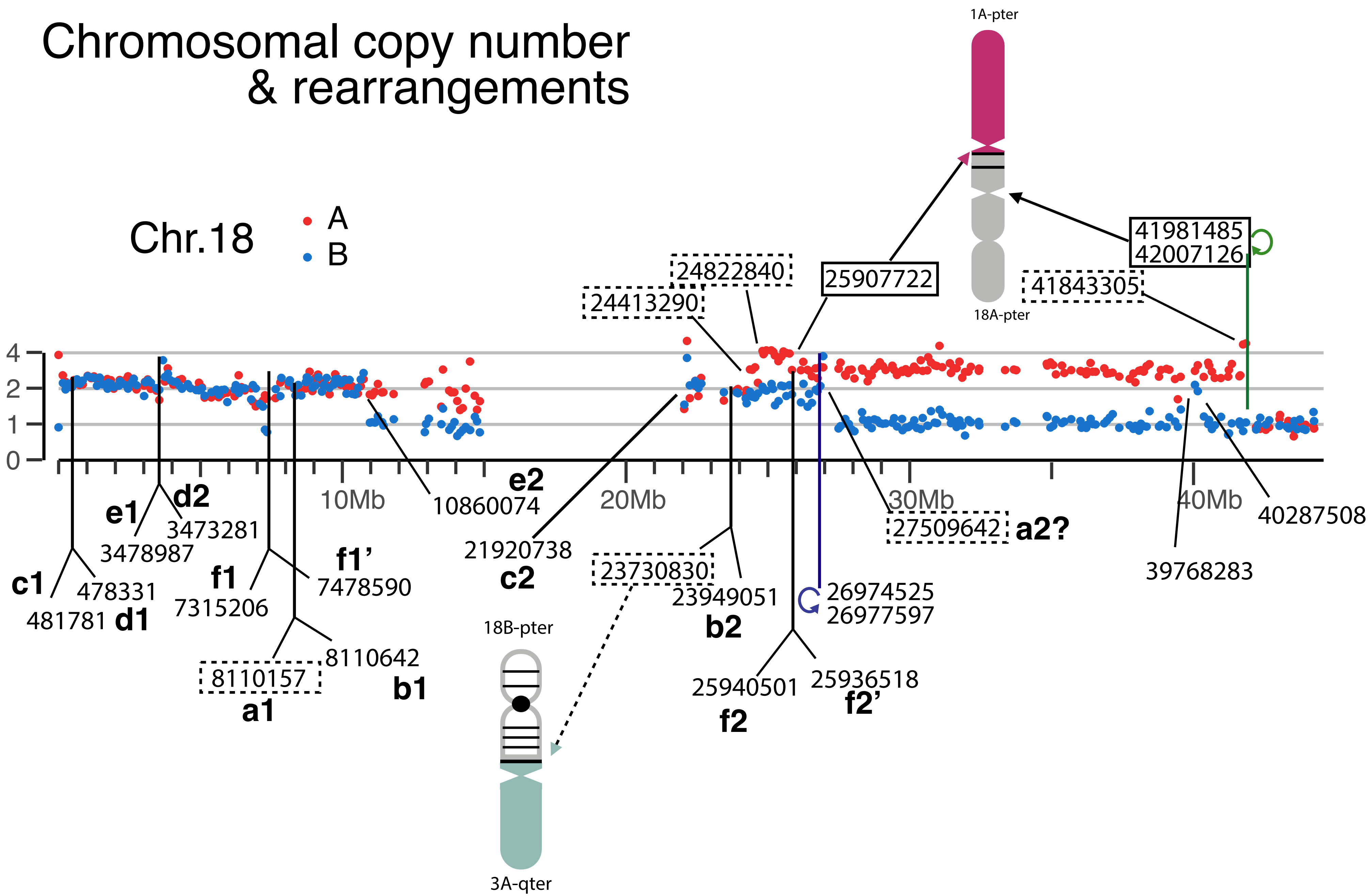

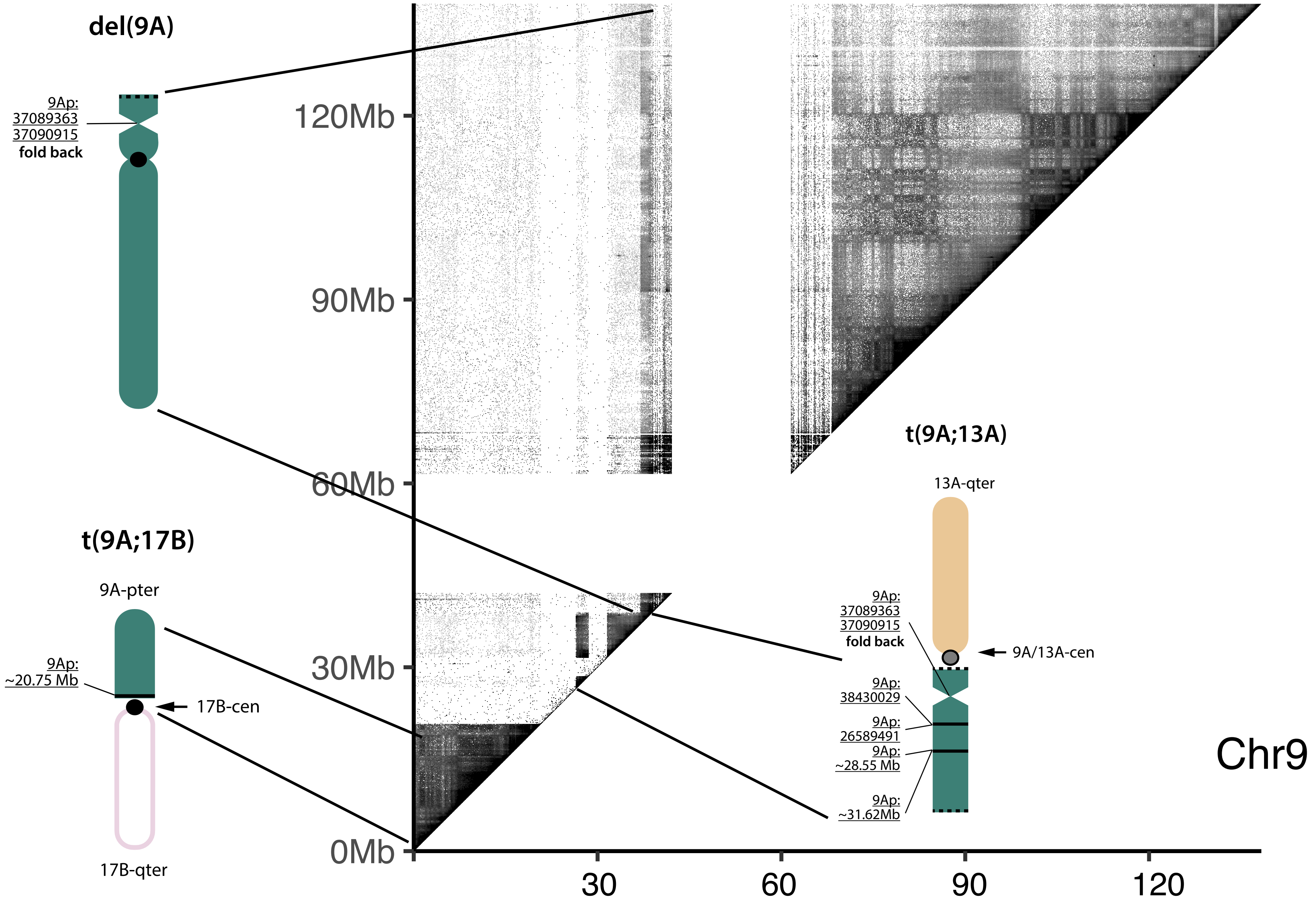

del(9A)
9Ap:
37089363
37090915
fold back
120Mb
90Mb
60Mb
30Mb
0Mb
t(9A;17B)
9A-pter
9Ap:
~20.75 Mb
17B-cen
17B-qter
30
60
90
120
t(9A;13A)
13A-qter
9Ap:
37089363
37090915
fold back
9A/13A-cen
9Ap:
38430029
9Ap:
26589491
9Ap:
~28.55 Mb
9Ap:
~31.62Mb
Chr9

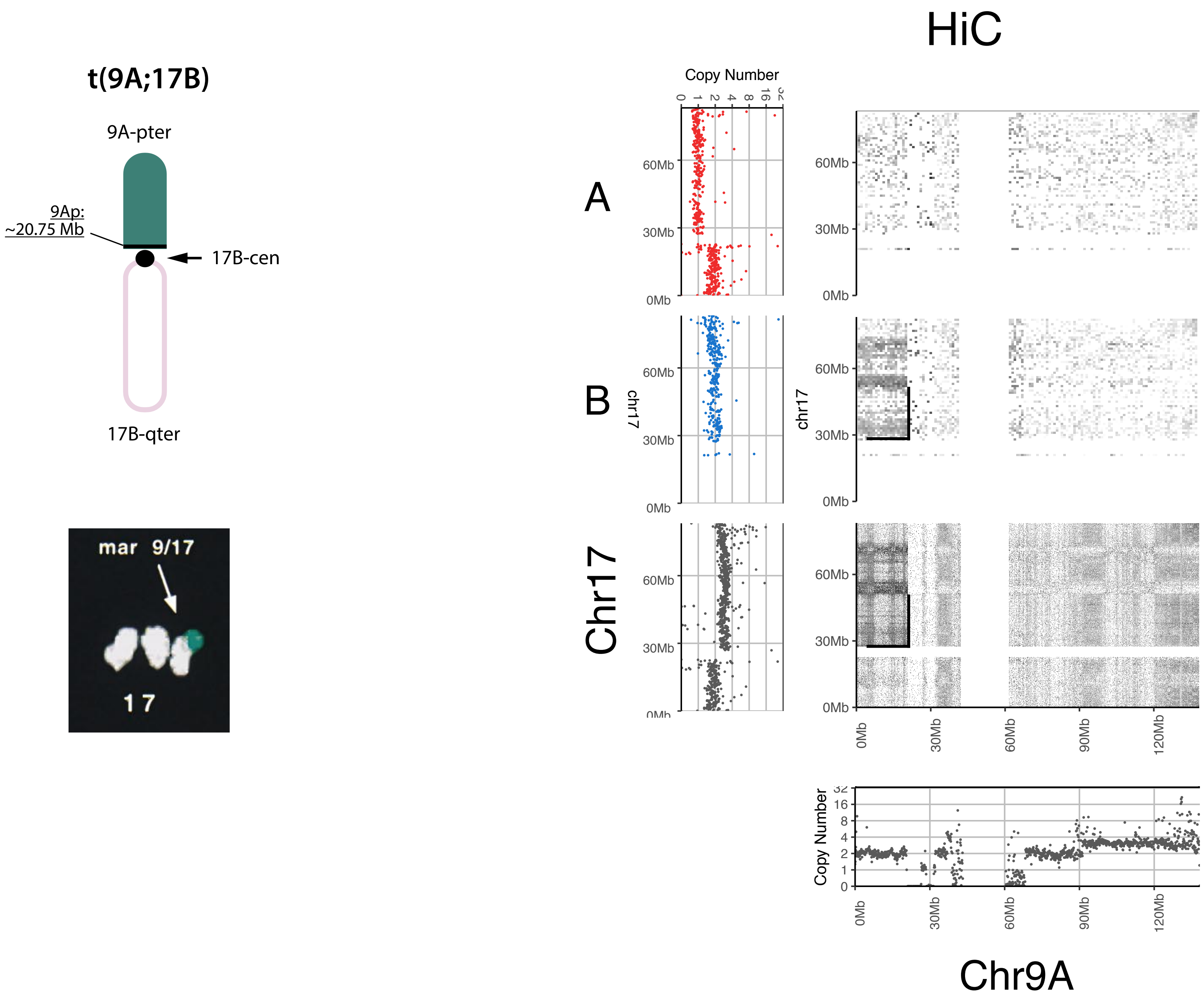
t(9A;17B)
9A-pter
9Ap:
~20.75 Mb
17B-cen
17B-qter
mar 9/17
17
HiC
Copy Number
0
1
2
4
8
16
32
A
B
Chr17
chr17
60Mb
30Mb
0Mb
90Mb
120Mb
Chr9A

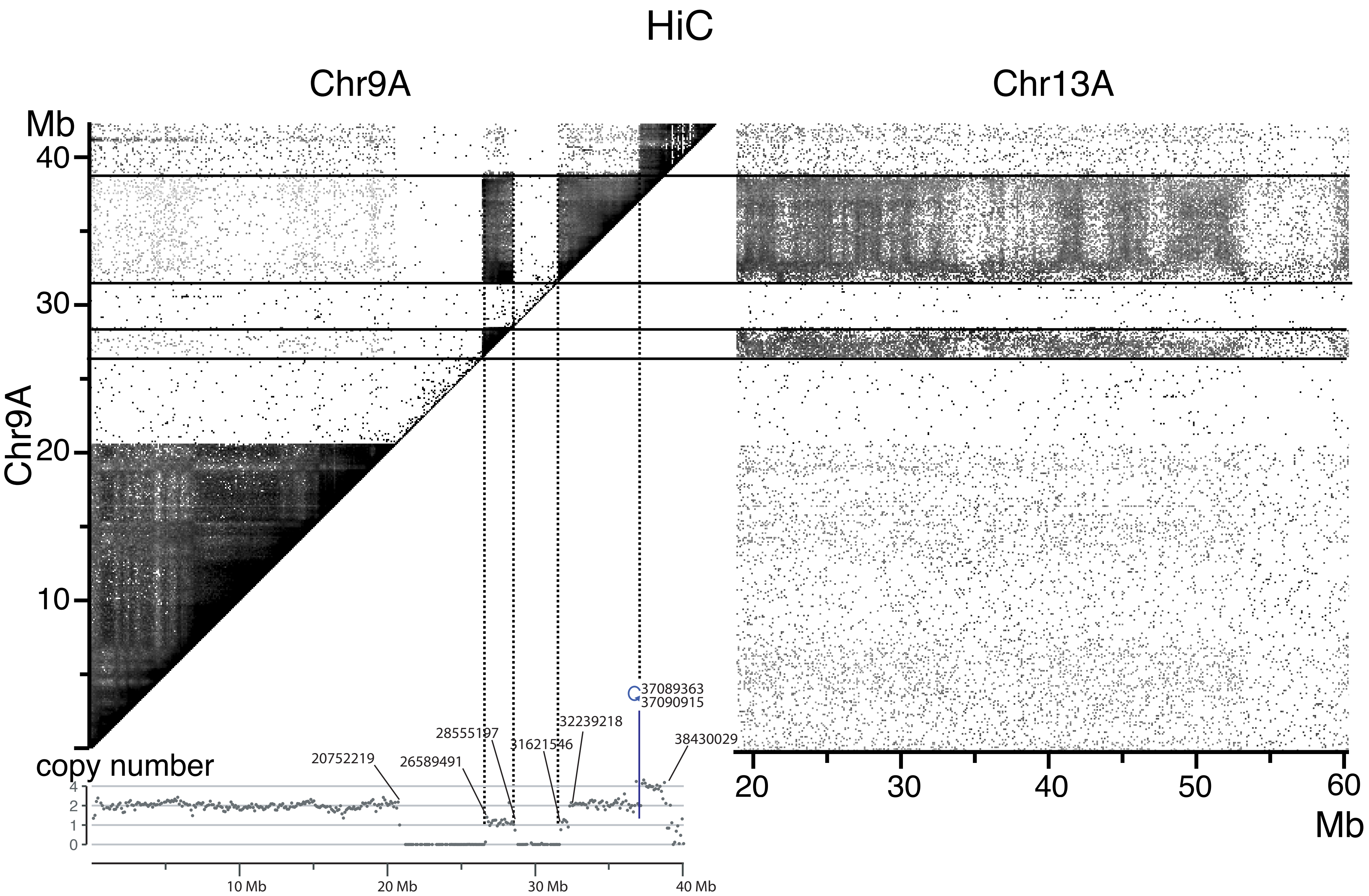
HiC
Chr9A
Chr13A
Mb
40
30
20
10
Chr9A
copy number
4
2
1
0
20752219
26589491
28555197
31621546
32239218
37089363
37090915
38430029
10 Mb
20 Mb
30 Mb
40 Mb
20
30
40
50
60
Mb

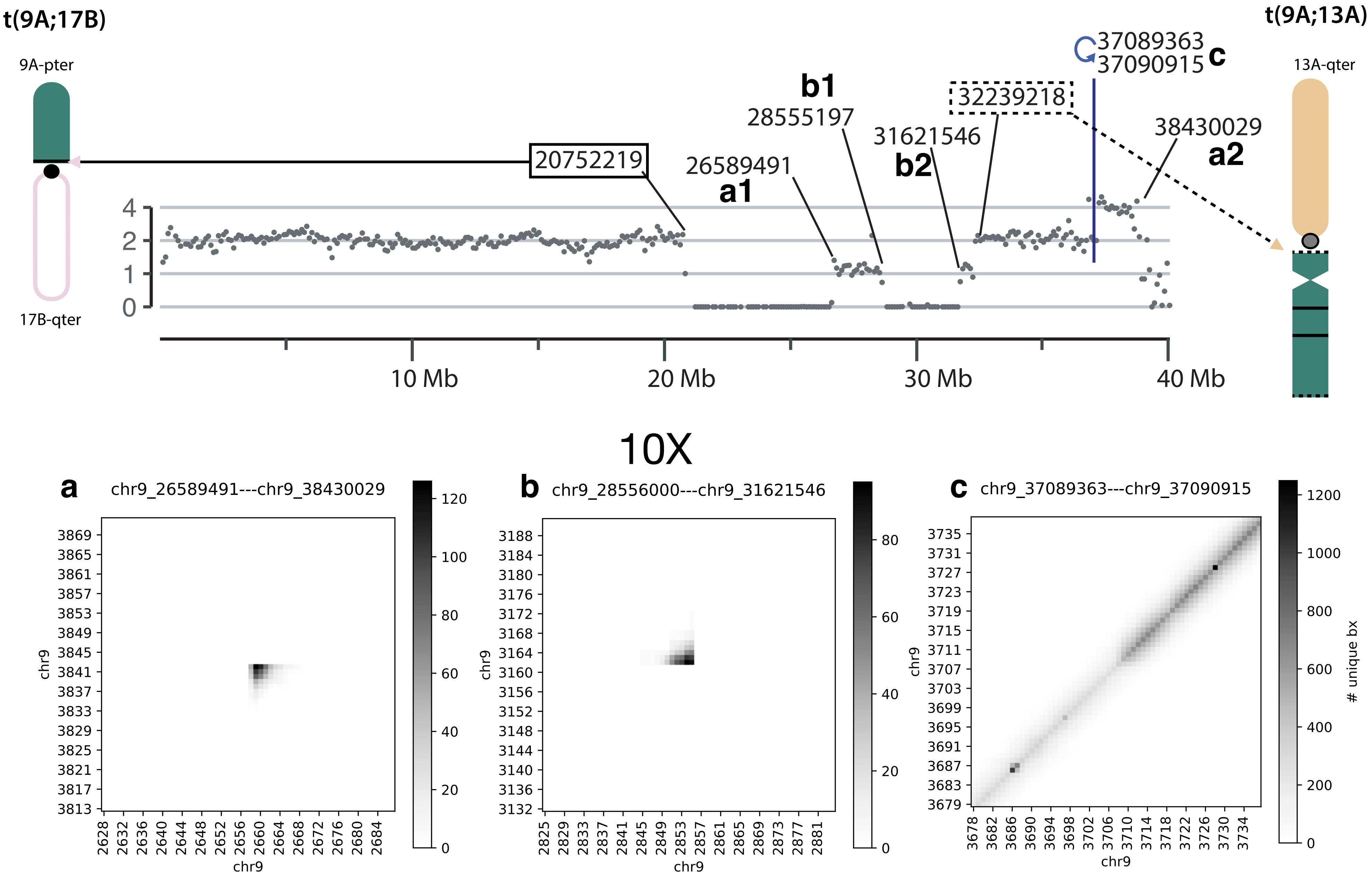

t(9A;17B)
9A-pter
17B-qter
20752219
26589491
a1
b1
28555197
31621546
b2
32239218
37089363
37090915
c
38430029
a2
t(9A;13A)
13A-qter
0
1
2
4
10 Mb
20 Mb
30 Mb
40 Mb
10X
a
chr9_26589491---chr9_38430029
b
chr9_28556000---chr9_31621546
c
chr9_37089363---chr9_37090915
chr9
### unique bx

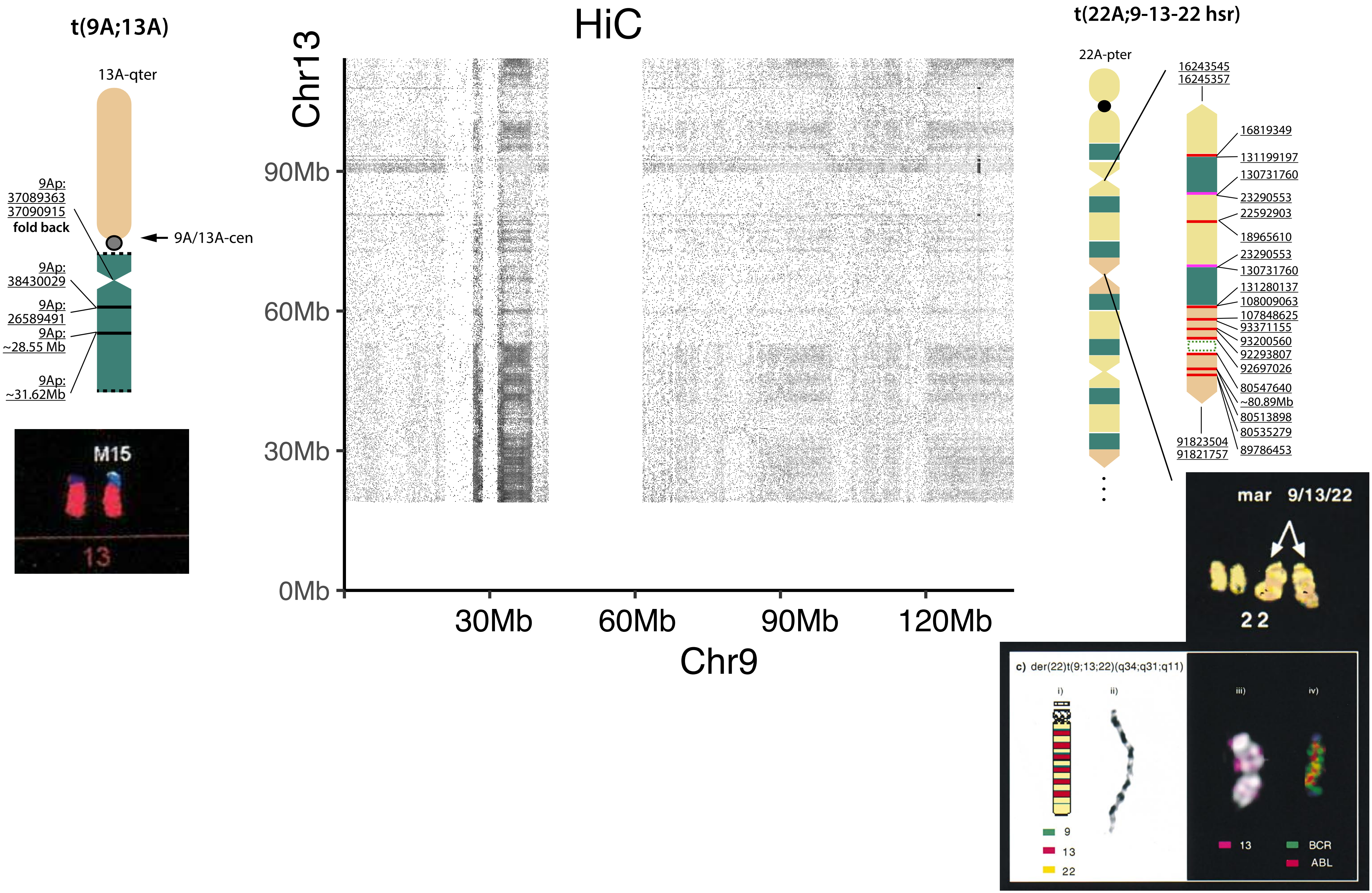
t(9A;13A)
13A-qter
9Ap:
37089363
37090915
fold back
9A/13A-cen
9Ap:
38430029
9Ap:
26589491
9Ap:
~28.55 Mb
9Ap:
~31.62Mb
M15
13
HiC
Chr13
90Mb
60Mb
30Mb
0Mb
30Mb
60Mb
90Mb
120Mb
Chr9
t(22A;9-13-22 hsr)
22A-pter
16243545
16245357
16819349
131199197
130731760
23290553
22592903
18965610
23290553
130731760
131280137
108009063
107848625
93371155
93200560
92293807
92697026
80547640
~80.89Mb
80513898
80535279
89786453
91823504
91821757
mar 9/13/22
22
c) der(22)t(9;13;22)(q34;q31;q11)
i)
ii)
iii)
iv)
9
13
22
13
BCR
ABL

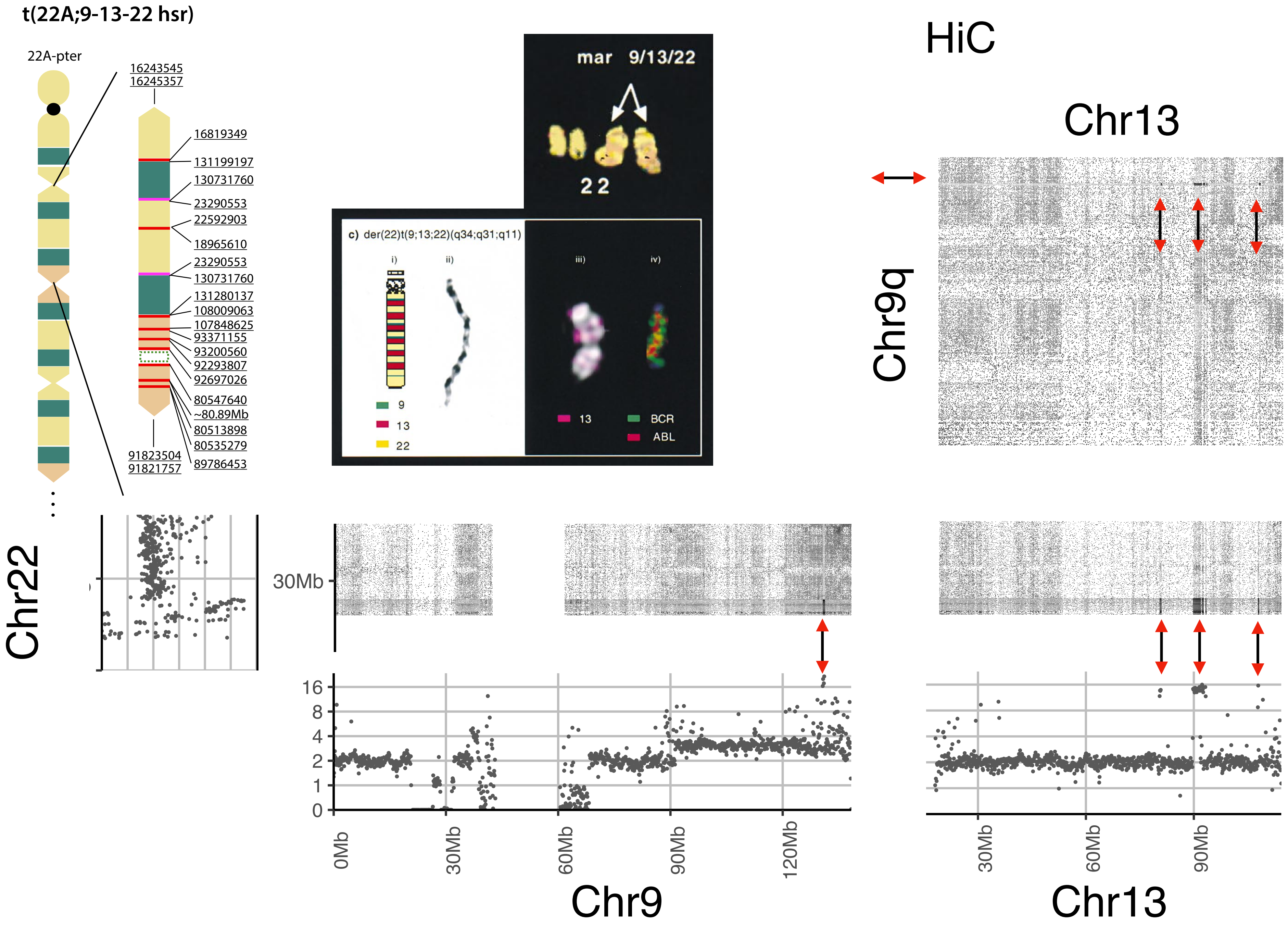
t(22A;9-13-22 hsr)
22A-pter
16243545
16245357
16819349
131199197
130731760
23290553
22592903
18965610
23290553
130731760
131280137
108009063
107848625
93371155
93200560
92293807
92697026
80547640
~80.89Mb
80513898
80535279
89786453
91823504
91821757
mar 9/13/22
22
c) der(22)t(9;13;22)(q34;q31;q11)
i)
ii)
iii)
iv)
9
13
22
13
BCR
ABL
HiC
Chr13
Chr9q
Chr22
30Mb
16
8
4
2
1
0
0Mb
30Mb
60Mb
90Mb
120Mb
Chr9
30Mb
60Mb
90Mb
Chr13

#### t(22A;9-13-22 hsr)

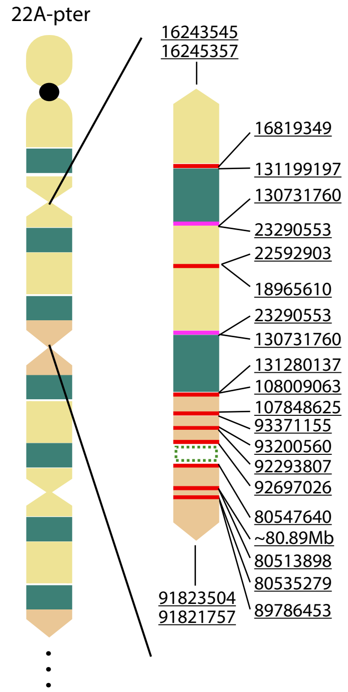


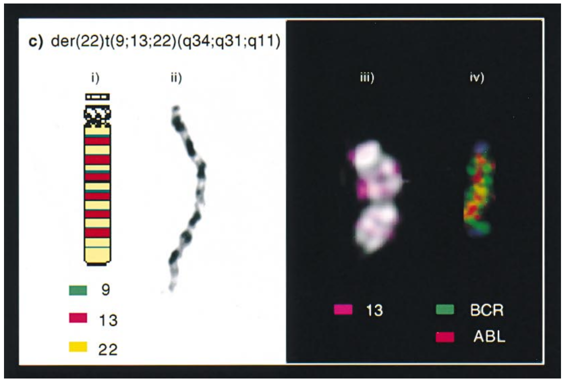


# 10X

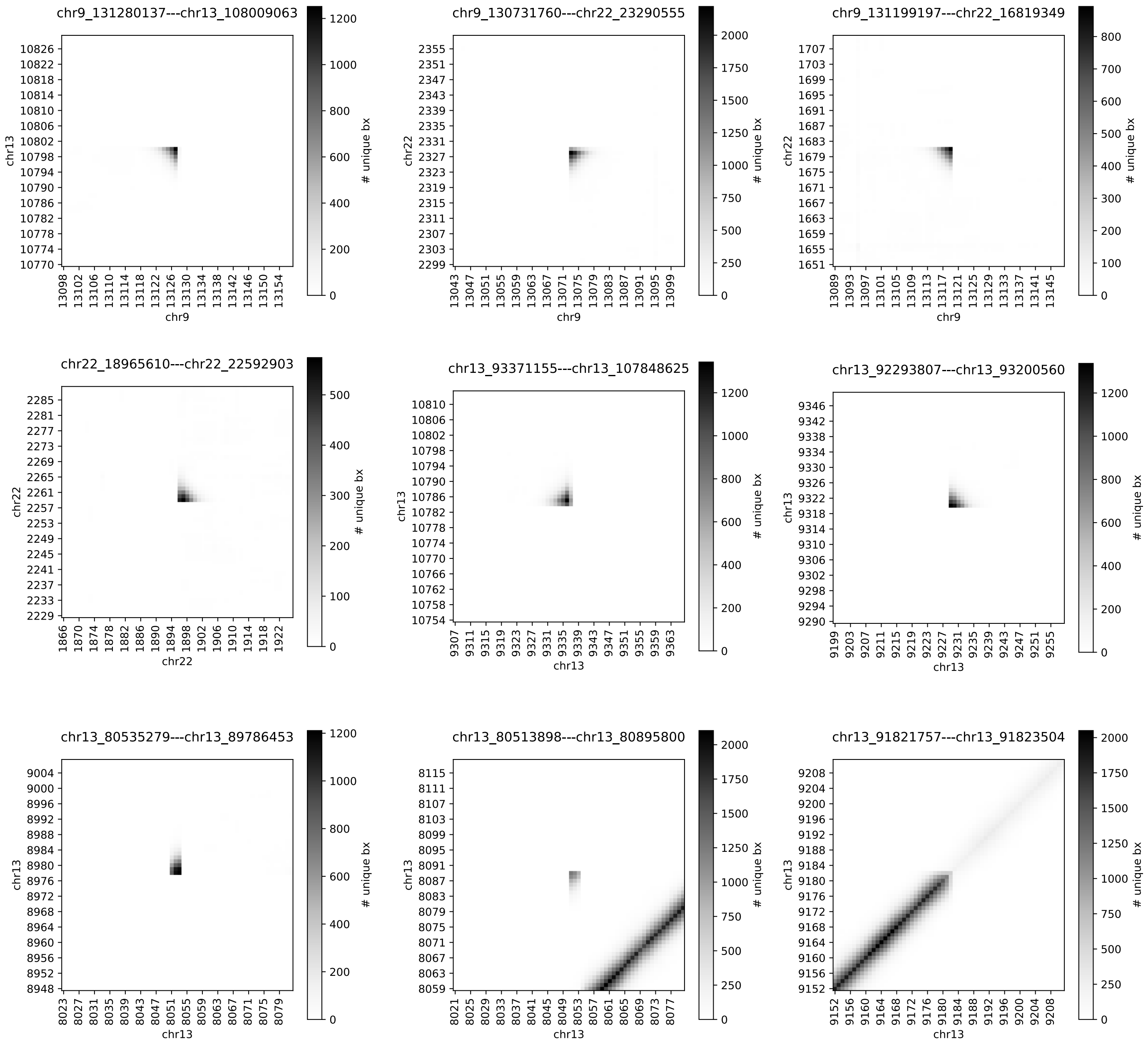
